# Generalizable neuromarker for autism spectrum disorder across imaging sites and developmental stages: A multi-site study

**DOI:** 10.1101/2023.03.26.534053

**Authors:** Takashi Itahashi, Ayumu Yamashita, Yuji Takahara, Noriaki Yahata, Yuta Y. Aoki, Junya Fujino, Yujiro Yoshihara, Motoaki Nakamura, Ryuta Aoki, Haruhisa Ohta, Yuki Sakai, Masahiro Takamura, Naho Ichikawa, Go Okada, Naohiro Okada, Kiyoto Kasai, Saori C. Tanaka, Hiroshi Imamizu, Nobumasa Kato, Yasumasa Okamoto, Hidehiko Takahashi, Mitsuo Kawato, Okito Yamashita, Ryu-ichiro Hashimoto

## Abstract

Autism spectrum disorder (ASD) is a lifelong condition, and its underlying biological mechanisms remain elusive. The complexity of various factors, including inter-site and development-related differences, makes it challenging to develop generalizable neuroimaging-based biomarkers for ASD. This study used a large-scale, multi-site dataset of 730 Japanese adults to develop a generalizable neuromarker for ASD across independent sites (U.S., Belgium, and Japan) and different developmental stages (children and adolescents). Our adult ASD neuromarker achieved successful generalization for the US and Belgium adults (area under the curve [AUC] = 0.70) and Japanese adults (AUC = 0.81). The neuromarker demonstrated significant generalization for children (AUC = 0.66) and adolescents (AUC = 0.71; all *P*<0.05, family-wise-error corrected). We identified 141 functional connections (FCs) important for discriminating individuals with ASD from TDCs. These FCs largely centered on social brain regions such as the amygdala, hippocampus, dorsomedial and ventromedial prefrontal cortices, and temporal cortices. Finally, we mapped schizophrenia (SCZ) and major depressive disorder (MDD) onto the biological axis defined by the neuromarker and explored the biological continuity of ASD with SCZ and MDD. We observed that SCZ, but not MDD, was located proximate to ASD on the biological dimension defined by the ASD neuromarker. The successful generalization in multifarious datasets and the observed relations of ASD with SCZ on the biological dimensions provide new insights for a deeper understanding of ASD.

## INTRODUCTION

Establishing robust biomarkers for autism spectrum disorder (ASD) is essential for understanding the pathophysiological mechanisms of this disorder and for early diagnosis and appropriate interventions. A plethora of modalities, including genetics, electrophysiology, and neuroimaging, are employed to identify biomarkers for ASD *(1)*. Yet, no reliable biomarker has been established *(2)* because of the complicated relationships of various factors, such as genetic and environmental factors *(3, 4)*, biological sex *(5)*, cultural factors *(6)*, and developmental factors *(7, 8)*, all of which form the heterogeneity of ASD. Neuroimaging-based biomarkers, here we call neuromarkers, hold promise in their potential to achieve greater classification accuracy with fewer participants than genetic biomarkers, which only explain 2.45% of risk variance even with more than 10,000 cases *(9, 10)*. However, there are several challenges to the development of robust neuromarkers.

One of the challenges is the requirement for a large-scale dataset that often exceeds the capacity of a single institution. Multi-site initiatives, such as the Autism Brain Imaging Data Exchange (ABIDE) *(11, 12)*, have enabled researchers to leverage neuroimaging and machine learning techniques to develop neuromarkers *(13, 14)*. The classification performance of current neuromarkers ranges from 60% to 90%, depending on the sample size, cross-validation scheme, and brain features utilized in the classification *(15)*. This multi-site, large-scale data-sharing framework has greatly facilitated the development of neuromarkers for ASD. A recent meta-analytic study, however, has reported that 93% of classification studies rely on the ABIDE data *(16)*, potentially introducing biases associated with repeatedly using the same dataset *(9, 17)*. It is, thus, crucial to validate these neuromarkers with completely independent datasets to ensure their generalizability and robustness.

Another challenge is the inter-site variability caused by differences in the scanning protocols and MRI scanners. The diagnostic effect is smaller than the inter-site difference inherent in brain features *(18)*, which makes it challenging to capture the reliable diagnostic effect in multi-site neuroimaging data. Various methods have been proposed to correct inter-site differences *(18, 19)*; prior multi-site, case-control studies have confirmed the effectiveness of these methods *(20, 21)*. Some classification studies have employed an inter-site cross-validation scheme to evaluate the generalizability of their classifiers to other imaging sites within a dataset *(22–24)*. However, this approach may not be enough as an index for generalizability, as it only repeats the analyses on the same dataset. Only a few studies have attempted to construct ASD neuromarkers and explicitly test them on completely independent datasets *(9, 25–27)*.

Once a generalizable neuromarker is established, it paves the way towards addressing further core questions on the neural mechanisms of ASD and psychiatric disorders more generally. One application is to examine whether persistent alterations exist in the autistic brain across different developmental stages. As suggested by developmental changes in the severity of clinical symptoms *(7, 8)*, the patterns of structural and functional alterations in the autistic brain may change with the developmental stages *(28–30)*. Prior classification studies have reported that classifiers trained on the adult sample cannot distinguish adolescents with ASD from typically developing adolescents and vice versa *(31, 32)*. On the other hand, recent large-scale, case-control studies have reported reproducible and trait-like alterations of resting-state functional connectivity (FC) in the ASD population *(21, 33)*. These findings suggest that if any development-independent, atypical neural substrate exists throughout the lifespan, then a neuromarker may successfully generalize across developmental stages.

Another possible application is to investigate the biological continuity between ASD and other psychiatric disorders. There has been a growing interest in recent years to clarify the biological continuum between ASD and other psychiatric disorders due to the overlaps in the symptoms and impaired cognitive functions *(34, 35)*. Utilizing neuromarkers as biological axes and projecting categorically-distinct disorders onto the biologically defined space allows for examining cross-disease relationships without collecting additional symptom scales *(25)*. Our prior study has revealed the asymmetric relationships whereby schizophrenia (SCZ) exhibits greater proximity towards ASD than typically developing controls (TDCs) on the ASD neuromarker, while ASD exhibits greater closeness towards TDCs than SCZ on the SCZ neuromarker *(36)*. Despite the known high comorbidity rate of major depressive disorder (MDD) in ASD *(37)*, the relationship between ASD and MDD has yet to be examined on these biological dimensions. Replication of the relationship with SCZ and exploration of the relationship with MDD may disentangle the complicated relationships between ASD and these disorders.

In this study, we tackled the above challenges through the following procedures. First, we built an FC-based classifier using the multi-site, multi-disease dataset for the Japanese adult population, called the Strategic Research Program for the Promotion of Brain Science (SRPBS) dataset *(38)*. We then applied this adult ASD classifier to the U.S., Belgium, and Japanese adult validation datasets to evaluate its generalization performance to independent imaging sites. Generalization performance was further tested across different developmental stages by applying our adult ASD classifier to child and adolescent validation datasets. These analyses identified a set of FCs associated with ASD status that are invariant across different developmental stages. Finally, we examined the relationships between ASD and two major psychiatric disorders (i.e., SCZ and MDD) in the dimensional space defined by the neuromarkers.

The strengths of our framework are multifaceted. Firstly, we have evaluated the generalization performance of our ASD classifier on different types of validation datasets representing diverse ethnic and cultural backgrounds and developmental stages. This methodology allows for the assessment of generalization performance across independent imaging sites, geographical and cultural variations and different developmental stages.Secondly, we identified vital FCs in a data-driven manner. This approach can eliminate unwanted bias in identifying the neural substrate of ASD. Indeed, this procedure has identified FCs with a consistent diagnostic effect across datasets, potentially representing the core neural substrate of ASD. Lastly, the classifier was developed based on a dataset from the adult population. The utilization of adult data, in contrast to data from children, enables us to examine the commonalities and distinctions between ASD and major psychiatric disorders that manifest post-adolescence. This is of crucial importance for a deeper understanding of the relationship between ASD and other psychiatric disorders.

## RESULTS

### Japanese-adult-based neuromarker for ASD generalized to two adult validation datasets

The neuromarker built on the discovery dataset (550 TDC adults and 180 adults with ASD; see table S1 for participant characteristics) with 71,631 FCs discriminated the ASD group from the TDC group with an accuracy of 76%, AUC of 0.84, and MCC of 0.46. The corresponding sensitivity and specificity were 76% and 75%, respectively (Fig. 1A; see fig. S1A and table S2 for the classification performance in each imaging site). These results demonstrate the acceptable discrimination ability of our ASD classifier on the discovery dataset.

**Fig. 1.**
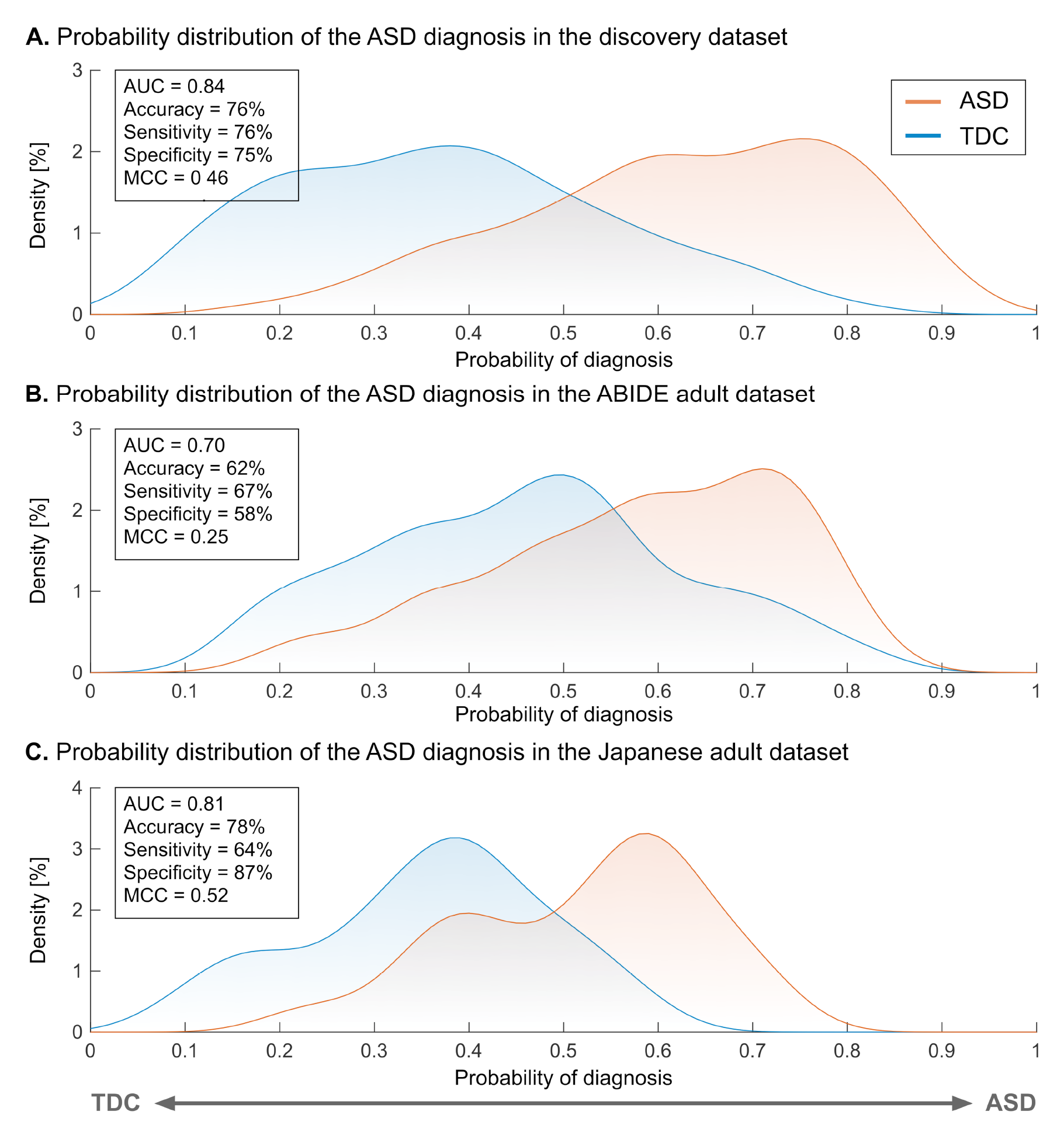
The classification performance of the ASD classifier in the discovery and adult validation datasets. (A) The probability of the ASD diagnosis in the discovery dataset. The probability of the ASD diagnosis in the ABIDE adult validation dataset. (C) The probability of the ASD diagnosis in the Japanese adult validation dataset. **Abbreviations:** AUC: area under the curve, ASD: autism spectrum disorder, MCC: Matthews correlation coefficient, and TDC: typically developing control.

We tested the generalizability of our neuromarker on the two independent adult validation datasets (see table S1 for participant characteristics). For the ABIDE adult dataset, our neuromarker showed an accuracy of 62% (Fig. 1B). The corresponding AUC and MCC were 0.70 and 0.25 (all *P* < 0.05, family-wise error (FWE) corrected). The sensitivity and specificity were 67% and 58%, respectively (see fig. S1B and table S2 for the classification performance in each imaging site). These results indicate that our marker has significant discrimination ability in the ABIDE adult dataset. For the newly collected Japanese adult dataset, our neuromarker exhibited an accuracy of 78% (Fig. 1C and table S2). The corresponding AUC and MCC were 0.81 and 0.52, respectively (all *P* < 0.05, FWE-corrected). The sensitivity and specificity of this marker were 64% and 87%, respectively. These results suggest that our neuromarker for the ASD diagnosis generalizes to data from independent imaging sites in the same developmental stage.

### Adult ASD neuromarker is generalizable to the children and adolescents

We further tested the generalizability of our adult ASD neuromarker to children (< 12 years old) and adolescents (12 < age < 18 years old) datasets (see table S3 for participant characteristics). Our neuromarker showed significant classification performance in the children (accuracy = 61%, AUC = 0.66 and MCC = 0.27; *P* < 0.05, FWE-corrected; Fig. 2 and table S4) and the adolescents (accuracy = 66%, AUC = 0.71, and MCC = 0.32; *P* < 0.05, FWE-corrected). These results suggest that our neuromarker for adult ASD could generalize to other developmental stages even in independent imaging sites.

**Fig. 2.**
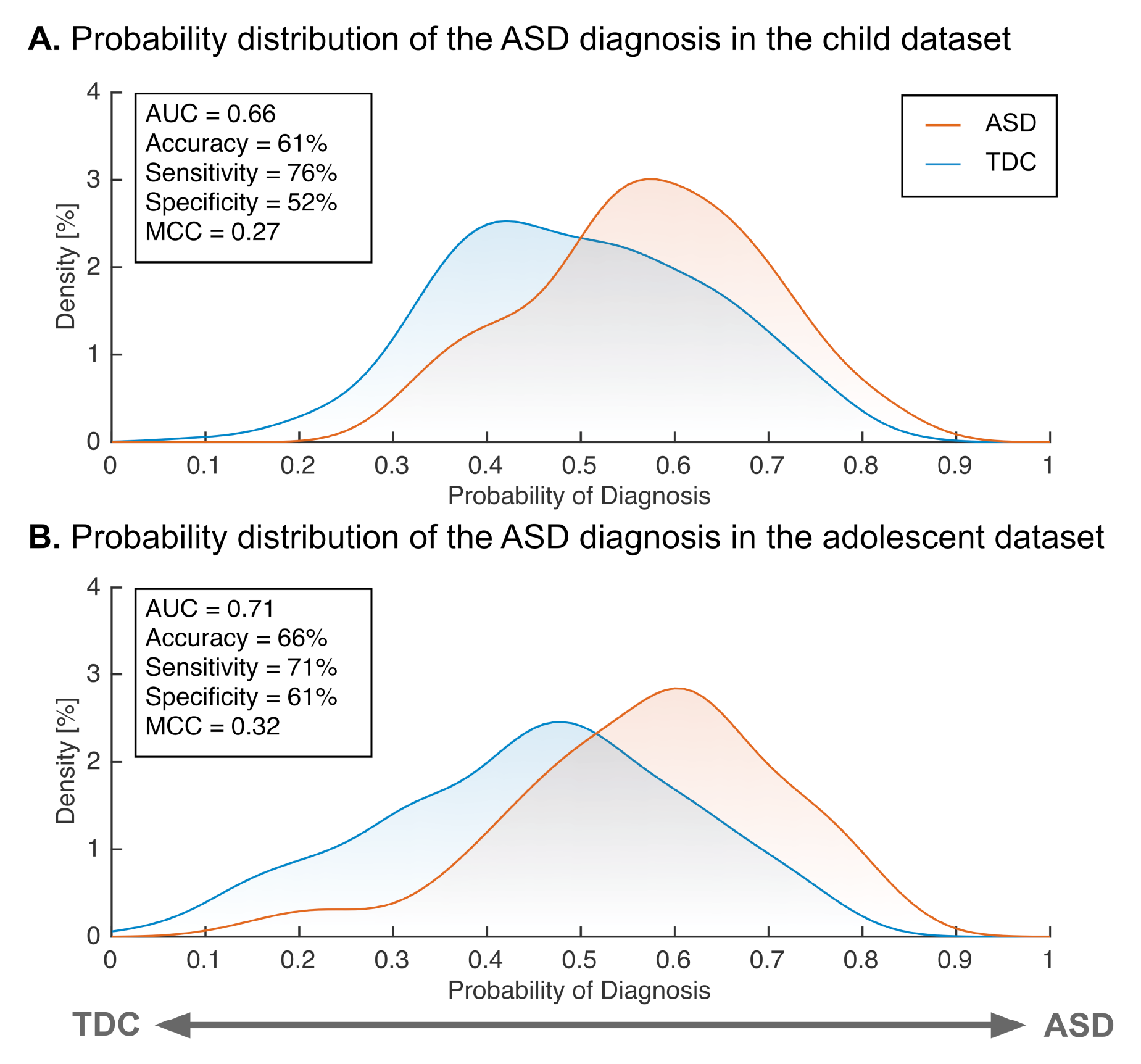
The classification performance of the ASD classifier in the child and adolescent validation datasets. (A) The probability of ASD diagnosis in children. (B) The probability of ASD diagnosis in adolescents. **Abbreviations:** AUC: area under the curve, ASD: autism spectrum disorder, MCC: Matthews correlation coefficient, and TDC: typically developing control.

### Potential factors affecting the generalization performance

We explored the impacts of head motion, harmonization, and other experimental factors (i.e., diversity in imaging sites and the choice of atlas) on the generalization performance. The details were described in **Supplementary Materials**. We first assessed whether the head motion artificially improved the generalization performance. A significant positive correlation was not found between the AUC and mean FD (*r* = −0.56, *P* = 0.002; fig. S2), indicating that head motion did not improve the generalizability of our neuromarker. We next assessed the impacts of the harmonization method on generalization performance. In both validation datasets, we observed degraded generalization performance without the harmonization (ABIDE adults: accuracy = 60% and AUC = 0.64; Japanese adults: accuracy = 67% and AUC = 0.73; table S5), supporting the necessity of the harmonization method for improving the generalization performance. We then examined the impacts of imaging sites on the generalization performance because our discovery dataset comprised some imaging sites having either the imbalanced patient/control ratio (i.e., COI, KUT, and UTO1) or a different scanning protocol (i.e., UTO2). Neither the inclusion of imbalanced imaging sites nor that of a site with a different scanning protocol affects the generalizability of our classifier (tables S6 and S7). Finally, we checked whether our generalization performance was atlas-dependent. We constructed classifiers using the Schaefer cortical atlas *(39)* that provided atlases with multiple levels of resolutions ranging from 100 to 900. We did not observe any improvements specific to the Glasser atlas in the generalization performance (table S8).

The tests of generalizability of our ASD neuromarker to independent validation datasets depend on the assumption that altered FC patterns characteristic to ASD if any, should be reproducible between the discovery dataset and the validation datasets. To directly test this assumption, we used a mass-univariate analysis similar to previous studies *(40)*. Briefly, we first quantified the between-group difference in each FC by calculating the *t*-value (the diagnosis effect) for each dataset. We then computed the Pearson correlation coefficient of *t*-values between the datasets. Statistical significance was tested using a permutation test with 1,000 iterations, and the statistical threshold was set to *P* < 0.05, one-sided. The discovery dataset showed statistically significant positive correlations with other datasets (ABIDE adult: *r* = 0.16; Japanese adult: *r* = 0.31; and adolescent: *r* = 0.20, all *P* < 0.05) except for the child dataset (*r* = 0.02, *P* = 0.34; fig. S3).

### FCs associated with the ASD diagnosis

We identified a set of discriminative FCs associated with the ASD status (see **Methods** for details): 65 hyper-connections and 76 hypo-connections. We called these FCs as “discriminative FCs” hereafter. To examine the spatial distribution of these FCs, we counted the number of occurrences in which each brain region was selected as at least one of the terminations of each discriminative FC. Figures 3A and 3B show the spatial distributions of identified FCs and their terminal regions, and table S9 provides the connection details. In the cerebral cortex, the bilateral temporal cortices and dorsomedial and ventromedial prefrontal cortices (dmPFC and vmPFC) were notably affected. On the other hand, the right amygdala, midbrain, hippocampus, globus pallidum, and putamen were affected among the subcortical regions.

**Fig. 3.**
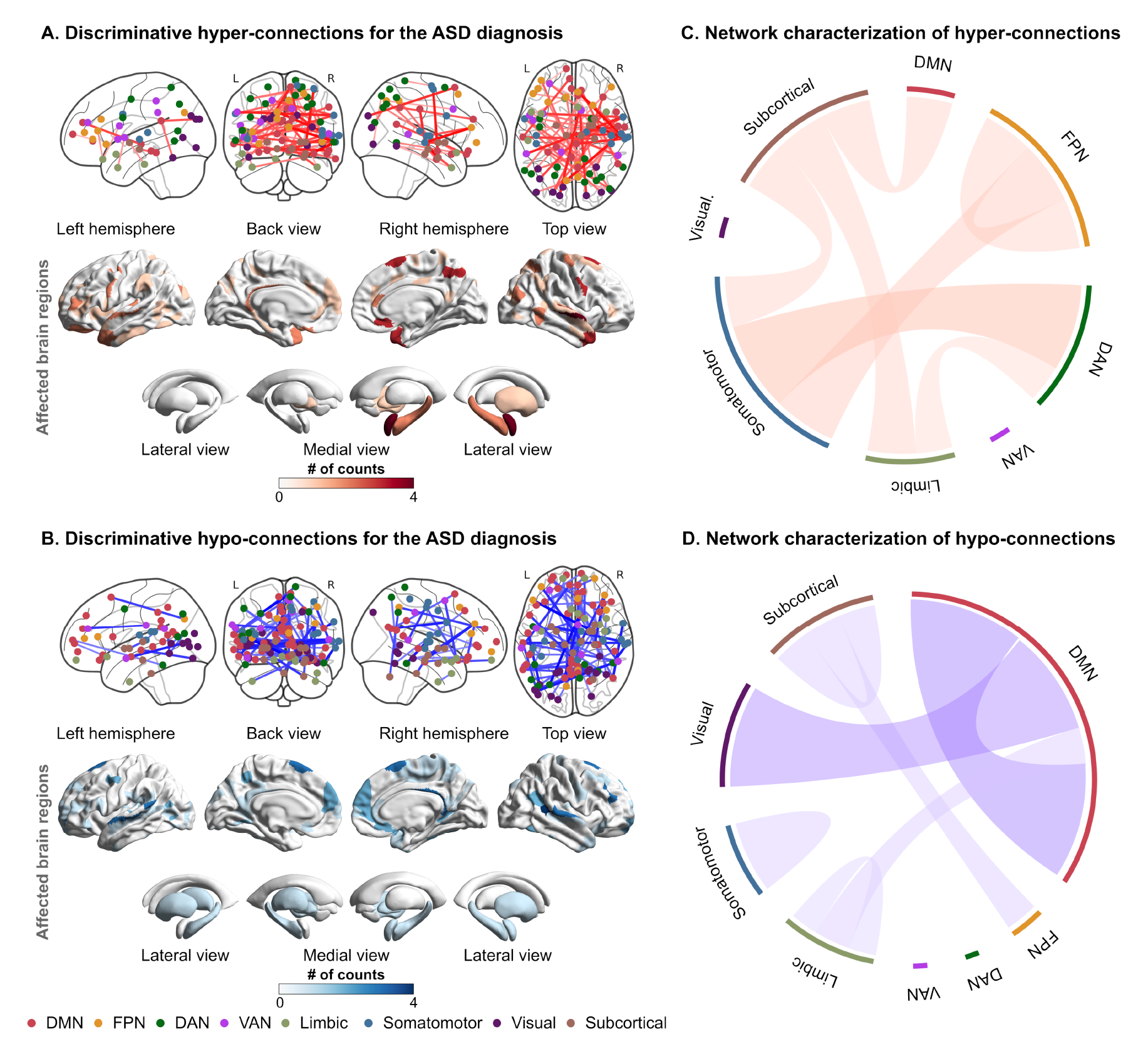
The spatial distribution and network-based characterization of discriminative functional connections for the ASD diagnosis. (A) The spatial distribution of discriminative hyper-connections and affected brain regions for the ASD diagnosis. (B) The spatial distribution of discriminative hypo-connections and affected brain regions for the ASD diagnosis. The node color represents the corresponding resting-state network. (C) Network-based characterization of hyper-connections. (D) Network-based characterization of hypo-connections. **Abbreviations:** ASD: autism spectrum disorder, DAN: dorsal attention network, DMN: default mode network, FPN: frontoparietal network, TDC: typically developing control, and VAN: ventral attention network.

To further functionally characterize these FCs, we delineated each of the hyper-and hypo-connections into functional network anatomy *(41)*. Hyper-connections were dominantly characterized by between-network connections stemming from the subcortical and somatomotor networks to other networks, such as the default mode network (DMN), dorsal attention network (DAN), and frontoparietal network (FPN) (Fig. 3C). On the other hand, hypo-connections were notably characterized by within-network connections of the DMN and between-network connections of the DMN and visual network (Fig. 3D).

### Consistency of discriminative FCs across datasets

Among 141 discriminative FCs identified by permutation tests, we further investigated which discriminative FCs showed consistent alterations across the five datasets. We computed a *t*- value as the effect of diagnosis in each FC of each dataset. We then counted the number of FCs showing the same sign across the five datasets. Forty-two out of 141 discriminative FCs showed the same sign of *t*-values across all the datasets (Hyper-connection: 52.4% and hypo-connection: 47.6%; Fig. 4A and table S10). When considering over-and under-connectivity (i.e., the difference in the absolute of FC strength between groups), 69% of the reproducible FCs were under-connectivity (Fig. 4B). The ASD group exhibited atypical connections stemming from subcortical networks, including the midbrain, hippocampus, and amygdala, to other networks, such as the somatomotor network and FPN (Figs. 4C and 4D).

**Fig. 4.**
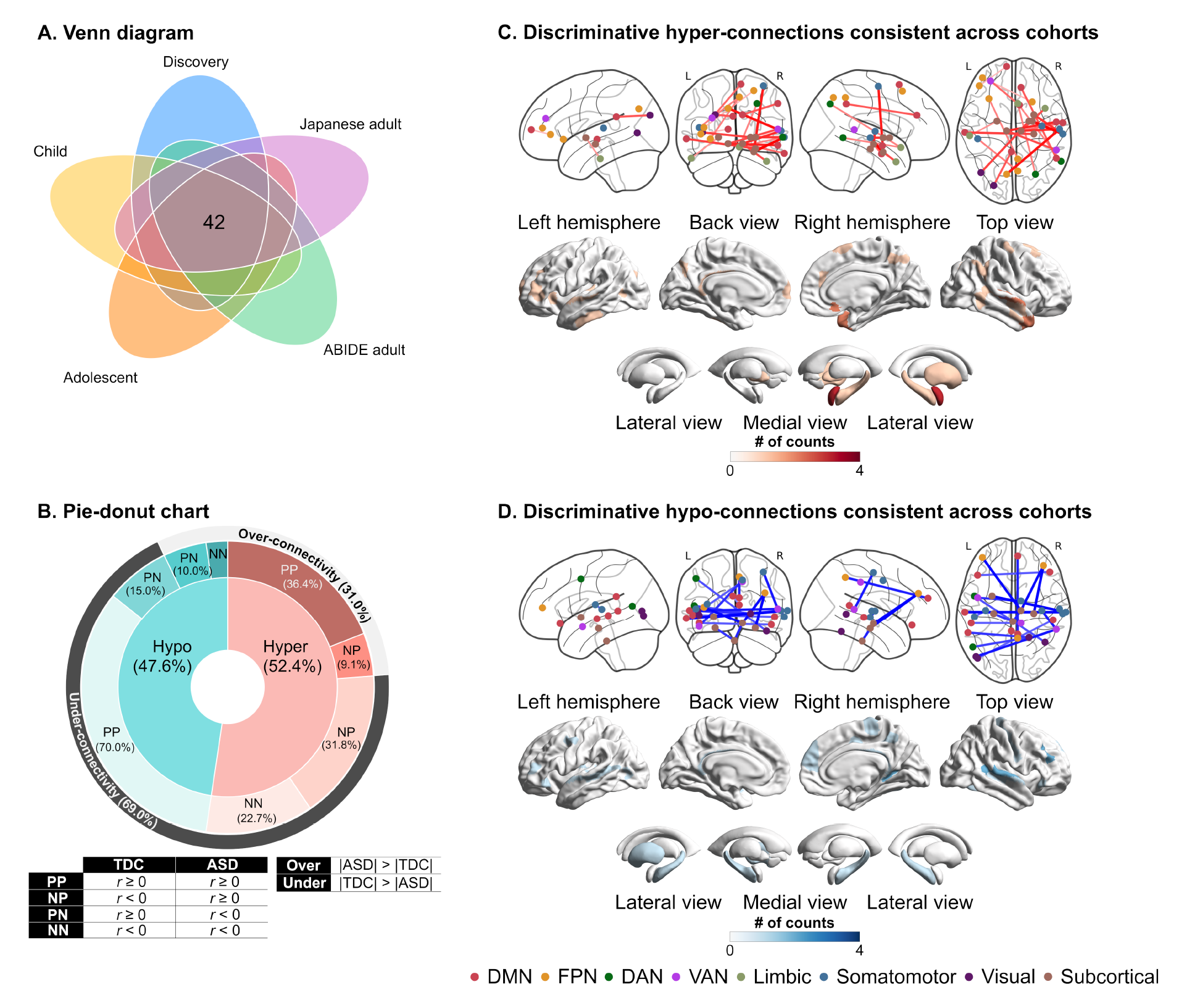
The distribution of atypical functional connections that are consistent across the datasets. (A) The spatial distribution of discriminative hyper-connections consistent across cohorts. (B) The spatial distribution of discriminative hypo-connections consistent across cohorts. (C) Pie-donut chart showing the details of reproducible hypo-and hyper-connections. **Abbreviations:** ABIDE: autism brain imaging data exchange, DAN: dorsal attention network, DMN: default mode network, FPN: frontoparietal network, L: left, N: Negative, P: positive, R: right, and VAN: ventral attention network.

The statistical significance of the consistency among the five datasets was examined using a binomial test *(42)*. The number of consistent FCs within the discriminative FCs was 42, while the number of FCs showing the same sign of diagnostic effects in the whole FCs (i.e., 71,631 FCs) was 9,547. We assumed a binomial distribution, *Bi*(*n*, *p*), where *n* stands for the number of discriminative FCs (i.e., *n* = 141), and *p* stands for the probability of being consistent across the datasets in the whole FCs (i.e., *p* = 9,547/71,631). The binomial test confirmed that discriminative FCs were reproducible across different ethnicities and cultures (i.e., U.S., Belgium, and Japan) and different developmental stages (i.e., child, adolescent, and adult) (*P* < 0.05). The binomial tests confirmed the statistical significance of the consistency between the discovery dataset and every dataset (see **Supplementary Materials** for details). These results indicate that the selection of discriminative FCs showing consistent alterations may yield the generalizability of our neuromarker across independent imaging sites and developmental stages.

### Specificity of the ASD neuromarker

We applied our ASD neuromarker to SCZ and MDD to test the specificity of the neuromarker (see table S11 for participant characteristics). Our ASD neuromarker exhibited 56% and 33% sensitivities for SCZ and MDD, respectively (Fig. 5A). As expected, the sensitivities for both disorders were greatly decreased compared with that for the ASD diagnosis. However, the sensitivity to the SCZ diagnosis was still statistically significant (*P* = 0.014), whereas the marker showed no sensitivity to the MDD diagnosis (*P* = 0.638). These results indicate that SCZ, but not MDD, was located proximate to ASD on the biological dimension defined by the ASD neuromarker.

**Fig. 5.**
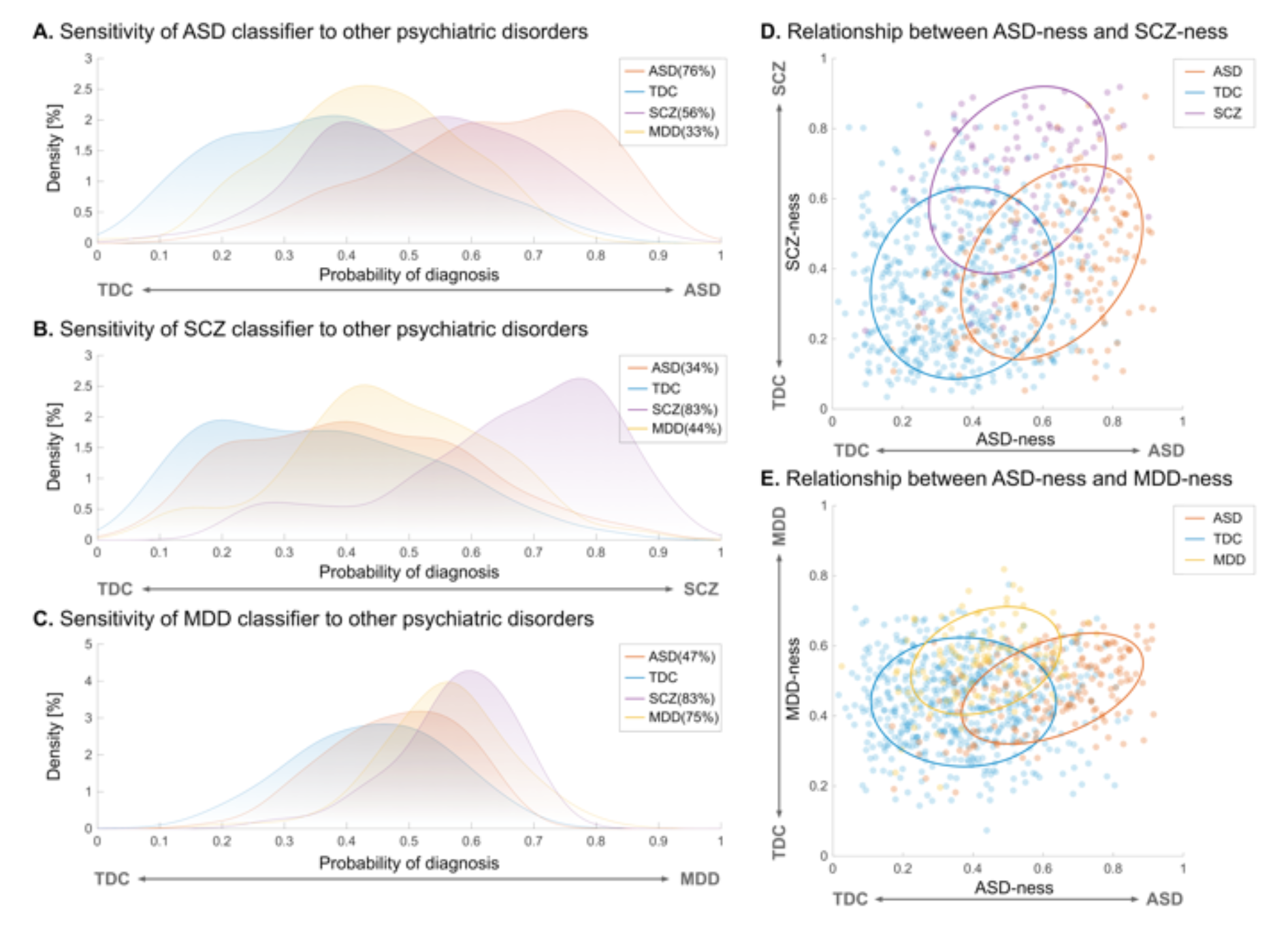
Sensitivity of each neuromarker to other psychiatric disorders and dimensional relationships between ASD and other psychiatric disorders. (A) Sensitivity of the neuromarker for autism spectrum disorder (ASD) to schizophrenia (SCZ) and major depressive disorder (MDD). (B) Sensitivity of the SCZ neuromarker to ASD and MDD. (C) Sensitivity of the MDD neuromarker to ASD and SCZ. (D) Dimensional relationship between the ASD-ness and SCZ-ness defined by the diagnostic probability in ASD and SCZ classifiers, respectively. (E) Dimensional relationship between the ASD-ness and MDD-ness defined by the diagnostic probability in ASD and MDD neuromarkers, respectively.

### Relationships between ASD and other psychiatric disorders on biological dimensions

To further explore the relationships of the three disorders on multiple biological dimensions, we built two additional classifiers for the SCZ and MDD (see table S12 for the classification performance of both classifiers). We then applied each of them to the other two disorders (see **Supplementary Materials** for details). The SCZ classifier exhibited poor sensitivity to the ASD and MDD diagnoses (ASD = 34% and MDD = 44%, all *P* > 0.39; Fig. 5B). On the other hand, the MDD classifier showed poor sensitivity to the ASD diagnosis (sensitivity = 47%) but high sensitivity to the SCZ diagnosis (sensitivity = 83%; Fig. 5C). We next plotted each participant onto the two-dimensional planes defined by the neuromarkers to illustrate the relationship between ASD and other psychiatric disorders. On the two-dimensional plane characterized by the ASD and SCZ neuromarkers, ASD predominantly resided in the lower right quadrant, while SCZ was primarily situated in the upper right quadrant (Fig. 5D), suggesting an asymmetric proximity between ASD and SCZ. In contrast, on the two-dimensional space demarcated by the ASD and MDD neuromarkers, ASD was mainly distributed in the lower right quadrant, while MDD was mainly localized in the upper left quadrant (Fig. 5E). These results suggest that, although SCZ shares characteristics with ASD, ASD and MDD appear to be distinct entities on the biological dimensions.

We further investigated why ASD and SCZ neuromarkers showed an asymmetric proximity by first checking the spatial overlap of discriminative FCs between the two neuromarkers. Permutation tests with 100 iterations identified discriminative FCs for SCZ (fig. S4 and table S13; see fig S5 and table S14 for the discriminative FCs for MDD). When comparing them with those for ASD, we observed no spatial overlap between the two neuromarkers. Next, we examined the similarities of the diagnosis effects between the two groups. We expected that the effects of diagnoses were similar in the ASD neuromarker but not in the SCZ neuromarker. We observed significant positive correlations in the ASD neuromarker (hyper-connection: *r* = 0.63, *P* < 0.05; hypo-connection: *r* = 0.60, *P* < 0.05) but not in the SCZ marker (hyper-connection: *r* = 0.05, *P* > 0.86; hypo-connection: *r* = −0.44, *P* < 0.05; fig. S6). These results indicate that the asymmetric diagnostic effects on both neuromarkers yield such relationships between ASD and SCZ on the biological dimensions.

## DISCUSSION

In this study, we developed an FC-based neuromarker for ASD diagnosis using the Japanese adult multi-site dataset. Our ASD neuromarker was generalizable with >70% AUC value to both the ABIDE and Japanese adults from independent imaging sites. This neuromarker also exhibited acceptable generalization performance to children and adolescents, suggesting the presence of atypical neural basis that persists in ASD from childhood to adulthood. We identified 141 FCs that were pivotal in determining the ASD status. These FCs spanned multiple networks, including the DMN, somatomotor, and subcortical networks, and comprised social brain regions, such as the vmPFC, dmPFC, STG, right amygdala, midbrain, and hippocampus. Forty-two out of 141 FCs displayed consistent effects of diagnostic status across all the five datasets. By utilizing our neuromarker as a biological axis and mapping SCZ and MDD onto the axis, we replicated previous findings suggesting proximity between ASD and SCZ and a greater distance between ASD and MDD. Our neuromarker-based analytical framework offers novel insights into the neural underpinnings of the autistic brain and provides an effective tool for elucidating transdiagnostic continuity.

We confirmed the validity of our neuromarker through stringent evaluations of its generalization performance on independent validation datasets. Most prior studies have relied on cross-validation procedures within the ABIDE dataset *(15, 16)*. Reliance on these methods poses the risk of overestimating the classification performance and introducing biases into the results *(9, 17)*. It is, therefore, critical to evaluate the generalization performance of a neuromarker using a completely independent external dataset *(43)*. A handful of studies have evaluated the external validity of their neuromarkers using independent validation datasets and have reported generalization performance ranging from 67% to 75% *(9, 25, 26)*, comparable to our results. The forte of our study is the thorough evaluation of generalization performance. Through our assessment, we verified the generalizability of our neuromarker utilizing four independent validation datasets that varied in terms of imaging sites, ethnic and cultural backgrounds, and developmental stages. Additionally, we confirmed that the generalizability of our neuromarker remained intact when utilizing another brain atlas featuring multiple resolution levels (table S8). These thorough evaluations ensure the robustness of our neuromarker and provide a new tool for further exploring several hypotheses in the autistic brain.

Developmental-dependent changes in FCs have been a major bottleneck for transferring neuromarkers to other developmental stages *(31, 32)*. Nevertheless, our adult ASD neuromarker successfully distinguished children and adolescents with ASD from TDCs. The successful generalization may be due to the LASSO method used in this study. The autistic brain likely presents a mosaic pattern of age-specific and age-unspecific atypical FCs *(44–46)*. To support this view, Alaerts and colleagues have reported that, depending on the terminal regions, the posterior superior temporal sulcus exhibits distinct age-related patterns of abnormal FCs in the autistic brain *(29)*. In such a complicated situation, non-sparse machine learning algorithms such as support vector machines may have failed to select features appropriately, yielding reduced generalization performance when transferring neuromarkers to other developmental stages. The LASSO method tends to choose atypical FCs consistent within the discovery dataset during nested cross-validation. As a result, dataset-independent atypical FCs may have been selected in our adult ASD neuromarker.

Our ASD neuromarker identified 141 discriminative FCs distributed across multiple networks, most notably the DMN, somatomotor, and subcortical networks. One of the most prominent features is hypo-connectivity within the DMN, including the dmPFC, vmPFC, and temporal cortices. The hypo-connectivity within the DMN and its association with socio-communicative deficits in ASD have been repeatedly reported *(47–49)*. While, in this study, discriminative FCs stemming from temporal cortices exhibited the effect of ASD status consistently across the five datasets, those stemming from dmPFC and vmPFC did not. Given that population heterogeneity is associated with the DMN, especially dmPFC, vmPFC, and precuneus *(50)*, the dataset-dependent effect of ASD status on these discriminative FCs across the datasets may reflect differences in the population diversity.

Another prominent feature is alterations in FCs within and between subcortical and somatomotor networks, encompassing the amygdala, hippocampus, midbrain, and STG. In addition, we observed that 26 of 42 discriminative FCs that showed a consistent effect of ASD status across the five datasets belonged to these networks. Our findings imply that abnormalities in these networks may serve as the core neural basis for this disorder. To support this possibility, several lines of evidence suggest functional alterations within and between subcortical and somatomotor networks *(11, 45, 46)* and their associations with the clinical symptoms, such as restricted and repetitive behaviors and atypical sensory processing *(21, 41)*. Furthermore, the association between atypical sensory processing and socio-communicative impairments is supported by previous literature *(51, 52)*. Replicability, associations with core symptoms, and consistency of abnormalities through developmental stages may distinguish these networks as a core basis for ASD.

We demonstrated the utility of our generalizable neuromarker-based analytic framework for examining the biological continuum of ASD with SCZ and MDD by mapping these disorders onto the biological dimensions defined by the neuromarkers. Previous studies have highlighted the substantial overlap between ASD and SCZ at genetic *(53)*, neural *(54)*, and behavioral levels *(55)*. However, these findings are less replicable due to methodological differences *(56)*. In contrast, we reproduced the asymmetric proximity between the ASD and SCZ group, whereby the SCZ group manifested stronger adjacency to the ASD group than the TDC group on the ASD neuromarker, whereas the ASD group displayed stronger proximity to the TDC group than the SCZ group on the SCZ neuromarker, by using different datasets and methods *(36)*. Our replicable approach may provide a clue to investigating the spectrum structure among psychiatric disorders.

Despite the asymmetric proximity between the ASD and SCZ groups, there was no spatial overlap in discriminative FCs between the ASD and SCZ neuromarkers. One possibility is that the SCZ neuromarker in this study was built on patients with chronic SCZ. Previous studies have reported that abnormal FCs in SCZ tend to change with the disease progression *(36, 57)*. Owing to this change of abnormal FCs with disease progression, patients with SCZ may show abnormalities in discriminative FCs for the ASD neuromarker, which may result in asymmetric proximity between ASD and SCZ. Indeed, we observed that ASD and SCZ status had similar diagnosis effects on discriminative FCs for the ASD neuromarker but not on those for the SCZ neuromarker (fig. S6). The inclusion of patients with SCZ from different disease stages may provide a clue for how this asymmetricity may arise.

In addition to the relationship between ASD and SCZ, we examined the relationship between ASD and MDD, but no asymmetric proximity was observed between the two disorders. Although the comorbidity rate of MDD in ASD is known to be as high as 11% *(37)*, there are no case-control studies to date that examine FCs among ASD, MDD, and ASD comorbid with MDD. A few studies have examined the relationship between atypical FCs and depressive symptoms in ASD *(58, 59)*. For example, Kleinhans and colleagues have reported that social impairments and depressive symptoms in ASD are separately associated with abnormal amygdala circuits, and the patterns of these associations in the autistic brain differ from those in TDCs *(59)*. Our findings may reflect that the diagnoses of ASD and MDD affect different neural circuits. The relationship between ASD and MDD on the biological dimensions could be further clarified by studies that were designed to include ASD cases comorbid with MDD.

In conclusion, we developed an ASD neuromarker generalizable to diverse validation datasets from independent imaging sites and different developmental stages. We further demonstrated the applicability of our generalizable neuromarker to examine the effects of developmental stages on the autistic brain and the biological continuum between ASD and other disorders. Using discriminative FCs identified by our ASD neuromarker, prospective future directions will be opened, including identifying biological subtypes within ASD *(60)* and interventions based on those FCs as targets *(61)*.

## MATERIALS AND METHODS

### Study design

The aim of this study was twofold: the first was to construct a generalizable neuromarker for distinguishing adults with ASD from TDCs across diverse imaging sites, and the second was to demonstrate the utility of this neuromarker by examining its generalizability to children and adolescents and investigating the transdiagnostic continuity of ASD with SCZ, and MDD. This study used three adult resting-state fMRI (R-fMRI) datasets for the analyses: one was used as the discovery dataset, and the remaining two were used as independent validation datasets. Tables S1, S15, and S16 show the demographic information and scanning parameters for the three datasets. Data were sourced from three multi-site initiatives: the SRPBS *(38)*, ABIDE *(11, 12)*, and Brain/MINDS Beyond (BMB) projects *(62)*. All the participants (if appropriate) and their respective parent/legal guardian provided written informed consent. See **Supplementary Materials** for the detailed descriptions of datasets and exclusion criteria.

### Participants

The discovery dataset contained data of 550 TDCs from 5 scanners at 4 imaging sites (University of Tokyo [UTO1 and UTO2], Kyoto University [KUT], Center for Innovation in Hiroshima University (COI), and Showa University [SWA1]) and 180 adults with ASD from 2 sites (SWA1 and UTO2). The first validation dataset (ABIDE adult dataset) consisted of participants from the ABIDE-I *(11)* and −II *(12)*. The ABIDE adult dataset comprised 54 adults with ASD and 67 TDCs that were selected from the following 3 sites. The second validation dataset (i.e., the Japanese adult dataset) consisted of 22 adults with ASD and 38 controls at Showa University (SWA2). The data were collected under the BMB project. The project details were described elsewhere *(62)*. See **Supplementary Materials** for more details of participants characteristics.

### R-fMRI data preprocessing and network construction

We preprocessed all the R-fMRI data using fMRIPrep version 1.1.8 *(63)* and the ciftify toolbox version 2.1.1 *(64)*. See **Supplementary Materials** for the details of preprocessing steps. We performed nuisance regression to remove the effects of artifactual and non-neural sources. Nuisance regressors consisted of six head-motion parameters, averaged signals from subject-specific white matter and cerebrospinal fluid masks, global signal, their temporal derivatives, and linear detrending. After nuisance regression, we applied a band-pass filter (0.008–0.1 Hz) to the residuals. We used frame-wise displacement (FD) as a measure for detecting occasional head movement, and removed volumes with FD > 0.5 mm, as proposed in a previous study *(65)*. We used Glasser’s 379 surface-based brain parcellations (cortical 360 parcellations and subcortical 19 parcellations) as ROIs *(66)*. We computed the temporal correlations of signals among all possible pairs of ROIs and applied Fisher’s *r*-to-*z* transformation, resulting in 71,631 unique FCs for each participant. Because the label of each ROI in Glasser’s atlas was not intuitive, we utilized Yeo’s resting-state network (RSN) labels *(67)* to assign important ROIs to the corresponding RSN label. This study added the subcortical network label to the subcortical and cerebellar regions.

### Controlling for imaging site differences

We used the ComBat harmonization method *(68)* to control the imaging site differences inherently captured in FCs. Because of its simplicity and effectiveness, this method has been used for controlling the site differences in neuroimaging data *(69–71)*. In the current study, we incorporated disease status (0 = TDC, and 1 = ASD), age, and sex as covariates of interest into the ComBat model. Of note, we separately applied the ComBat harmonization method to both discovery and independent validation datasets.

### Construction of the ASD neuromarker using the discovery dataset

Based on previous studies *(25, 36, 40, 72–74)*, we assumed that psychiatric disorder factors were associated with the limited number of FCs, rather than the whole-brain connections. We used a logistic regression analysis with a least absolute shrinkage and selection operator (LASSO) method *(75)* that selects an optimal subset of FCs from the whole brain connections. See **Supplementary Materials** for the construction of our ASD neuromarker. We developed the ASD neuromarker using the LASSO method with 10-fold nested cross-validation (CV) and 10 subsampling, yielding 100 trained classifiers (fig. S7). The mean classifier output value was considered as diagnostic probability, indicating a likelihood of a participant belonging to the ASD class. We considered participants as those with ASD if their diagnostic probability values were higher than 0.5. We calculated the area under the curve (AUC) to assess the classification performance. We also computed accuracy, sensitivity, specificity, and the Matthews correlation coefficient (MCC). The MCC is suitable for the imbalanced dataset because this metric takes into account the ratio of the confusion matrix size *(76)*. We used AUC and MCC as performance indices throughout the paper.

### Generalizability of the ASD neuromarker to the ABIDE and Japanese adult datasets

We tested the generalizability of the ASD neuromarker using two independent validation datasets: the ABIDE adult and the Japanese adult datasets. We applied all the trained classifiers to the independent validation datasets and obtained 100 diagnostic probabilities for each participant. We averaged these diagnostic probabilities for each participant and considered a participant to be a participant with ASD if the mean diagnostic probability was > 0.5. To evaluate the generalization performance, we computed the following performance metrics: AUC, accuracy, sensitivity, specificity, and MCC.

### Generalizability of the ASD neuromarker to the child and adolescent datasets

We further tested the generalizability of the adult ASD neuromarker to children (< 12 years old) and adolescents (12 < age < 18). We used these age ranges according to *(33)*. We created the child and adolescent datasets using the same exclusion criteria as the ones applied to the adult dataset. The child dataset consists of 119 children with ASD and 202 those with TDC from 5 sites. The adolescent dataset consists of 121 adolescents with ASD and 132 TDCs from 8 sites. The details of demographic information and scanning parameters are in tables S3 and S16. We applied the same preprocessing pipeline and harmonization method to both datasets.

## Statistical analyses

### Evaluation of the statistical significance of neuromarker’s generalizability

The statistical significance of AUC and MCC was tested using a permutation test with 500 iterations. At each iteration, we created a permuted dataset by shuffling the diagnostic labels. We then built a classifier for the permuted dataset in the same way as for the non-permuted dataset using a 10-fold nested CV with 10 subsamples, which resulted in 100 permuted classifiers at each iteration. Each individual of the independent datasets was classified based on 100 diagnostic probability values, each of which was generated by 100 trained permuted classifiers. The threshold was set at the mean diagnostic probability value of 0.5. We calculated the AUC and MCC for a permuted classifier at each iteration. We constructed a null distribution for each performance index by aggregating those values across iterations. The statistical threshold was set to 0.05. The Holm-Bonferroni method *(77)* was used to control the FWE rate across all the validation datasets.

### Identifications of FCs associated with the clinical diagnosis of ASD

We identified discriminative FCs using a permutation test with 500 iterations in a similar manner to *(40)*. First, as to each FC, we counted the number of times the LASSO selected the FC during the 10-fold CV. If a given FC was consistently important for discrimination throughout the training dataset, we expected that it would be selected significantly more times by the LASSO than the chance derived from a null distribution. We shuffled the diagnostic labels at each iteration to create a permuted dataset and constructed permuted classifiers. For each FC, we counted the number when the FC was selected by the LASSO across 10-fold cross-validation and 10 subsampling (i.e., across 100 classifiers). We used the maximum counts among all the connections at each iteration to construct a null distribution using these maxima. We considered FCs as significant contributors to the ASD status if their *P*-values were below 0.05.

### Network-based characterization of discriminative FCs

To interpret the spatial distribution of discriminative FCs at the network level, we used the network anatomy, according to *(41)*. We identified discriminative FCs using the permutation test described above. We computed the mean weights for those FCs. We considered FCs with positive mean weights as hyper-connections and those with negative mean weights as hypo-connections in the ASD brain. It turned out that we found 65 hyper-connections and 76 hypo-connections (see **Results**). Then, we computed the probability of spatial overlap between FCs within or between-RSNs defined by the Yeo atlas and hyper-or hypo-connections identified by our neuromarker. *P*-values were computed using the hypergeometric cumulative density function “*hygecdf*” implemented in MATLAB as follows:

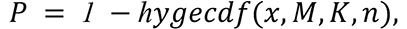

where *x* stands for the number of overlapping FCs between hyper-or hypo-connections and within or between predefined RSNs. The variable, *M*, stands for the number of total FCs in the whole brain (i.e., *M* = 71,631), while the variables, *n* and *K*, were the total numbers of FCs in our network of interest and the canonical network of interest, respectively. We computed the *P*-values for hyper-and hypo-connections separately. Statistical significance was set to *P* < 0.05, adjusted with Bonferroni correction for multiple comparisons.

### Sensitivity of the ASD neuromarker to other psychiatric disorders

We tested whether the developed ASD neuromarker was sensitive to SCZ and MDD. To test this, we applied the neuromarker to data from SCZ and MDD included in the SRPBS dataset *(38)*. The details of their demographic information are in table S11. We applied the same preprocessing pipeline, the exclusion criteria, and the harmonization method.

We applied 100 trained ASD classifiers to SCZ and MDD datasets and obtained 100 diagnostic probability values for each participant. We considered a participant to have the ASD label if the mean diagnostic probability was > 0.5. Since participants with TDC were identical among the datasets, we used the diagnostic probability values during the training to compute the classification performance. We focused on sensitivity rather than other performance indices. Statistical significance was calculated based on the null distribution obtained by the permutation test with 500 iterations. The statistical threshold was set to *P* < 0.05.

## List of Supplementary Materials

Materials and Methods

Fig. S1. The distribution of the ASD diagnosis probability in each imaging site in the discovery dataset and the ABIDE adult validation dataset.

Fig. S2. The relation between the head motion and performance index across the validation datasets.

Fig. S3. Results of mass-univariate analyses between the discovery and validation datasets. Fig. S4. Discriminative functional connections for the SCZ diagnosis.

Fig. S5. Discriminative functional connections for the MDD diagnosis.

Fig. S6. Relationship of the ASD and SCZ diagnosis on ASD and SCZ neuromarkers.

Fig. S7. Schematic representation of training and evaluation procedures of the ASD neuromarker.

Table S1. Demographic information of discovery and adult validation datasets.

Table S2. The classification performance of the ASD classifier in the discovery and adult validation datasets.

Table S3. Demographic information for the child and adolescent validation datasets.

Table S4. The classification performance of the ASD classifier in the child and adolescent datasets.

Table S5. The classification performance of the ASD classifier on the discovery dataset without ComBat harmonization method.

Table S6. The classification performance of the ASD classifier trained on SWA1 and UTO2 only.

Table S7. The classification performance of the ASD classifier trained on the SRPBS dataset only.

Table S8. Comparisons of classification and generalization performance between higher/lower resolutions of regions of interest.

Table S9. The list of discriminative FCs for the ASD diagnosis.

Table S10. The list of discriminative FCs reproducible across the five datasets. Table S11. Demographic information of SCZ and MDD datasets.

Table S12. The classification performance of the SCZ and MDD neuromarkers.

Table S13. The list of discriminative FCs for SCZ.

Table S14. The list of discriminative FCs for MDD.

Table S15. Image scanning protocols for R-fMRI data in the discovery dataset.

Table S16. Image scanning protocols for R-fMRI data in the validation datasets. References *(11, 12, 25, 36, 38-40, 42, 62-67, 72-76, 78-83)*

## Acknowledgments

We are very grateful to the participants in the SRPBS, ABIDE-I, ABIDE-II, and BMB projects.

## Funding

This study was supported by JSPS KAKENHI Grant Numbers JP21H05171 and JP21H05174, AMED under Grant Numbers JP19dm0207069, JP18dm0307001, JP18dm0307004, JP18dm0307008, and JP18dm0307009, Moonshot R&D Grant Number JPMJMS2021, and by UTokyo Institute for Diversity and Adaptation of Human Mind (UTIDAHM), and the International Research Center for Neurointelligence (WPI-IRCN) at The University of Tokyo Institutes for Advanced Study (UTIAS). ABIDE-I was supported by NIMH (K23MH087770), NIMH (R03MH096321), Leon Levy Foundation, Joseph P. Healy, and the Stavros Niarchos Foundation. ABIDE-II was supported by NIMH (5R21MH107045), NIMH (5R21MH107045), Nathan S. Kline Institute of Psychiatric Research, Joseph P. Healey, Phyllis Green, and Randolph Cowen.

## Author contributions

Conceptualization: TI, AY, MK, RH

Methodology: TI, AK

Investigation: TI, NY, JF, YY, MN, RA, MT, NI, GO, NO, KK, YO, HT

Visualization: TI, AY, MK, RH

Funding acquisition: GO, KK, YO, HT, MK, OY, RH

Project administration: RH

Supervision: MK, OY, RH

Writing – original draft: TI, AY, RH

Writing – review & editing: TI, AY, YT, NY, YYA, JF, YY, MN, RA, HO, YS, MT, NI, GO, NO, KK, SCT, HI, NK, YO, HT, MK, OY, RH

## Competing interests

MK is an inventor of patents owned by the Advanced Telecommunications Research Institute International related to the present work (PCT/JP2014/061544 [WO2014178323] and JP2015-228970/6195329). AY and MK are inventors of a patent application submitted by the Advanced Telecommunications Research Institute International related to the present work (JP2018-192842). YT is an employee of SHIONOGI & CO., LTD.

## Data and materials availability

All the discovery data, except for a part of the discovery dataset (i.e., SWA1 and UTO2), were available from the Decoded Neurofeedback (DecNef) Project Brain Data Repository (https://bicr-resource.atr.jp/decnefpro/). ABIDE-I and ABIDE-II were available from https://fcon_1000.projects.nitrc.org/indi/abide/. The BMB data will be available from its project website (https://hbm.brainminds-beyond.jp/).

## References

1. M. Parellada, Á. Andreu-Bernabeu, M. Burdeus, A. San José Cáceres, E. Urbiola, L. L. Carpenter, N. V. Kraguljac, W. M. McDonald, C. B. Nemeroff, C. I. Rodriguez, A. S. Widge, M. W. State, S. J. Sanders, In Search of Biomarkers to Guide Interventions in Autism Spectrum Disorder: A Systematic Review. Am. J. Psychiatry 180, 23–40 (2023).

2. S. Cortese, M. Solmi, G. Michelini, A. Bellato, C. Blanner, A. Canozzi, L. Eudave, L. C. Farhat, M. Højlund, O. Köhler-Forsberg, D. T. Leffa, C. Rohde, G. S. de Pablo, G. Vita, R. Wesselhoeft, J. Martin, S. Baumeister, N. S. Bozhilova, C. O. Carlisi, V. C. Leno, D. L. Floris, N. E. Holz, E. J. Kraaijenvanger, S. Sacu, I. Vainieri, G. Ostuzzi, C. Barbui, C. U. Correll, Candidate diagnostic biomarkers for neurodevelopmental disorders in children and adolescents: a systematic review. World Psychiatry 22, 129–149 (2023).

3. J. Hallmayer, S. Cleveland, A. Torres, J. Phillips, B. Cohen, T. Torigoe, J. Miller, A. Fedele, J. Collins, K. Smith, L. Lotspeich, L. A. Croen, S. Ozonoff, C. Lajonchere, J. K. Grether, N. Risch, Genetic heritability and shared environmental factors among twin pairs with autism. Arch. Gen. Psychiatry 68, 1095–1102 (2011).

4. E. Colvert, B. Tick, F. McEwen, C. Stewart, S. R. Curran, E. Woodhouse, N. Gillan, V. Hallett, S. Lietz, T. Garnett, A. Ronald, R. Plomin, F. Rijsdijk, F. Happé, P. Bolton, Heritability of Autism Spectrum Disorder in a UK Population-Based Twin Sample. JAMA Psychiatry 72, 415–423 (2015).

5. D. L. Floris, H. Peng, V. Warrier, M. V. Lombardo, C. M. Pretzsch, C. Moreau, A. Tsompanidis, W. Gong, M. Mennes, A. Llera, D. van Rooij, M. Oldehinkel, N. J. Forde, T. Charman, J. Tillmann, T. Banaschewski, C. Moessnang, S. Durston, R. J. Holt, C. Ecker, F. Dell’Acqua, E. Loth, T. Bourgeron, D. G. M. Murphy, A. F. Marquand, M.-C. Lai, J. K. Buitelaar, S. Baron-Cohen, C. F. Beckmann, APEX Group, EU-AIMS LEAP Group, The Link Between Autism and Sex-Related Neuroanatomy, and Associated Cognition and Gene Expression. *Am. J. Psychiatry*, appiajp20220194 (2022).

6. A. de Leeuw, F. Happé, R. A. Hoekstra, A Conceptual Framework for Understanding the Cultural and Contextual Factors on Autism Across the Globe. Autism Res. 13, 1029–1050 (2020).

7. C. Fountain, A. S. Winter, P. S. Bearman, Six developmental trajectories characterize children with autism. Pediatrics 129, e1112–20 (2012).

8. E. Waizbard-Bartov, E. Ferrer, B. Heath, S. J. Rogers, C. W. Nordahl, M. Solomon, D. G. Amaral, Identifying autism symptom severity trajectories across childhood. Autism Res. 15, 687–701 (2022).

9. N. Traut, K. Heuer, G. Lemaître, A. Beggiato, D. Germanaud, M. Elmaleh, A. Bethegnies, L. Bonnasse-Gahot, W. Cai, S. Chambon, F. Cliquet, A. Ghriss, N. Guigui, A. de Pierrefeu, M. Wang, V. Zantedeschi, A. Boucaud, J. van den Bossche, B. Kegl, R. Delorme, T. Bourgeron, R. Toro, G. Varoquaux, Insights from an autism imaging biomarker challenge: Promises and threats to biomarker discovery. Neuroimage 255, 119171 (2022).

10. J. Grove, S. Ripke, T. D. Als, M. Mattheisen, R. K. Walters, H. Won, J. Pallesen, E. Agerbo, O. A. Andreassen, R. Anney, S. Awashti, R. Belliveau, F. Bettella, J. D. Buxbaum, J. Bybjerg-Grauholm, M. Bækvad-Hansen, F. Cerrato, K. Chambert, J. H. Christensen, C. Churchhouse, K. Dellenvall, D. Demontis, S. De Rubeis, B. Devlin, S. Djurovic, A. L. Dumont, J. I. Goldstein, C. S. Hansen, M. E. Hauberg, M. V. Hollegaard, S. Hope, D. P. Howrigan, H. Huang, C. M. Hultman, L. Klei, J. Maller, J. Martin, A. R. Martin, J. L. Moran, M. Nyegaard, T. Nærland, D. S. Palmer, A. Palotie, C. B. Pedersen, M. G. Pedersen, T. dPoterba, J. B. Poulsen, B. S. Pourcain, P. Qvist, K. Rehnström, A. Reichenberg, J. Reichert, E. B. Robinson, K. Roeder, P. Roussos, E. Saemundsen, S. Sandin, F. K. Satterstrom, G. Davey Smith, H. Stefansson, S. Steinberg, C. R. Stevens, P. F. Sullivan, P. Turley, G. B. Walters, X. Xu, Autism Spectrum Disorder Working Group of the Psychiatric Genomics Consortium, BUPGEN, Major Depressive Disorder Working Group of the Psychiatric Genomics Consortium, 23andMe Research Team, K. Stefansson, D. H. Geschwind, M. Nordentoft, D. M. Hougaard, T. Werge, O. Mors, P. B. Mortensen, B. M. Neale, M. J. Daly, A.D. Børglum, Identification of common genetic risk variants for autism spectrum disorder. Nat. Genet. 51, 431–444 (2019).

11. A. Di Martino, C.-G. Yan, Q. Li, E. Denio, F. X. Castellanos, K. Alaerts, J. S. Anderson, M. Assaf, S. Y. Bookheimer, M. Dapretto, B. Deen, S. Delmonte, I. Dinstein, B. Ertl-Wagner, D. A. Fair, L. Gallagher, D. P. Kennedy, C. L. Keown, C. Keysers, J. E. Lainhart, C. Lord, B. Luna, V. Menon, N. J. Minshew, C. S. Monk, S. Mueller, R.-A. Müller, M. B. Nebel, J. T. Nigg, K. O’Hearn, K. A. Pelphrey, S. J. Peltier, J. D. Rudie, S. Sunaert, M. Thioux, J. M. Tyszka, L. Q. Uddin, J. S. Verhoeven, N. Wenderoth, J. L. Wiggins, S. H. Mostofsky, M. P. Milham, The autism brain imaging data exchange: towards a large-scale evaluation of the intrinsic brain architecture in autism. Mol. Psychiatry 19, 659–667 (2014).

12. A. Di Martino, D. O’Connor, B. Chen, K. Alaerts, J. S. Anderson, M. Assaf, J. H. Balsters, L. Baxter, A. Beggiato, S. Bernaerts, L. M. E. Blanken, S. Y. Bookheimer, B. B. Braden, L. Byrge, F. X. Castellanos, M. Dapretto, R. Delorme, D. A. Fair, I. Fishman, J. Fitzgerald, L. Gallagher, R. J. J. Keehn, D. P. Kennedy, J. E. Lainhart, B. Luna, S. H. Mostofsky, R.-A. Müller, M. B. Nebel, J. T. Nigg, K. O’Hearn, M. Solomon, R. Toro, C. J. Vaidya, N. Wenderoth, T. White, R. C. Craddock, C. Lord, B. Leventhal, M. P. Milham, Enhancing studies of the connectome in autism using the autism brain imaging data exchange II. Sci Data 4, 170010 (2017).

13. W. Feng, G. Liu, K. Zeng, M. Zeng, Y. Liu, A review of methods for classification and recognition of ASD using fMRI data. J. Neurosci. Methods 368, 109456 (2021).

14. C. Horien, D. L. Floris, A. S. Greene, S. Noble, M. Rolison, L. Tejavibulya, D. O’Connor, J. C. McPartland, D. Scheinost, K. Chawarska, E. M. R. Lake, R. T. Constable, Functional Connectome-Based Predictive Modeling in Autism. Biol. Psychiatry 92, 626–642 (2022).

15. T. Wolfers, D. L. Floris, R. Dinga, D. van Rooij, C. Isakoglou, S. M. Kia, M. Zabihi, A. Llera, R. Chowdanayaka, V. J. Kumar, H. Peng, C. Laidi, D. Batalle, R. Dimitrova, T. Charman, E. Loth, M.-C. Lai, E. Jones, S. Baumeister, C. Moessnang, T. Banaschewski, C. Ecker, G. Dumas, J. O’Muircheartaigh, D. Murphy, J. K. Buitelaar, A. F. Marquand, C. F. Beckmann, From pattern classification to stratification: towards conceptualizing the heterogeneity of Autism Spectrum Disorder. Neurosci. Biobehav. Rev. 104, 240–254 (2019).

16. C. P. Santana, E. A. de Carvalho, I. D. Rodrigues, G. S. Bastos, A. D. de Souza, L.L. de Brito, rs-fMRI and machine learning for ASD diagnosis: a systematic review and meta-analysis. Sci. Rep. 12, 6030 (2022).

17. W. H. Thompson, J. Wright, P. G. Bissett, R. A. Poldrack, Dataset decay and the problem of sequential analyses on open datasets. Elife 9, e53498 (2020).

18. A. Yamashita, N. Yahata, T. Itahashi, G. Lisi, T. Yamada, N. Ichikawa, M. Takamura, Y. Yoshihara, A. Kunimatsu, N. Okada, H. Yamagata, K. Matsuo, R. Hashimoto, G. Okada, Y. Sakai, J. Morimoto, J. Narumoto, Y. Shimada, K. Kasai, N. Kato, H. Takahashi, Y. Okamoto, S. C. Tanaka, M. Kawato, O. Yamashita, H. Imamizu, Harmonization of resting-state functional MRI data across multiple imaging sites via the separation of site differences into sampling bias and measurement bias. PLoS Biol. 17, e3000042 (2019).

19. A. A. Chen, J. C. Beer, N. J. Tustison, P. A. Cook, R. T. Shinohara, H. Shou, Alzheimer’s Disease Neuroimaging Initiative, Mitigating site effects in covariance for machine learning in neuroimaging data. Hum. Brain Mapp. 43, 1179–1195 (2022).

20. Y. Xie, Z. Xu, M. Xia, J. Liu, X. Shou, Z. Cui, X. Liao, Y. He, Alterations in Connectome Dynamics in Autism Spectrum Disorder: A Harmonized Mega- and Meta-Analysis Study Using the ABIDE Dataset. Biol. Psychiatry 0 (2021), doi:10.1016/j.biopsych.2021.12.004.

21. I. Ilioska, M. Oldehinkel, A. Llera, S. Chopra, T. Looden, R. Chauvin, D. Van Rooij, D. L. Floris, J. Tillmann, C. Moessnang, T. Banaschewski, R. J. Holt, E. Loth, T. Charman, D. G. M. Murphy, C. Ecker, M. Mennes, C. F. Beckmann, A. Fornito, J. K. Buitelaar, Connectome-wide mega-analysis reveals robust patterns of atypical functional connectivity in autism. Biol. Psychiatry 0 (2022), doi:10.1016/j.biopsych.2022.12.018.

22. G. Spera, A. Retico, P. Bosco, E. Ferrari, L. Palumbo, P. Oliva, F. Muratori, S. Calderoni, Evaluation of Altered Functional Connections in Male Children With Autism Spectrum Disorders on Multiple-Site Data Optimized With Machine Learning. Front. Psychiatry 10, 620 (2019).

23. Y. Duan, W. Zhao, C. Luo, X. Liu, H. Jiang, Y. Tang, C. Liu, D. Yao, Identifying and Predicting Autism Spectrum Disorder Based on Multi-Site Structural MRI With Machine Learning. Front. Hum. Neurosci. 15, 765517 (2021).

24. A. Abraham, M. P. Milham, A. Di Martino, R. C. Craddock, D. Samaras, B. Thirion, G. Varoquaux, Deriving reproducible biomarkers from multi-site resting-state data: An Autism-based example. Neuroimage 147, 736–745 (2017).

25. N. Yahata, J. Morimoto, R. Hashimoto, G. Lisi, K. Shibata, Y. Kawakubo, H. Kuwabara, M. Kuroda, T. Yamada, F. Megumi, H. Imamizu, J. E. Náñez Sr, H. Takahashi, Y. Okamoto, K. Kasai, N. Kato, Y. Sasaki, T. Watanabe, M. Kawato, A small number of abnormal brain connections predicts adult autism spectrum disorder. Nat. Commun. 7, 11254 (2016).

26. A. Jahedi, C. A. Nasamran, B. Faires, J. Fan, R.-A. Müller, Distributed Intrinsic Functional Connectivity Patterns Predict Diagnostic Status in Large Autism Cohort. Brain Connect. 7, 515–525 (2017).

27. K. Supekar, S. Ryali, R. Yuan, D. Kumar, C. de los Angeles, V. Menon, Robust, Generalizable, and Interpretable Artificial Intelligence–Derived Brain Fingerprints of Autism and Social Communication Symptom Severity. Biol. Psychiatry 92, 643–653 (2022).

28. L. Q. Uddin, K. Supekar, V. Menon, Reconceptualizing functional brain connectivity in autism from a developmental perspective. Front. Hum. Neurosci. 7, 458 (2013).

29. K. Alaerts, K. Nayar, C. Kelly, J. Raithel, M. P. Milham, A. Di Martino, Age-related changes in intrinsic function of the superior temporal sulcus in autism spectrum disorders. Soc. Cogn. Affect. Neurosci. 10, 1413–1423 (2015).

30. G. L. Wallace, N. Dankner, L. Kenworthy, J. N. Giedd, A. Martin, Age-related temporal and parietal cortical thinning in autism spectrum disorders. Brain 133, 3745–3754 (2010).

31. A. Kazeminejad, R. C. Sotero, Topological Properties of Resting-State fMRI Functional Networks Improve Machine Learning-Based Autism Classification. Front. Neurosci. 12, 1018 (2018).

32. P. Lanka, D. Rangaprakash, M. N. Dretsch, J. S. Katz, T. S. Denney Jr, G. Deshpande, Supervised machine learning for diagnostic classification from large-scale neuroimaging datasets. Brain Imaging Behav. 14, 2378–2416 (2020).

33. Š. Holiga, J. F. Hipp, C. H. Chatham, P. Garces, W. Spooren, X. L. D’Ardhuy, A. Bertolino, C. Bouquet, J. K. Buitelaar, C. Bours, A. Rausch, M. Oldehinkel, M. Bouvard, A. Amestoy, M. Caralp, S. Gueguen, M. Ly-Le Moal, J. Houenou, C. F. Beckmann, E. Loth, D. Murphy, T. Charman, J. Tillmann, C. Laidi, R. Delorme, A. Beggiato, A. Gaman, I. Scheid, M. Leboyer, M.-A. d’Albis, J. Sevigny, C. Czech, F. Bolognani, G. D. Honey, J. Dukart, Patients with autism spectrum disorders display reproducible functional connectivity alterations. Sci. Transl. Med. 11 (2019), doi:10.1126/scitranslmed.aat9223.

34. Y.-H. Tung, H.-Y. Lin, C.-L. Chen, C.-Y. Shang, L.-Y. Yang, Y.-C. Hsu, W.-Y. I. Tseng, S. S.-F. Gau, Whole Brain White Matter Tract Deviation and Idiosyncrasy From Normative Development in Autism and ADHD and Unaffected Siblings Link With Dimensions of Psychopathology and Cognition. Am. J. Psychiatry 178, 730–743 (2021).

35. Y. Aoki, Y. N. Yoncheva, B. Chen, T. Nath, D. Sharp, M. Lazar, P. Velasco, M. P. Milham, A. Di Martino, Association of White Matter Structure With Autism Spectrum Disorder and Attention-Deficit/Hyperactivity Disorder. JAMA Psychiatry 74, 1120–1128 (2017).

36. Y. Yoshihara, G. Lisi, N. Yahata, J. Fujino, Y. Matsumoto, J. Miyata, G.-I. Sugihara, S.-I. Urayama, M. Kubota, M. Yamashita, R. Hashimoto, N. Ichikawa, W. Cahn, N. E. M. van Haren, S. Mori, Y. Okamoto, K. Kasai, N. Kato, H. Imamizu, R. S. Kahn, A. Sawa, M. Kawato, T. Murai, J. Morimoto, H. Takahashi, Overlapping but Asymmetrical Relationships Between Schizophrenia and Autism Revealed by Brain Connectivity. Schizophr. Bull. (2020), doi:10.1093/schbul/sbaa021.

37. M.-C. Lai, C. Kassee, R. Besney, S. Bonato, L. Hull, W. Mandy, P. Szatmari, S. H. Ameis, Prevalence of co-occurring mental health diagnoses in the autism population: a systematic review and meta-analysis. Lancet Psychiatry 6, 819–829 (2019).

38. S. C. Tanaka, A. Yamashita, N. Yahata, T. Itahashi, G. Lisi, T. Yamada, N. Ichikawa, M. Takamura, Y. Yoshihara, A. Kunimatsu, N. Okada, R. Hashimoto, G. Okada, Y. Sakai, J. Morimoto, J. Narumoto, Y. Shimada, H. Mano, W. Yoshida, B. Seymour, T. Shimizu, K. Hosomi, Y. Saitoh, K. Kasai, N. Kato, H. Takahashi, Y. Okamoto, O. Yamashita, M. Kawato, H. Imamizu, A multi-site, multi-disorder resting-state magnetic resonance image database. Sci Data 8, 227 (2021).

39. A. Schaefer, R. Kong, E. M. Gordon, T. O. Laumann, X.-N. Zuo, A. J. Holmes, S. B. Eickhoff, B. T. T. Yeo, Local-Global Parcellation of the Human Cerebral Cortex from Intrinsic Functional Connectivity MRI. Cereb. Cortex 28, 3095–3114 (2018).

40. A. Yamashita, Y. Sakai, T. Yamada, N. Yahata, A. Kunimatsu, N. Okada, T. Itahashi, R. Hashimoto, H. Mizuta, N. Ichikawa, M. Takamura, G. Okada, H. Yamagata, K. Harada, K. Matsuo, S. C. Tanaka, M. Kawato, K. Kasai, N. Kato, H. Takahashi, Y. Okamoto, O. Yamashita, H. Imamizu, Generalizable brain network markers of major depressive disorder across multiple imaging sites. PLoS Biol. 18, e3000966 (2020).

41. E. M. R. Lake, E. S. Finn, S. M. Noble, T. Vanderwal, X. Shen, M. D. Rosenberg, M. N. Spann, M. M. Chun, D. Scheinost, R. T. Constable, The Functional Brain Organization of an Individual Allows Prediction of Measures of Social Abilities Transdiagnostically in Autism and Attention-Deficit/Hyperactivity Disorder. Biol. Psychiatry 86, 315–326 (2019).

42. M. Yamashita, M. Kawato, H. Imamizu, Predicting learning plateau of working memory from whole-brain intrinsic network connectivity patterns. Sci. Rep. 5, 7622 (2015).

43. D. Scheinost, S. Noble, C. Horien, A. S. Greene, E. M. Lake, M. Salehi, S. Gao, X. Shen, D. O’Connor, D. S. Barron, S. W. Yip, M. D. Rosenberg, R. T. Constable, Ten simple rules for predictive modeling of individual differences in neuroimaging. Neuroimage 193, 35–45 (2019).

44. A. Padmanabhan, A. Lynn, W. Foran, B. Luna, K. O’Hearn, Age related changes in striatal resting state functional connectivity in autism. Front. Hum. Neurosci. 7, 814 (2013).

45. L. Cerliani, M. Mennes, R. M. Thomas, A. Di Martino, M. Thioux, C. Keysers, Increased Functional Connectivity Between Subcortical and Cortical Resting-State Networks in Autism Spectrum Disorder. JAMA Psychiatry 72, 767–777 (2015).

46. S. Park, K. V. Haak, H. B. Cho, S. L. Valk, R. A. I. Bethlehem, M. P. Milham, B. C. Bernhardt, A. Di Martino, S.-J. Hong, Atypical integration of sensory-to-transmodal functional systems mediates symptom severity in autism. Front. Psychiatry 12 (2021), doi:10.3389/fpsyt.2021.699813.

47. S.-J. Hong, R. Vos de Wael, R. A. I. Bethlehem, S. Lariviere, C. Paquola, S. L. Valk, M. P. Milham, A. Di Martino, D. S. Margulies, J. Smallwood, B. C. Bernhardt, Atypical functional connectome hierarchy in autism. Nat. Commun. 10, 1022 (2019).

48. B. E. Yerys, E. M. Gordon, D. N. Abrams, T. D. Satterthwaite, R. Weinblatt, K. F. Jankowski, J. Strang, L. Kenworthy, W. D. Gaillard, C. J. Vaidya, Default mode network segregation and social deficits in autism spectrum disorder: Evidence from non-medicated children. Neuroimage Clin 9, 223–232 (2015).

49. A. Padmanabhan, C. J. Lynch, M. Schaer, V. Menon, The Default Mode Network in Autism. Biol Psychiatry Cogn Neurosci Neuroimaging 2, 476–486 (2017).

50. O. Benkarim, C. Paquola, B.-Y. Park, V. Kebets, S.-J. Hong, R. Vos de Wael, S. Zhang, B. T. T. Yeo, M. Eickenberg, T. Ge, J.-B. Poline, B. C. Bernhardt, D. Bzdok, Population heterogeneity in clinical cohorts affects the predictive accuracy of brain imaging. PLoS Biol. 20, e3001627 (2022).

51. A. Kaneko, R. Ohshima, H. Noda, T. Matsumaru, R. Iwanaga, M. Ide, Sensory and social subtypes of Japanese individuals with autism spectrum disorders (2021), doi:10.31234/osf.io/umvcn.

52. M. D. Thye, H. M. Bednarz, A. J. Herringshaw, E. B. Sartin, R. K. Kana, The impact of atypical sensory processing on social impairments in autism spectrum disorder. Dev. Cogn. Neurosci. 29, 151–167 (2018).

53. Cross-Disorder Group of the Psychiatric Genomics Consortium, Identification of risk loci with shared effects on five major psychiatric disorders: a genome-wide analysis. Lancet 381, 1371–1379 (2013).

54. H. Chen, L. Q. Uddin, X. Duan, J. Zheng, Z. Long, Y. Zhang, X. Guo, Y. Zhang, J. Zhao, H. Chen, Shared atypical default mode and salience network functional connectivity between autism and schizophrenia. Autism Res. 10, 1776–1786 (2017).

55. L. D. Oliver, I. Moxon-Emre, M.-C. Lai, L. Grennan, A. N. Voineskos, S. H. Ameis, Social Cognitive Performance in Schizophrenia Spectrum Disorders Compared With Autism Spectrum Disorder: A Systematic Review, Meta-analysis, and Meta-regression. JAMA Psychiatry 78, 281–292 (2021).

56. A. Jutla, J. Foss-Feig, J. Veenstra-VanderWeele, Autism spectrum disorder and schizophrenia: An updated conceptual review. Autism Res. 15, 384–412 (2022).

57. W. Zhao, V. Voon, K. Xue, C. Xie, J. Kang, C.-P. Lin, J. Wang, J. Cheng, J. Feng, Common abnormal connectivity in first-episode and chronic schizophrenia in pre- and post-central regions: Implications for neuromodulation targeting. Prog. Neuropsychopharmacol. Biol. Psychiatry 117, 110556 (2022).

58. J. Hogeveen, M. K. Krug, M. V. Elliott, M. Solomon, Insula-Retrosplenial Cortex Overconnectivity Increases Internalizing via Reduced Insight in Autism. Biol. Psychiatry 84, 287–294 (2018).

59. N. M. Kleinhans, M. A. Reiter, E. Neuhaus, G. Pauley, N. Martin, S. Dager, A. Estes, Subregional differences in intrinsic amygdala hyperconnectivity and hypoconnectivity in autism spectrum disorder. Autism Res. 9, 760–772 (2016).

60. S. G. W. Urchs, A. Tam, P. Orban, C. Moreau, Y. Benhajali, H. D. Nguyen, A. C. Evans, P. Bellec, Functional connectivity subtypes associate robustly with ASD diagnosis. Elife 11 (2022), doi:10.7554/eLife.56257.

61. T. Yamada, R.-I. Hashimoto, N. Yahata, N. Ichikawa, Y. Yoshihara, Y. Okamoto, N. Kato, H. Takahashi, M. Kawato, Resting-State Functional Connectivity-Based Biomarkers and Functional MRI-Based Neurofeedback for Psychiatric Disorders: A Challenge for Developing Theranostic Biomarkers. Int. J. Neuropsychopharmacol. 20, 769–781 (2017).

62. S. Koike, S. C. Tanaka, T. Okada, T. Aso, A. Yamashita, O. Yamashita, M. Asano, N. Maikusa, K. Morita, N. Okada, M. Fukunaga, A. Uematsu, H. Togo, A. Miyazaki, K. Murata, Y. Urushibata, J. Autio, T. Ose, J. Yoshimoto, T. Araki, M. F. Glasser, D. C. Van Essen, M. Maruyama, N. Sadato, M. Kawato, K. Kasai, Y. Okamoto, T. Hanakawa, T. Hayashi, Brain/MINDS Beyond Human Brain MRI Group, Brain/MINDS beyond human brain MRI project: A protocol for multi-level harmonization across brain disorders throughout the lifespan. Neuroimage Clin 30, 102600 (2021).

63. O. Esteban, C. J. Markiewicz, R. W. Blair, C. A. Moodie, A. I. Isik, A. Erramuzpe, J. D. Kent, M. Goncalves, E. DuPre, M. Snyder, H. Oya, S. S. Ghosh, J. Wright, J. Durnez, R. A. Poldrack, K. J. Gorgolewski, fMRIPrep: a robust preprocessing pipeline for functional MRI. Nat. Methods 16, 111–116 (2019).

64. E. W. Dickie, A. Anticevic, D. E. Smith, T. S. Coalson, M. Manogaran, N. Calarco, J. D. Viviano, M. F. Glasser, D. C. Van Essen, A. N. Voineskos, Ciftify: A framework for surface-based analysis of legacy MR acquisitions. Neuroimage 197, 818–826 (2019).

65. J. D. Power, K. A. Barnes, A. Z. Snyder, B. L. Schlaggar, S. E. Petersen, Spurious but systematic correlations in functional connectivity MRI networks arise from subject motion. Neuroimage 59, 2142–2154 (2012).

66. M. F. Glasser, T. S. Coalson, E. C. Robinson, C. D. Hacker, J. Harwell, E. Yacoub, K. Ugurbil, J. Andersson, C. F. Beckmann, M. Jenkinson, S. M. Smith, D. C. Van Essen, A multi-modal parcellation of human cerebral cortex. Nature 536, 171–178 (2016).

67. B. T. T. Yeo, F. M. Krienen, J. Sepulcre, M. R. Sabuncu, D. Lashkari, M. Hollinshead, J. L. Roffman, J. W. Smoller, L. Zöllei, J. R. Polimeni, B. Fischl, H. Liu, R. L. Buckner, The organization of the human cerebral cortex estimated by intrinsic functional connectivity. J. Neurophysiol. 106, 1125–1165 (2011).

68. W. E. Johnson, C. Li, A. Rabinovic, Adjusting batch effects in microarray expression data using empirical Bayes methods. Biostatistics 8, 118–127 (2007).

69. J.-P. Fortin, N. Cullen, Y. I. Sheline, W. D. Taylor, I. Aselcioglu, P. A. Cook, P. Adams, C. Cooper, M. Fava, P. J. McGrath, M. McInnis, M. L. Phillips, M. H. Trivedi, M. M. Weissman, R. T. Shinohara, Harmonization of cortical thickness measurements across scanners and sites. Neuroimage 167, 104–120 (2018).

70. J.-P. Fortin, D. Parker, B. Tunç, T. Watanabe, M. A. Elliott, K. Ruparel, D. R. Roalf, T. D. Satterthwaite, R. C. Gur, R. E. Gur, R. T. Schultz, R. Verma, R. T. Shinohara, Harmonization of multi-site diffusion tensor imaging data. Neuroimage 161, 149–170 (2017).

71. M. Yu, K. A. Linn, P. A. Cook, M. L. Phillips, M. McInnis, M. Fava, M. H. Trivedi, M. M. Weissman, R. T. Shinohara, Y. I. Sheline, Statistical harmonization corrects site effects in functional connectivity measurements from multi-site fMRI data. Hum. Brain Mapp. 39, 4213– 4227 (2018).

72. N. Ichikawa, G. Lisi, N. Yahata, G. Okada, M. Takamura, R.-I. Hashimoto, T. Yamada, M. Yamada, T. Suhara, S. Moriguchi, M. Mimura, Y. Yoshihara, H. Takahashi, K. Kasai, N. Kato, S. Yamawaki, B. Seymour, M. Kawato, J. Morimoto, Y. Okamoto, Primary functional brain connections associated with melancholic major depressive disorder and modulation by antidepressants. Sci. Rep. 10, 3542 (2020).

73. F. Almuqhim, F. Saeed, ASD-SAENet: A Sparse Autoencoder, and Deep-Neural Network Model for Detecting Autism Spectrum Disorder (ASD) Using fMRI Data. Front. Comput. Neurosci. 15, 654315 (2021).

74. H. Kwon, J. I. Kim, S.-Y. Son, Y. H. Jang, B.-N. Kim, H. J. Lee, J.-M. Lee, Sparse Hierarchical Representation Learning on Functional Brain Networks for Prediction of Autism Severity Levels. Front. Neurosci. 16, 935431 (2022).

75. R. Tibshirani, Regression shrinkage and selection via the lasso. J. R. Stat. Soc. 58, 267–288 (1996).

76. D. Chicco, Ten quick tips for machine learning in computational biology. BioData Min. 10, 35 (2017).

77. S. Holm, A Simple Sequentially Rejective Multiple Test Procedure. Scand. Stat. Theory Appl. 6, 65–70 (1979).

78. L. Parkes, B. Fulcher, M. Yücel, A. Fornito, An evaluation of the efficacy, reliability, and sensitivity of motion correction strategies for resting-state functional MRI. Neuroimage 171, 415–436 (2018).

79. H. Chen, J. S. Nomi, L. Q. Uddin, X. Duan, H. Chen, Intrinsic functional connectivity variance and state-specific under-connectivity in autism. Hum. Brain Mapp. 38, 5740–5755 (2017).

80. M. F. Glasser, S. N. Sotiropoulos, J. A. Wilson, T. S. Coalson, B. Fischl, J. L. Andersson, J. Xu, S. Jbabdi, M. Webster, J. R. Polimeni, D. C. Van Essen, M. Jenkinson, The minimal preprocessing pipelines for the Human Connectome Project. Neuroimage 80, 105–124 (2013).

81. B. C. Wallace, K. Small, C. E. Brodley, T. A. Trikalinos, in 2011 IEEE 11th International Conference on Data Mining, (2011), pp. 754–763.

82. B. W. Matthews, Comparison of the predicted and observed secondary structure of T4 phage lysozyme. Biochim. Biophys. Acta 405, 442–451 (1975).

83. B. Yamagata, T. Itahashi, J. Fujino, H. Ohta, M. Nakamura, N. Kato, M. Mimura, R.-I. Hashimoto, Y. Aoki, Machine learning approach to identify a resting-state functional connectivity pattern serving as an endophenotype of autism spectrum disorder. Brain Imaging Behav. (2018), doi:10.1007/s11682-018-9973-2.

