## Supplementary Materials for "Generalizable neuromarker for autism spectrum disorder across imaging sites and developmental stages: A multi-site study"

**This PDF file includes:**

Materials and Methods

Fig. S1. The distribution of the ASD diagnosis probability in each imaging site in the discovery dataset and the ABIDE adult validation dataset.

Fig. S2. The relation between the head motion and performance index across the validation datasets.

Fig. S3. Results of mass-univariate analyses between the discovery and validation datasets.

Fig. S4. Discriminative functional connections for the SCZ diagnosis.

Fig. S5. Discriminative functional connections for the MDD diagnosis.

Fig. S6. Relationship of the ASD and SCZ diagnosis on ASD and SCZ neuromarkers.

Fig. S7. Schematic representation of training and evaluation procedures of the ASD neuromarker.

Table S1. Demographic information of discovery and adult validation datasets.

Table S2. The classification performance of the ASD classifier in the discovery and adult validation datasets.

Table S3. Demographic information for the child and adolescent validation datasets.

Table S4. The classification performance of the ASD classifier in the child and adolescent datasets.

Table S5. The classification performance of the ASD classifier on the discovery dataset without ComBat harmonization method.

Table S6. The classification performance of the ASD classifier trained on SWA1 and UTO2 only.

Table S7. The classification performance of the ASD classifier trained on the SRPBS dataset only.

Table S8. Comparisons of classification and generalization performance between higher/lower resolutions of regions of interest.

Table S9. The list of discriminative FCs for the ASD diagnosis.

Table S10. The list of discriminative FCs reproducible across the five datasets.

Table S11. Demographic information of SCZ and MDD datasets.

Table S12. The classification performance of the SCZ and MDD neuromarkers.

Table S13. The list of discriminative FCs for SCZ.

Table S14. The list of discriminative FCs for MDD.

Table S15. Image scanning protocols for R-fMRI data in the discovery dataset.

Table S16. Image scanning protocols for R-fMRI data in the validation datasets.

References *(11, 12, 25, 36, 38- 40, 42, 62-67, 72-76,  78-83)*

### Supplementary Materials

#### Materials and Methods

##### Ethics of statement

All participants (if appropriate) and their parent/legal guardian provided written informed consent. The institutional review boards approved recruitment procedures and experimental protocols at the principal investigators' respective institutions. These procedures were conducted in accordance with the Declaration of Helsinki.

##### Participants

The current study used three adult resting-state fMRI (R-fMRI) datasets for the analyses: one was used as the discovery dataset, and the remaining two were used as independent validation datasets. Tables S1 and S15 show the demographic information and scanning parameters for the three datasets.

*Discovery dataset*

The discovery dataset contained data of 550 typically developing controls (TDCs) from five scanners at four imaging sites (University of Tokyo [UTO1 and UTO2], Kyoto University [KUT], Center for Innovation in Hiroshima University (COI), and Showa University [SWA1]) and 180 adults with autism spectrum disorder (ASD) from two institutes (SWA1 and UTO2). This dataset consisted of a part of the Strategic Research Program for the Promotion of Brain Science (SRPBS) dataset (https://bicr-resource.atr.jp/srpbsopen/) *(38)*. We also included participants at SWA1 and a part of the dataset used in our prior study *(25)*.

*The ABIDE adult validation dataset*

The first independent adult validation dataset consisted of participants from the Autism Brain Imaging Data Exchange I (ABIDE-I) *(11)* and -II datasets *(12)*. Since adult participants were limited in both releases, we combined the two datasets. Of note, since some imaging sites participated in both datasets (e.g., New York University Langone Medical Center [NYU] and the University of Utah School of Medicine [USM]), we checked the consistency of the scanning protocols in each imaging site. We used 54 adults with ASD and 67 TDCs that were selected from the following three sites: the University of Leuven (Leuven), NYU, and USM. In this study, we referred this validation dataset to the ABIDE adult dataset.

*The Japanese adult validation dataset*

To further validate the generalization performance of our classifier, we newly collected R-fMRI data from 22 adults with ASD and 38 controls at Showa University, Karasuyama Hospital (SWA2). The data were collected under the Brain/MINDS Beyond project (BMB; https://brainminds-beyond.jp/). The project details were described elsewhere *(62)*.

**Exclusion criteria**

We applied the following exclusion criteria for participant selection; 1) participants with no whole-brain coverage were excluded; 2) participants who had less than 4 min of uncontaminated R-fMRI data *(78)*; 3) participants with errors in any preprocessing steps (e.g., segmentation failure and failures in spatial normalization and surface mapping). For the ABIDE adult validation dataset, we excluded imaging sites that contained less than 10 participants per group, according to prior studies *(79)*. We also excluded imaging sites that did not have some missing information about the scanning protocol (e.g., slice acquisition order).

**R-fMRI data preprocessing**

We preprocessed all the R-fMRI data using fMRIPrep version 1.1.8 *(63)*. The fMRIPrep performs a series of preprocessing steps, including head motion estimation, slice timing correction, co-registration of EPI data to the corresponding T1-weighted anatomical image, distortion correction, and normalization to a standard Montreal Neurological Institute (MNI) space. We used the "fieldmap-less" distortion correction method if the fieldmap data were unavailable. We used the ciftify toolbox version 2.1.1 *(64)* to map the preprocessed data onto the grayordinate *(80)*.

For each vertex, we performed nuisance regression to remove the effects of artifactual and non-neural sources. Nuisance regressors consisted of six head-motion parameters, averaged signals from subject-specific white matter and cerebrospinal fluid masks, global signal, their temporal derivatives, and linear detrending. After nuisance regression, we applied a band-pass filter (0.008–0.1 Hz) to the residuals. We computed frame-wise displacement (FD) *(65)* for each participant to characterize the frame-by-frame head motion during the scans. We used FD as a measure for detecting occasional head movement. To reduce spurious changes in FC due to head motion, we removed volumes with FD > 0.5 mm, as proposed in a previous study *(65)*.

**Parcellation and network construction**

We used Glasser's 379 surface-based brain parcellations (cortical 360 parcellations and subcortical 19 parcellations) as ROIs *(66)*. We extracted the averaged, denoised signals from these ROIs. We computed the temporal correlations of signals among all possible pairs of ROIs and applied Fisher's $r$-to-$z$ transformation, resulting in 71,631 unique FCs for each participant. Because the label of each ROI in Glasser's atlas was not intuitive, we utilized Yeo's resting-state network (RSN) labels *(67)* to assign important ROIs to the corresponding RSN label. This study added the subcortical network label to the subcortical and cerebellar regions.

**Construction of the ASD neuromarker using the discovery dataset**

Based on previous studies *(25, 36, 40, 72–74)*, we assumed that psychiatric disorder factors were associated with the limited number of FCs, rather than the whole-brain connections. We, thus, used a logistic regression analysis with least absolute shrinkage and selection operator (LASSO) method that selects an optimal subset of FCs from the whole brain connections *(75)*. The details of our procedures were described elsewhere *(40)*. Briefly, a logistic function is used to define the probability of a participant belonging to the ASD class label as follows:

$$P_i(y_i = 1 \mid x_i;\, w) = \frac{1}{1 + exp\,(-x_i \cdot w)},$$

where $y_i$ and $x_i$ represent the $i$-th participant's class label and FC vector, respectively. The class label was set to 1 if the participant belonged to the ASD group while setting to 0 if the participant belonged to the TDC group. The weight vector was denoted as $w$. The weight vector was optimized by minimizing the following objective function:

$$L(w) = -\frac{1}{N}\sum_{i=1}^{N} \quad log\,\big(P_i(y_i = 1 \mid x_i;\, w)\big) + \lambda \cdot ||w||_1,$$

where $||\cdot||_1$ represents $L_1$-norm and $\lambda$ stands for a hyperparameter that regulates the sparsity of the weight vector.

We developed the ASD neuromarker using the LASSO method with 10-fold nested cross-validation (CV) and 10 subsampling, yielding 100 trained classifiers (fig. S7). To estimate the optimal weights and tune the hyperparameter, we used a 10-fold nested CV procedure with an undersampling method. In this procedure, we first divided the whole discovery dataset into training (9 folds of 10 folds) and test (1 fold of 10 folds) datasets using the "*cvpartition*" implemented in MATLAB (R2020b, Mathworks, USA). We then applied an undersampling method to alleviate a bias due to the imbalance in the number of participants between the groups *(81)*. In this undersampling method, we randomly selected participants from the discovery dataset to match the number of participants between the ASD and TDC groups. Similar to our previous study, we matched the mean age between ASD and TDC groups in each subsample. To avoid any subsampling bias, we repeated this undersampling ten times, yielding ten subsampled matched training datasets. For each subsampled dataset, we fitted the logistic regression model while tuning the hyperparameter in the inner loop of the nested CV. For building a logistic regression model, we used the "*lassoglm*" function implemented in MATLAB. The inner CV (i.e., "CV" parameter in the lassoglm function) was set to 10, and "NumLambda" was set to 25. We determined the optimal $\lambda$ according to the one standard error rule in which we selected the largest $\lambda$ within the standard deviation of the minimum prediction error among all $\lambda$. The mean classifier output value was considered as diagnostic probability, indicating a likelihood of a participant belonging to the ASD class. We considered participants as those with ASD if their diagnostic probability values were higher than 0.5. We calculated the area under the curve (AUC) to assess the classification performance using the "*perfcurve*" implemented in MATLAB. We also computed accuracy, sensitivity, specificity, and the Matthews correlation coefficient (MCC). The MCC is suitable for the imbalanced dataset because this metric takes into account the ratio of the confusion matrix size *(76, 82)*. We used AUC and MCC as performance indices throughout the paper.

**The details of the effects of head motion, harmonization, and experimental settings on the generalization performance**

To assess the impacts of head motion, harmonization, and experimental factors (i.e., diversity in the characteristics of scanning protocols and choice of atlas), we conducted several control analyses. We first assessed the association between head motion and classification performance. We calculated the value of the area under the curve (AUC) and the mean frame-wise displacement (FD) value at each imaging site in each validation dataset. We then calculated the Pearson correlation coefficient between the mean FD and AUC values across the validation datasets. A significant positive correlation was not found between the AUC and mean FD ($r$ = -0.56, $P$ = 0.002; fig. S2). This result indicates that the head motion did not artificially improve the generalizability of our neuromarker.

We next assessed the effects of the harmonization method on classification performance in the discovery and adult validation datasets, respectively. We trained an ASD classifier on the discovery dataset without the ComBat harmonization method. The ASD classifier showed better classification performance (accuracy = 77%, AUC = 0.85, and MCC = 0.50) in the discovery dataset (table S5). The ASD classifier, however, exhibited reduced generalizability to the ABIDE adult (accuracy = 60%, AUC = 0.64, and MCC = 0.19) and the Japanese adult (accuracy = 67%, AUC = 0.73, and MCC = 0.23). These results suggest that applying the ComBat harmonization method plays a vital role in improving the generalizability of the ASD classifier to other datasets.

We further investigated the impacts of diversity in imaging sites on the generalization performance since our discovery dataset comprised imbalanced imaging sites (i.e., COI, KUT, and UTO1) and a site with a different scanning protocol (i.e., UTO2). Three imaging sites (i.e., COI, KUT, and UTO1) contained the TDC population only, and thus there is a possibility that the inclusion of imbalanced data might influence the classifier's generalization performance. To test this, we built a neuromarker for the ASD diagnosis using two imaging sites (i.e., SWA1 and UTO2) by removing the three sites. The trained classifier showed a slightly reduced accuracy of 74%, an AUC of 0.80, and an MCC of 0.47 in the discovery dataset (table S6). The trained classifier did not change the discrimination abilities in the ABIDE adult (accuracy = 58%, AUC = 0.67, and MCC = 0.21) and the Japanese adult (accuracy = 72%, AUC = 0.83, MCC = 0.42). We next investigated the impacts of an imaging site with a different scanning protocol (i.e., UTO2) on the classification and generalization performance. We repeated the same analyses while excluding the UTO2. We did not observe significant changes in the classification and generalization performance (table S7). These results suggest that, at least in our experimental setting, the inclusion of diverse imaging sites does not influence the generalization performance of our ASD neuromarker.

Finally, we assessed whether our generalization performance was not atlas-dependent. To test this, we used Schaefer's cortical atlas *(39)* as an alternative to the Glasser atlas *(66)* because this atlas provided an atlas with multiple levels of resolutions ranging from 100 to 1,000 and thus it is suitable to investigate the effects of ROI resolutions on the generalization performance. Of note, we excluded 1,000 ROIs from analyses because some brain regions could not extract signals reliably. Classifiers with Schaefer's atlas exhibited similar or higher generalization performance in the ABIDE adult and Japanese adult datasets (table S8). These results suggest that our generalization performance was not atlas-dependent.

**Consistency of the effect of ASD diagnosis across the datasets**

In the mass-univariate analyses, we did not observe a significant positive correlation between the discovery dataset and the child dataset (see **Results** in the main text and fig. S3). We speculated that, instead of the whole-brain pattern, a specific set of important FCs were reproducible between the discovery dataset and the child dataset. To test this, we used a binomial test, similar to previous studies *(42, 83)*. We calculated a *t*-value as the effect of the ASD diagnosis in each FC of each dataset. We then counted the number of FCs showing the same sign (i.e., the same direction of the effect of diagnosis) within the set of discriminative FCs, $k$, and the whole FCs, $m$, respectively. Since the permutation test identified 141 FCs as important FCs for the ASD diagnosis (see **Results** in the main text) and 98 out of 141 FCs showed the same direction of between-group difference between the discovery dataset and the child dataset (i.e., the same sign of *t*-values between the datasets), we set $n = 141$ (the number of important FCs), $k = 98$ (the number of FCs showing the same sign of *t*-values within the important FCs), and $m = 35{,}443$ (the number of FCs showing the same direction of between-group difference in the whole connections), respectively. The binomial test confirmed that the set of important FCs was reproducible between the discovery dataset and the child dataset ($P < 0.05$). This observation was replicated between the discovery dataset and the adult and adolescent datasets ($k = 94$ and $m = 38{,}969$ for the ABIDE adult; $k = 105$ and $m = 42{,}649$ for the Japanese adult; $k = 98$ and $m = 40{,}124$ for the adolescent dataset; all $P < 0.05$). These results suggest that the reproducible FCs selected by our method might contribute to the generalizability of our neuromarker.

**Dimensional relationships among three psychiatric disorders**

We examined the exchangeability among the three psychiatric disorders, constructing classifiers for the SCZ and MDD diagnoses and applying these classifiers to the remaining psychiatric disorders. We used the same analytical procedures, including 10-fold nested CV, 10 subsampling, and parameter settings for the LASSO method. We then calculated AUC, accuracy, sensitivity, specificity, and MCC as performance indices. To test the statistical significance, we constructed null distributions using permutation tests with 100 iterations. We set iterations to 100 instead of 500 because this procedure was computationally expensive. Statistical significance was set to $P < 0.05$.

We tested the sensitivity of each classifier to the remaining psychiatric disorders by applying each classifier to other psychiatric disorders. Since participants with TDC were identical to those in the discovery dataset, we focused on sensitivity instead of other performance indices. We constructed null distributions of performance indices using a permutation test with 100 iterations and applied a statistical threshold of $P < 0.05$, one-sided.

**Cross-validated neuromarkers for schizophrenia and major depressive disorder**

Similar to the ASD classifier, we used a 10-fold nested CV with 10 subsamples to determine the weights and hyperparameters. The statistical significance of both classifiers was confirmed by permutation tests with 100 iterations. The SCZ classifier exhibited an accuracy of 82%, AUC of 0.89, and MCC of 0.51 (permutation test, all $P < 0.05$; table S12). The corresponding sensitivity and specificity were 83% and 82%, respectively. On the other hand, the MDD classifier exhibited an accuracy of 68%, AUC of 0.78, and MCC of 0.32 (permutation test, $P < 0.05$; table S12). The corresponding sensitivity and specificity were 75% and 66%, respectively. These results indicate that both classifiers hold acceptable classification performance to the training dataset.

**Identification of discriminative FCs for SCZ and MDD**

Similar to identifying discriminative FCs for the ASD diagnosis, we used permutation tests with 100 iterations to identify important FCs for schizophrenia (SCZ) and major depressive disorder (MDD), respectively. At each iteration, we shuffled the diagnostic labels to create a permuted dataset and constructed permuted classifiers. We used the number of counts for each FC selected by the LASSO across 10-fold cross-validation and 10 subsampling (i.e., across 100 classifiers). To control for the multiple comparisons, we only kept the maximum counts among all the connections at each iteration and constructed a null distribution using these maxima. We considered FCs as important contributors to the SCZ or MDD diagnosis if their *P*-values were below 0.05. The list of discriminative FCs and their spatial distributions are provided in fig. S4 and table S13, while those for MDD are in fig. S5 and table S14.

#### Supplementary Figures

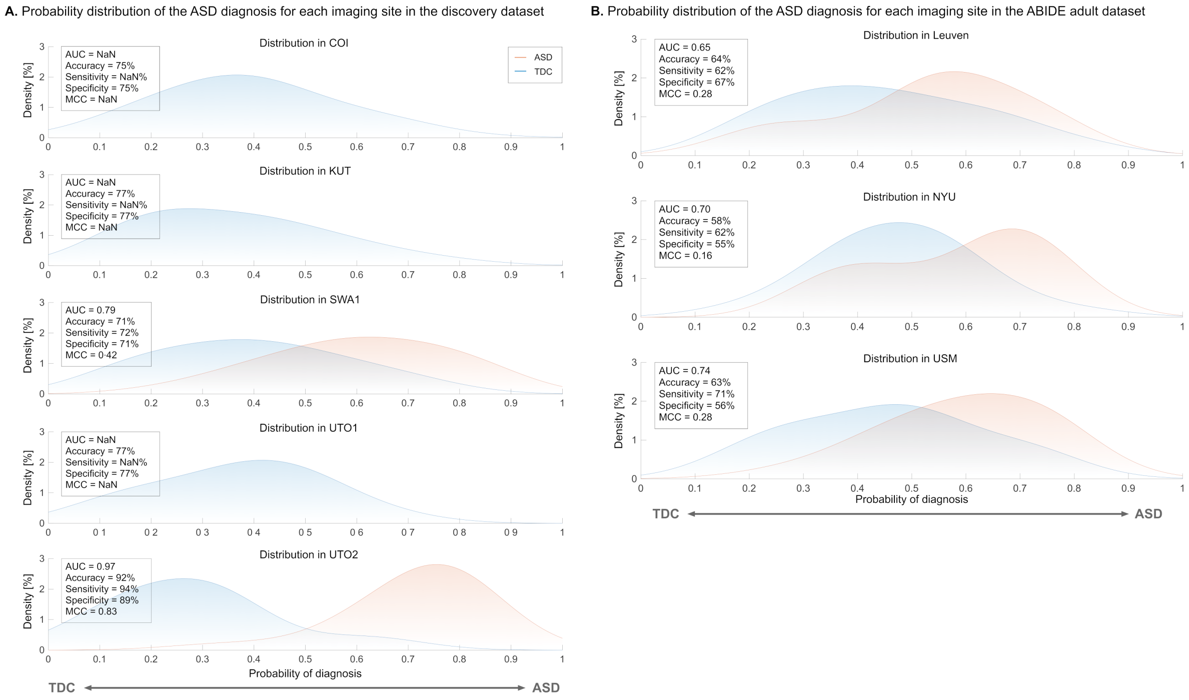


**Fig. S1. The distribution of the ASD diagnosis probability in each imaging site in the discovery dataset and the ABIDE adult validation dataset.** (A) In the discovery dataset, we visualized the probability distributions in each imaging site. (B) In the ABIDE adult dataset, we visualized the probability distributions in each imaging site. **Abbreviations:** AUC: area under the curve, ASD: autism spectrum disorder, MCC: Matthews correlation coefficient, and TDC: typically developing control.

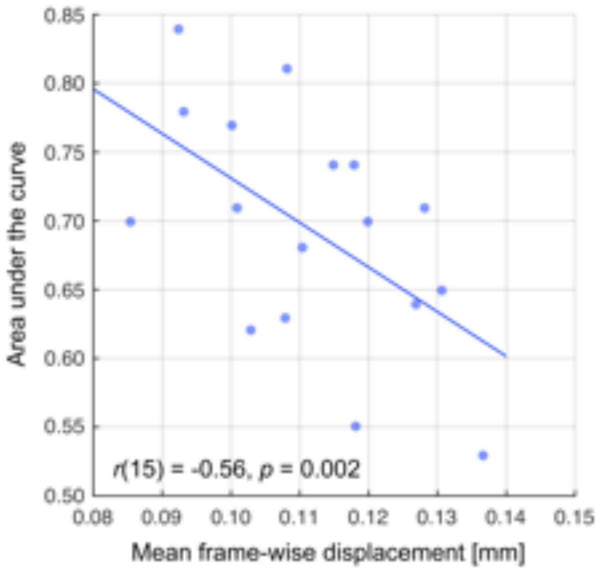


**Fig. S2. The relation between the head motion and performance index across the validation datasets.** We computed the area under the curve (AUC) and the mean frame-wise displacement (FD) in every imaging site. We then computed the Pearson correlation coefficient between the AUC and mean FD across imaging sites to investigate whether the head motion improved the classification performance.

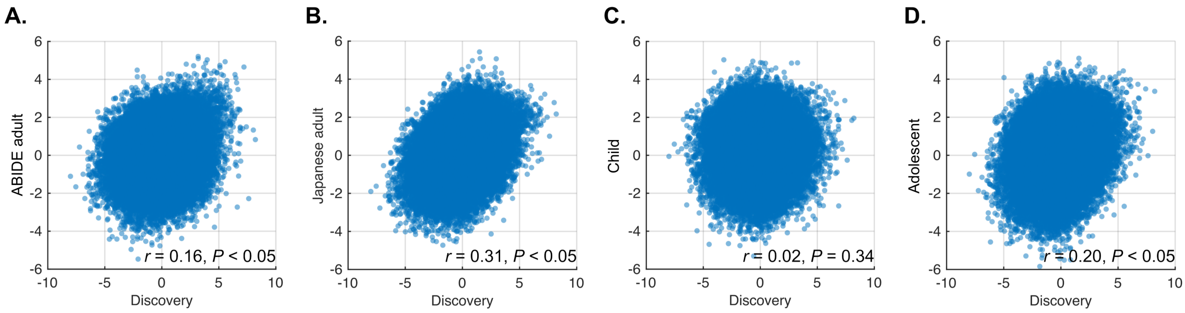


**Fig. S3. Results of mass-univariate analyses between the discovery and validation datasets.** (A) The relationship between the discovery dataset and the ABIDE adult dataset. (B) The relationship between the discovery dataset and the Japanese adult dataset. (C) The relationship between the discovery dataset and the child dataset. (D) The relationship between the discovery dataset and the adolescent dataset. **Abbreviations:** ABIDE: autism brain imaging data exchange.

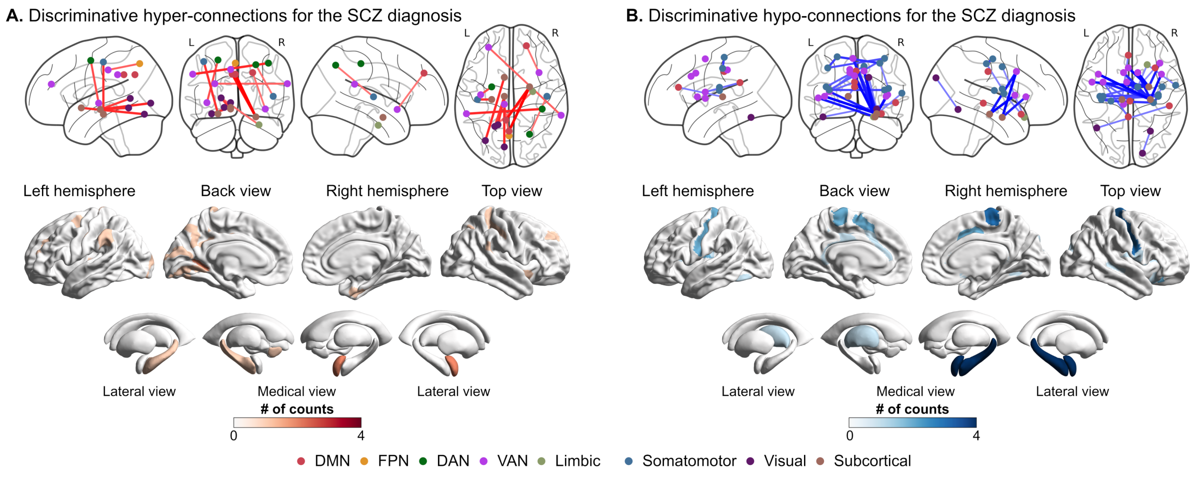


**Fig. S4. Discriminative functional connections for the SCZ diagnosis.** (A) The spatial distribution of hyper-connections for the diagnosis of schizophrenia (SCZ). (B) The spatial distribution of hypo-connections or the diagnosis of SCZ.

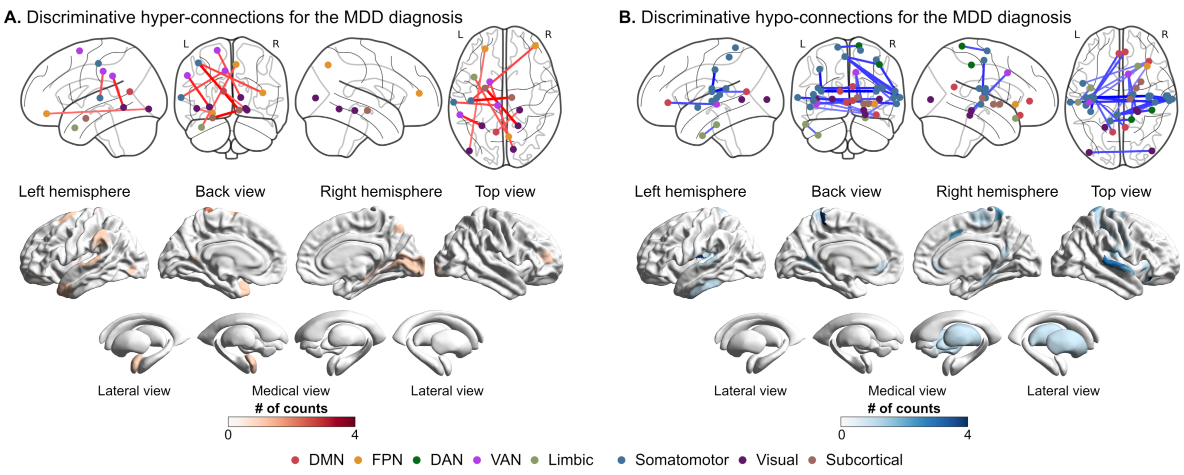


**Fig. S5. Discriminative functional connections for the MDD diagnosis.** (A) The spatial distribution of hyper-connections for the diagnosis of major depressive disorder (MDD). (B) The spatial distribution of hypo-connections or the diagnosis of MDD.

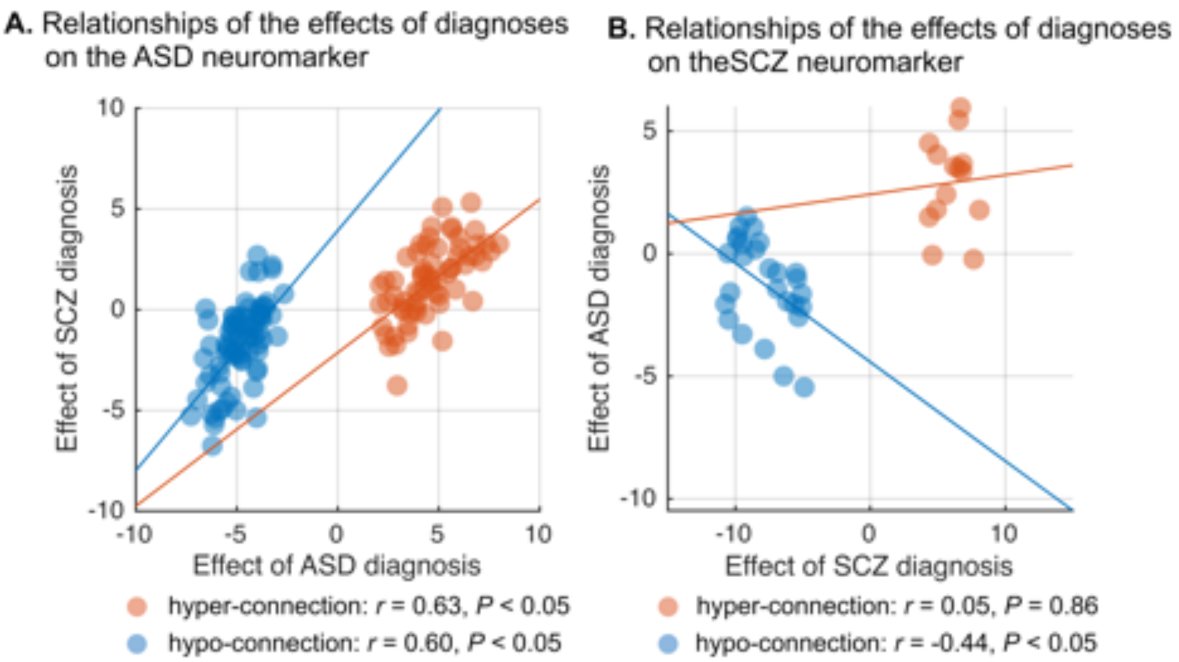


**Fig. S6. Relationship of the ASD and SCZ diagnosis on ASD and SCZ neuromarkers.** (A)The relationship of both diagnoses on ASD neuromarker. (B) The relationship of both diagnoses on SCZ neuromarker. **Abbreviations:** ASD: autism spectrum disorder, and SCZ: schizophrenia.

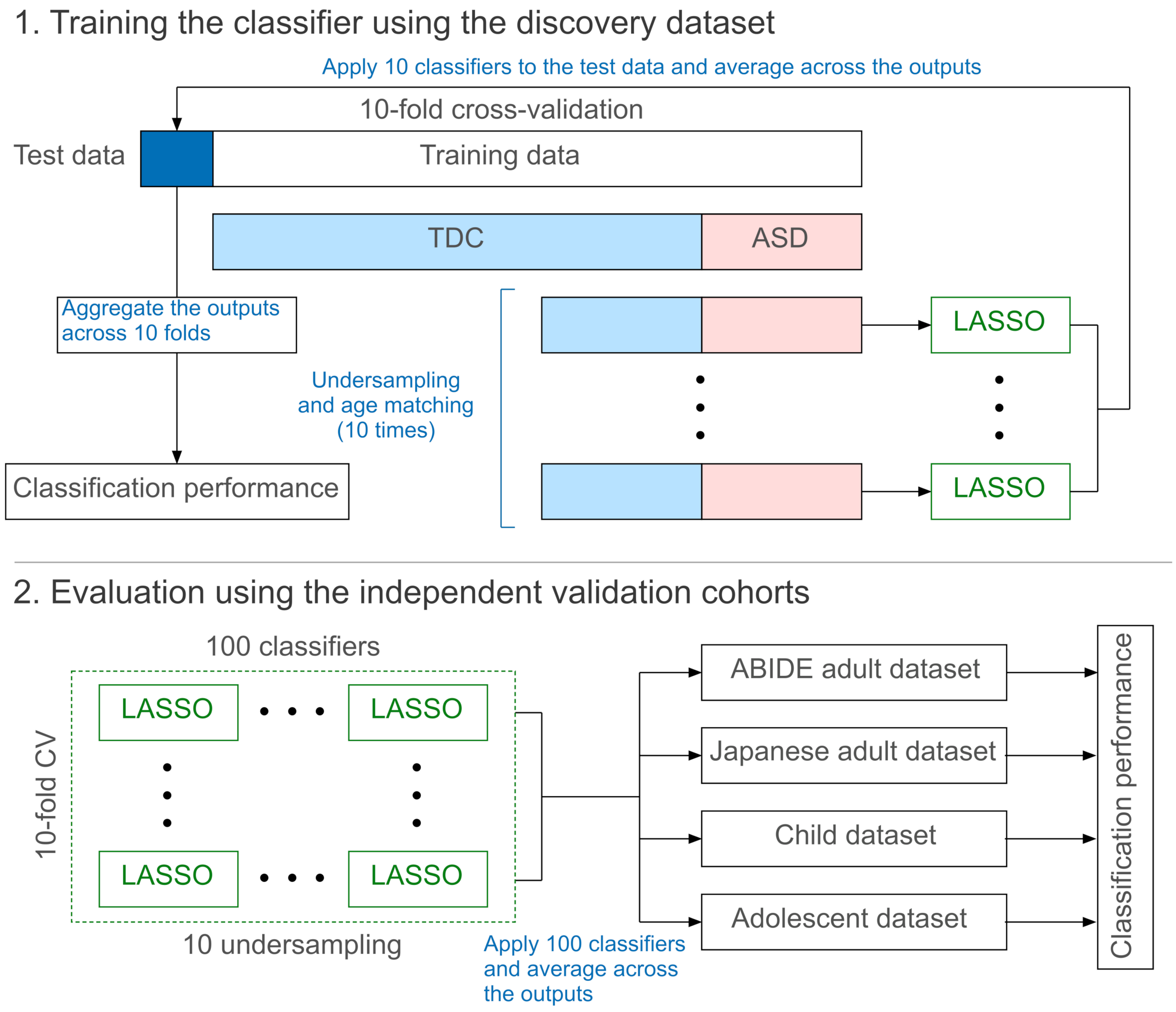


**Fig. S7. Schematic representation of training and evaluation procedures of the ASD neuromarker.** We built a neuromarker for autism spectrum disorder (ASD) using a 10-fold nested cross-validation (CV) procedure with ten undersamplings. The neuromarker's generalizability was evaluated by applying 100 trained classifiers to the validation datasets. **Abbreviations:** ABIDE: autism brain imaging data exchange, ASD: autism spectrum disorder, and TDC: typically developing control.

**Table S1.** Demographic information of the discovery and adult validation datasets.

| | TDC | | | ASD | | |
|---|---|---|---|---|---|---|
| | **N (M/F)** | **Age** | **Mean FD (SD) [mm]** | **N (M/F)** | **Age** | **Mean FD (SD) [mm]** |
| | | | **Discovery cohort** | | | |
| COI | 106 (41/65) | 50.75 (13.02) | 0.18 (0.05) | - | - | - |
| KUT | 148 (85/63) | 36.64 (13.41) | 0.13 (0.04) | - | - | - |
| SWA1 | 174 (151/23) | 30.94 (7.64) | 0.13 (0.04) | 145 (122/23) | 31.37 (7.78) | 0.14 (0.06) |
| UTO1 | 86 (29/57) | 44.45 (13.82) | 0.11 (0.04) | - | - | - |
| UTO2 | 36 (18/18) | 35.5 (7.57) | 0.16 (0.06) | 35 (23/12) | 31.94 (8.78) | 0.15 (0.04) |
| **Total** | 550 (324/226) | 38.70 (13.65) | 0.14 (0.05) | 180 (145/35) | 31.48 (7.96) | 0.14 (0.06) |
| | | | **ABIDE adult validation dataset** | | | |
| Leuven | 15 (15/0) | 23.27 (2.91) | 0.13 (0.04) | 13 (13/0) | 21.31 (3.71) | 0.13 (0.03) |
| NYU | 20 (15/5) | 23.81 (3.78) | 0.08 (0.02) | 13 (9/4) | 24.17 (4.44) | 0.08 (0.03) |
| USM | 32 (29/3) | 25.79 (6.09) | 0.11 (0.04) | 28 (26/2) | 25.69 (6.37) | 0.12 (0.04) |
| **Total** | 67 (59/8) | 24.63 (4.96) | 0.11 (0.04) | 54 (48/6) | 24.27 (5.61) | 0.11 (0.04) |
| | | | **Japanese adult validation dataset** | | | |
| SWA2 | 38 (27/11) | 28.5 (6.83) | 0.10 (0.03) | 22 (19/3) | 25.77 (5.23) | 0.12 (0.05) |

**Abbreviations:** ABIDE: autism brain imaging data exchange, ASD: autism spectrum disorder, COI: Center of Innovation, F: female, FD: frame-wise displacement, M: male, NYU: New York University, KUT: Kyoto University TimTrio, SD: standard deviation, SWA: Showa University, TDC: typically developing control, UTO: University of Tokyo, and USM: University of Utah School of Medicine

*The discovery dataset was matched for the mean FD ($p > 0.05$), except for age and sex ($p < 0.05$).

**The US adult validation cohort was matched for age, sex, and mean FD (p > 0.05).

***The Japanese adult validation cohort was matched for age and sex ($p > 0.05$) , but not matched for mean FD ($p < 0.05$)

**Table S2.** The classification performance of the ASD classifier in the discovery and adult validation datasets.

| | AUC | Accuracy [%] | Sensitivity [%] | Specificity [%] | MCC | PPV | NPV |
|---|---|---|---|---|---|---|---|
| **Classification performance on the discovery dataset** | | | | | | | |
| **All** | **0.84** | **75.62** | **76.11** | **75.45** | **0.46** | **0.50** | **0.91** |
| COI | - | 75.47 | - | 75.47 | - | - | - |
| KUT | - | 77.03 | - | 77.03 | - | - | - |
| SWA1 | 0.79 | 71.16 | 71.72 | 70.69 | 0.42 | 0.67 | 0.75 |
| UTO1 | - | 76.74 | - | 76.74 | - | - | - |
| UTO2 | 0.97 | 91.55 | 94.29 | 88.89 | 0.83 | 0.89 | 0.94 |
| **Generalizability of ASD classifier to the ABIDE adult dataset** | | | | | | | |
| **All** | **0.70** | **61.98** | **66.67** | **58.21** | **0.25** | **0.56** | **0.68** |
| Leuven | 0.65 | 64.29 | 61.54 | 66.67 | 0.28 | 0.62 | 0.66 |
| NYU | 0.70 | 57.59 | 61.54 | 55.00 | 0.16 | 0.47 | 0.69 |
| USM | 0.74 | 63.33 | 71.43 | 56.25 | 0.29 | 0.59 | 0.69 |
| **Generalizability of ASD classifier to the Japanese adult dataset** | | | | | | | |
| **SWA2** | **0.81** | **78.33** | **63.64** | **86.84** | **0.52** | **0.74** | **0.80** |

**Abbreviations:** ABIDE: autism brain imaging data exchange, AUC: area under the curve, COI: Center of Innovation, NYU: New York University, KUT: Kyoto University TimTrio, MCC: Matthews correlation coefficient, SWA: Showa University, UTO: University of Tokyo, and USM: University of Utah School of Medicine

**Table S3.** Demographic information for the child and adolescent validation datasets.

| | TDC | | | ASD | | |
|---|---|---|---|---|---|---|
| | **N (M/F)** | **Age** | **Mean FD (SD) [mm]** | **N (M/F)** | **Age** | **Mean FD (SD) [mm]** |
| **Child dataset** | | | | | | |
| GU | 13 (6/7) | 9.46 (1.08) | 0.12 (0.03) | 19 (16/3) | 10.36 (1.12) | 0.12 (0.03) |
| OHSU | 35 (16/19) | 9.60 (0.98) | 0.10 (0.03) | 11 (8/3) | 9.64 (1.43) | 0.12 (0.04) |
| KKI | 105 (61/44) | 10.13 (1.01) | 0.12 (0.04) | 32 (24/8) | 9.81 (1.23) | 0.14 (0.03) |
| NYU | 33 (28/5) | 9.09 (1.62) | 0.10 (0.03) | 38 (34/4) | 8.81 (1.80) | 0.11 (0.04) |
| UCLA | 16 (12/4) | 10.30 (1.43) | 0.10 (0.04) | 19 (17/2) | 10.69 (0.97) | 0.12 (0.04) |
| **Total** | 202 (123/79) | 9.84 (1.22) | 0.11 (0.04) | 119 (99/20) | 9.70 (1.55) | 0.12 (0.04) |
| **Adolescent dataset** | | | | | | |
| OHSU | 12 (8/4) | 12.58 (0.79) | 0.10 (0.03) | 18 (15/3) | 13.39 (1.09) | 0.12 (0.03) |
| SDSU | 10 (9/1) | 15.15 (1.97) | 0.09 (0.04) | 12 (11/1) | 15.37 (1.95) | 0.09 (0.03) |
| NYU | 33 (24/9) | 14.65 (1.49) | 0.09 (0.04) | 13 (10/3) | 14.19 (1.46) | 0.11 (0.03) |
| TRINITY | 13 (13/0) | 14.81 (1.60) | 0.13 (0.02) | 12 (12/0) | 15.19 (1.37) | 0.14 (0.03) |
| UCLA | 30 (24/6) | 13.79 (1.47) | 0.09 (0.02) | 26 (24/2) | 14.73 (1.71) | 0.10 (0.03) |
| UM | 10 (7/3) | 15.82 (1.75) | 0.10 (0.03) | 12 (10/2) | 15.34 (1.55) | 0.12 (0.05) |
| USM | 11 (11/0) | 14.81 (1.50) | 0.12 (0.03) | 18 (18/0) | 16.30 (1.27) | 0.13 (0.04) |
| YALE | 13 (8/5) | 14.85 (1.64) | 0.11 (0.04) | 10 (7/3) | 14.54 (1.92) | 0.13 (0.03) |
| **Total** | 132 (104/28) | 14.44 (1.69) | 0.10 (0.03) | 121 (107/14) | 14.86 (1.73) | 0.12 (0.04) |

**Abbreviations:** ASD: autism spectrum disorder, GU: Georgetown University, KKI: Kennedy Krieger Institute, NYU: New York UniversityOHSU: Oregon Health and Science University, SD: standard deviation, SDSU: San Diego State University, TDC: typically developing control, TRINITY: Trinity Centre for Health Sciences, UCLA: the University of California, Los Angeles, UM: University of Michigan, USM: University of Utah School of Medicine, and YALE: Yale Child Study Center.

*The child cohort was matched for age ($p > 0.05$), but not matched for sex and mean FD ($p < 0.05$).

**The adolescent cohort was matched for age and sex ($p > 0.05$), but not matched for mean FD ($p < 0.05$).

**Table S4.** The classification performance of the ASD classifier in the child and adolescent datasets.

| | AUC | Accuracy [%] | Sensitivity [%] | Specificity [%] | MCC | PPV | NPV |
|---|---|---|---|---|---|---|---|
| **Generalizability of ASD classifier to the child dataset** | | | | | | | |
| **All** | **0.66** | **60.75** | **75.63** | **51.98** | **0.27** | **0.48** | **0.78** |
| GU | 0.70 | 68.75 | 84.21 | 46.15 | 0.33 | 0.70 | 0.67 |
| OHSU | 0.71 | 65.23 | 72.73 | 62.86 | 0.30 | 0.38 | 0.88 |
| KKI | 0.64 | 56.20 | 75.00 | 50.48 | 0.22 | 0.32 | 0.87 |
| NYU | 0.62 | 59.15 | 71.05 | 45.45 | 0.17 | 0.60 | 0.58 |
| UCLA | 0.68 | 68.57 | 78.95 | 56.25 | 0.36 | 0.68 | 0.69 |
| **Generalizability of ASD classifier to the adolescent dataset** | | | | | | | |
| **All** | **0.71** | **65.61** | **71.07** | **60.61** | **0.32** | **0.62** | **0.70** |
| OHSU | 0.55 | 46.67 | 55.56 | 33.33 | -0.112 | 0.56 | 0.33 |
| SDSU | 0.84 | 68.18 | 66.67 | 70.00 | 0.37 | 0.73 | 0.64 |
| NYU | 0.77 | 67.39 | 76.92 | 63.64 | 0.37 | 0.45 | 0.88 |
| TRINITY | 0.53 | 56.00 | 50.00 | 61.54 | 0.12 | 0.55 | 0.57 |
| UCLA | 0.78 | 69.64 | 76.92 | 63.33 | 0.40 | 0.65 | 0.76 |
| UM | 0.63 | 72.73 | 91.67 | 50.00 | 0.47 | 0.69 | 0.83 |
| USM | 0.71 | 68.97 | 77.78 | 54.55 | 0.33 | 0.74 | 0.60 |
| YALE | 0.74 | 73.91 | 70.00 | 76.92 | 0.47 | 0.70 | 0.77 |

**Abbreviations:** ASD: autism spectrum disorder, GU: Georgetown University, KKI: Kennedy Krieger Institute, NPV: negative predictive value, NYU: New York UniversityOHSU: Oregon Health and Science University, PPV: positive predictive value, SD: standard deviation, SDSU: San Diego State University, TDC: typically developing control, TRINITY: Trinity Centre for Health Sciences, UCLA: the University of California, Los Angeles, UM: University of Michigan, USM: University of Utah School of Medicine, and YALE: Yale Child Study Center.

**Table S5.** The classification performance of the ASD classifier on the discovery dataset without ComBat harmonization method.

| | AUC | Accuracy [%] | Sensitivity [%] | Specificity [%] | MCC |
|---|---|---|---|---|---|
| **Classification performance on the discovery dataset without harmonization** | | | | | |
| **All** | **0.85** | **76.71** | **80.56** | **75.45** | **0.50** |
| COI | - | 80.19 | - | 80.19 | - |
| KUT | - | 89.19 | - | 89.19 | - |
| SWA1 | 0.75 | 66.77 | 0.793103 | 56.32 | 0.36 |
| UTO1 | - | 79.07 | - | 79.07 | - |
| UTO2 | 0.90 | 87.32 | 0.857143 | 88.89 | 0.75 |
| **Generalizability of ASD classifier to the ABIDE adult dataset** | | | | | |
| **All** | **0.64** | **59.50** | **59.23** | **59.70** | **0.19** |
| Leuven | 0.60 | 57.14 | 38.46 | 73.33 | 0.13 |
| NYU | 0.62 | 51.52 | 69.23 | 40.00 | 0.09 |
| USM | 0.70 | 65.00 | 64.29 | 65.63 | 0.30 |
| **Generalizability of ASD classifier to the Japanese adult dataset** | | | | | |
| **SWA2** | **0.73** | **66.67** | **36.36** | **84.21** | **0.23** |

**Abbreviations:** AUC: area under the curve, COI: Center of Innovation, NYU: New York University, KUT: Kyoto University TimTrio, MCC: Matthews correlation coefficient, SWA: Showa University, UTO: University of Tokyo, and USM: University of Utah School of Medicine

**Table S6.** The classification performance of the ASD classifier trained on SWA1 and UTO2 only.

| | AUC | Accuracy [%] | Sensitivity [%] | Specificity [%] | MCC |
|---|---|---|---|---|---|
| **Classification performance on the balanced discovery dataset** | | | | | |
| **All** | **0.80** | **73.85** | **72.22** | **75.24** | **0.47** |
| SWA1 | 0.76 | 71.16 | 68.28 | 73.56 | 0.42 |
| UTO2 | 0.96 | 85.92 | 88.57 | 83.33 | 0.72 |
| **Generalizability of ASD classifier to the ABIDE adult dataset** | | | | | |
| **All** | **0.67** | **57.85** | **77.78** | **41.79** | **0.21** |
| Leuven | 0.56 | 57.14 | 69.23 | 46.67 | 0.16 |
| NYU | 0.60 | 48.48 | 76.92 | 30.00 | 0.08 |
| USM | 0.72 | 63.33 | 82.14 | 46.88 | 0.31 |
| **Generalizability of ASD classifier to the Japanese adult dataset** | | | | | |
| **SWA2** | **0.83** | **71.67** | **72.73** | **71.05** | **0.42** |

**Abbreviations:** AUC: area under the curve, COI: Center of Innovation, NYU: New York University, KUT: Kyoto University TimTrio, MCC: Matthews correlation coefficient, SWA: Showa University, UTO: University of Tokyo, and USM: University of Utah School of Medicine

**Table S7**. The classification performance of the ASD classifier trained on the SRPBS dataset only.

| | AUC | Accuracy [%] | Sensitivity [%] | Specificity [%] | MCC |
|---|---|---|---|---|---|
| **Classification performance on the SRPBS dataset** | | | | | |
| **All** | **0.84** | **76.33** | **77.93** | **75.88** | **0.46** |
| COI | - | 77.36 | - | 77.36 | - |
| KUT | - | 78.38 | - | 78.38 | - |
| SWA1 | 0.82 | 75.24 | 77.93 | 72.99 | 0.51 |
| UTO | - | 75.58 | - | 75.58 | - |
| **Generalizability of ASD classifier to the ABIDE adult dataset** | | | | | |
| **All** | **0.70** | **62.81** | **59.26** | **65.67** | **0.25** |
| Leuven | 0.65 | 60.71 | 53.85 | 66.67 | 0.21 |
| NYU | 0.68 | 60.61 | 53.85 | 65.00 | 0.19 |
| USM | 0.74 | 65.00 | 64.29 | 65.63 | 0.30 |
| **Generalizability of ASD classifier to the Japanese adult dataset** | | | | | |
| **SWA2** | **0.82** | **73.33** | **63.64** | **78.95** | **0.43** |

**Abbreviations:** AUC: area under the curve, COI: Center of Innovation, NYU: New York University, KUT: Kyoto University TimTrio, MCC: Matthews correlation coefficient, SWA: Showa University, UTO: University of Tokyo, and USM: University of Utah School of Medicine

**Table S8.** Comparisons of classification and generalization performance between higher/lower resolutions of regions of interest.

| Atlas | **AUC** | **Accuracy [%]** | **Sensitivity [%]** | **Specificity [%]** | **MCC** |
|---|---|---|---|---|---|
| **Classification performance on the discovery dataset** | | | | | |
| **Glasser** | 0.84 | **75.62** | 76.11 | **75.45** | 0.46 |
| Schaefer 100 | 0.81 | 71.91 | 72.78 | 71.64 | 0.39 |
| Schaefer 200 | **0.85** | 75.48 | **80.56** | 73.82 | **0.48** |
| Schaefer 300 | 0.82 | 73.70 | 75.56 | 73.09 | 0.43 |
| Schaefer 400 | 0.83 | 74.52 | 78.89 | 73.09 | 0.46 |
| Schaefer 500 | 0.82 | 74.52 | 76.67 | 73.82 | 0.45 |
| Schaefer 600 | 0.83 | 74.11 | 77.22 | 73.09 | 0.44 |
| Schaefer 700 | 0.83 | 74.38 | 77.78 | 73.27 | 0.45 |
| Schaefer 800 | 0.83 | 73.42 | 76.67 | 72.36 | 0.43 |
| Schaefer 900 | 0.83 | 73.42 | 75.56 | 72.73 | 0.43 |
| **Generalizability on the ABIDE adult dataset** | | | | | |
| **Glasser** | 0.70 | 61.98 | 66.67 | 58.21 | 0.25 |
| Schaefer 100 | 0.70 | 65.29 | 64.81 | 65.67 | 0.30 |
| Schaefer 200 | 0.73 | 65.29 | 59.26 | 70.15 | 0.30 |
| Schaefer 300 | 0.70 | 65.29 | 57.41 | **71.64** | 0.29 |
| Schaefer 400 | 0.72 | 66.12 | 59.26 | **71.64** | 0.31 |
| Schaefer 500 | **0.75** | **71.07** | 74.07 | 68.66 | 0.42 |
| Schaefer 600 | 0.73 | 69.42 | 68.52 | 70.15 | 0.39 |
| Schaefer 700 | 0.72 | 67.77 | 74.07 | 62.69 | 0.37 |
| Schaefer 800 | 0.72 | 66.94 | 64.81 | 68.66 | 0.33 |
| Schaefer 900 | 0.72 | **71.07** | **77.78** | 65.67 | **0.43** |
| **Generalizability on the Japanese adult dataset** | | | | | |
| **Glasser** | 0.81 | **78.33** | 63.64 | **86.84** | 0.52 |
| Schaefer 100 | 0.84 | 76.67 | **86.36** | 71.05 | **0.55** |
| Schaefer 200 | 0.83 | 73.33 | 68.18 | 76.32 | 0.44 |
| Schaefer 300 | 0.84 | 71.67 | 72.73 | 71.05 | 0.42 |
| Schaefer 400 | 0.84 | 76.67 | 72.73 | 78.95 | 0.51 |
| Schaefer 500 | 0.84 | 76.67 | 72.73 | 78.95 | 0.51 |
| Schaefer 600 | 0.83 | 73.33 | 63.64 | 78.95 | 0.43 |
| Schaefer 700 | 0.82 | 70.00 | 68.18 | 71.05 | 0.38 |
| Schaefer 800 | 0.82 | 73.33 | 72.73 | 73.68 | 0.45 |
| Schaefer 900 | **0.85** | 71.67 | 77.27 | 68.42 | 0.44 |

**Abbreviations:** ABIDE: autism brain imaging data exchange, AUC: area under the curve, and MCC: Matthews correlation coefficient.

**Table S9.** The list of discriminative FCs for the ASD diagnosis.

| ROI 1 | | | ROI 2 | | | $z$(ASD) | $z$(TDC) | $t$-value | Mean weight |
|---|---|---|---|---|---|---|---|---|---|
| **Glasser's label** | **AAL label** | **Network** | **Glasser's label** | **AAL label** | **Network** | | | | |
| R.Amy | Hippocampus_R | Subcortical | R.A5 | Temporal_Sup_R | Somatomotor | 0.12 | 0.01 | 8.03 | 1.51 |
| R.PBelt | Temporal_Sup_R | Somatomotor | L.POS2 | Precuneus_L | FPN | 0.03 | -0.08 | 7.42 | 0.48 |
| R.25 | Olfactory_R | Limbic | L.PHT | Temporal_Mid_L | DAN | 0.01 | -0.05 | 4.32 | 0.41 |
| R.MBelt | Heschl_R | Somatomotor | R.LIPv | Parietal_Inf_R | DAN | -0.02 | -0.08 | 4.51 | 0.41 |
| R.A5 | Temporal_Sup_R | Somatomotor | R.23d | Cingulum_Mid_R | DMN | 0.12 | 0.01 | 6.73 | 0.35 |
| R.PBelt | Temporal_Sup_R | Somatomotor | L.IFJp | Frontal_Inf_Oper_L | DAN | 0.08 | 0.03 | 3.90 | 0.26 |
| R.p24 | Cingulum_Ant_R | FPN | R.A5 | Temporal_Sup_R | Somatomotor | 0.12 | 0.01 | 7.62 | 0.26 |
| R.MidB | - | Subcortical | R.TGd | Temporal_Pole_Mid_R | Limbic | 0.03 | -0.06 | 5.70 | 0.26 |
| R.VMV2 | Lingual_R | Visual | L.STGa | Temporal_Pole_Sup_L | Limbic | 0.03 | -0.01 | 3.53 | 0.25 |
| L.TGd | Temporal_Pole_Mid_L | Limbic | L.FOP1 | Rolandic_Oper_L | VAN | -0.10 | -0.15 | 3.28 | 0.24 |
| R.Amy | Hippocampus_R | Subcortical | L.TGd | Temporal_Pole_Mid_L | Limbic | 0.01 | -0.04 | 4.22 | 0.21 |
| R.Amy | Hippocampus_R | Subcortical | R.6d | Precentral_R | Somatomotor | 0.07 | -0.04 | 7.20 | 0.16 |
| R.PCV | Precuneus_R | FPN | L.a9-46v | Frontal_Mid_L | FPN | -0.01 | -0.09 | 5.00 | 0.16 |
| R.Pallidum | Pallidum_R | Subcortical | L.MST | Occipital_Mid_L | Visual | 0.02 | -0.05 | 5.97 | 0.15 |
| R.s32 | Frontal_Med_Orb_R | DMN | L.MIP | Parietal_Sup_L | DAN | -0.06 | -0.11 | 3.65 | 0.15 |

| | | | | | | | | | |
|---|---|---|---|---|---|---|---|---|---|
| R.6v | Precentral_R | Somatomotor | L.MI | Insula_L | VAN | 0.21 | 0.12 | 6.09 | 0.14 |
| R.s32 | Frontal_Med_Orb_R | DMN | L.V6 | Cuneus_L | Visual | -0.07 | -0.11 | 2.42 | 0.14 |
| L.MidB | - | Subcortical | R.STSda | Temporal_Sup_R | DMN | 0.15 | 0.04 | 6.85 | 0.14 |
| R.SFL | Supp_Motor_Area_R | DMN | R.V3A | Occipital_Sup_R | Visual | 0.00 | -0.09 | 6.56 | 0.13 |
| L.MidB | - | Subcortical | R.TPOJ1 | Temporal_Mid_R | VAN | 0.01 | -0.06 | 5.68 | 0.12 |
| R.p47r | Frontal_Inf_Tri_R | FPN | L.TPOJ2 | Temporal_Mid_L | DAN | -0.07 | -0.13 | 3.51 | 0.12 |
| R.VIP | Parietal_Sup_R | DAN | L.FEF | Precentral_L | DAN | 0.28 | 0.20 | 4.43 | 0.11 |
| L.FFC | Fusiform_L | Visual | L.RSC | - | DMN | -0.08 | -0.14 | 4.59 | 0.11 |
| R.VMV2 | Lingual_R | Visual | R.STGa | Temporal_Pole_Sup_R | DMN | 0.02 | -0.01 | 2.87 | 0.11 |
| R.p24 | Cingulum_Ant_R | FPN | R.A4 | Temporal_Sup_R | Somatomotor | 0.10 | 0.02 | 5.49 | 0.11 |
| L.OFC | Rectus_L | Limbic | L.6r | Frontal_Inf_Oper_L | VAN | -0.03 | -0.07 | 2.92 | 0.10 |
| R.HC | Hippocampus_R | Subcortical | L.A5 | Temporal_Mid_L | DMN | 0.12 | 0.04 | 6.02 | 0.10 |
| R.PGp | Occipital_Mid_R | DAN | L.A1 | Rolandic_Oper_L | Somatomotor | -0.01 | -0.08 | 5.06 | 0.10 |
| R.LIPv | Parietal_Inf_R | DAN | L.6v | Precentral_L | Somatomotor | 0.28 | 0.20 | 4.89 | 0.10 |
| R.SFL | Supp_Motor_Area_R | DMN | L.V3A | Occipital_Sup_L | Visual | 0.02 | -0.07 | 6.92 | 0.10 |
| L.V7 | Occipital_Mid_L | Visual | L.RSC | - | DMN | -0.04 | -0.12 | 5.23 | 0.10 |
| R.6v | Precentral_R | Somatomotor | L.5mv | Cingulum_Mid_L | VAN | 0.15 | 0.06 | 5.64 | 0.10 |
| R.47m | Frontal_Inf_Orb_R | DMN | R.v23ab | Precuneus_R | DMN | 0.27 | 0.16 | 5.67 | 0.10 |
| R.LBelt | Temporal_Sup_R | Somatomotor | R.7Pm | Precuneus_R | FPN | -0.04 | -0.09 | 4.28 | 0.09 |
| R.V3CD | Occipital_Mid_R | Visual | L.MBelt | Temporal_Sup_L | Somatomotor | -0.02 | -0.08 | 4.63 | 0.09 |

| | | | | | | | | | |
|---|---|---|---|---|---|---|---|---|---|
| L.Pallidum | Pallidum_L | Subcortical | L.TE2a | Temporal_Inf_L | Limbic | 0.00 | -0.06 | 3.34 | 0.09 |
| R.VMV2 | Lingual_R | Visual | R.TGd | Temporal_Pole_Mid_R | Limbic | 0.00 | -0.04 | 2.51 | 0.09 |
| R.6d | Precentral_R | Somatomotor | L.FEF | Precentral_L | DAN | 0.19 | 0.12 | 3.47 | 0.08 |
| R.PFm | Parietal_Inf_R | DMN | R.d32 | Cingulum_Ant_R | DMN | 0.38 | 0.29 | 5.23 | 0.08 |
| R.PCV | Precuneus_R | FPN | L.AVI | Insula_L | FPN | 0.00 | -0.05 | 3.99 | 0.08 |
| R.MidB | - | Subcortical | R.PBelt | Temporal_Sup_R | Somatomotor | 0.17 | 0.06 | 6.65 | 0.08 |
| L.MidB | - | Subcortical | L.PHA2 | ParaHippocampal_L | DMN | 0.15 | 0.08 | 4.99 | 0.08 |
| R.Putamen | Putamen_R | Subcortical | R.A5 | Temporal_Sup_R | Somatomotor | 0.05 | -0.03 | 6.36 | 0.08 |
| R.MidB | - | Subcortical | R.TE1a | Temporal_Mid_R | DMN | -0.02 | -0.09 | 4.71 | 0.08 |
| L.A5 | Temporal_Mid_L | DMN | L.9m | Frontal_Sup_Medial_L | DMN | 0.26 | 0.16 | 5.88 | 0.07 |
| R.IP1 | Angular_R | DAN | R.PGp | Occipital_Mid_R | DAN | 0.26 | 0.16 | 4.63 | 0.07 |
| R.Amy | Hippocampus_R | Subcortical | R.TGd | Temporal_Pole_Mid_R | Limbic | -0.01 | -0.07 | 4.23 | 0.07 |
| R.PCV | Precuneus_R | FPN | L.a32pr | Cingulum_Ant_L | FPN | 0.10 | 0.03 | 4.15 | 0.07 |
| R.6ma | Frontal_Sup_R | DAN | L.PreS | ParaHippocampal_L | DMN | -0.05 | -0.08 | 2.14 | 0.07 |
| R.25 | Olfactory_R | Limbic | R.PHT | Temporal_Inf_R | DAN | 0.01 | -0.04 | 3.90 | 0.06 |
| R.s32 | Frontal_Med_Orb_R | DMN | L.7AL | Parietal_Sup_L | DAN | -0.06 | -0.10 | 2.59 | 0.06 |
| R.PHT | Temporal_Inf_R | DAN | L.a24 | Cingulum_Ant_L | DMN | -0.13 | -0.16 | 2.25 | 0.06 |
| R.SFL | Supp_Motor_Area_R | DMN | L.9-46d | Frontal_Mid_L | VAN | 0.14 | 0.05 | 4.71 | 0.06 |
| R.IPS1 | Occipital_Sup_R | DAN | L.6v | Precentral_L | Somatomotor | 0.20 | 0.12 | 4.39 | 0.06 |
| R.HC | Hippocampus_R | Subcortical | L.V7 | Occipital_Mid_L | Visual | -0.02 | -0.08 | 5.10 | 0.06 |
| R.25 | Olfactory_R | Limbic | R.IPS1 | Occipital_Sup_R | DAN | -0.04 | -0.09 | 3.94 | 0.06 |
| R.s6-8 | Frontal_Sup_R | FPN | L.a9-46v | Frontal_Mid_L | FPN | 0.09 | 0.04 | 2.40 | 0.06 |

| | | | | | | | | | |
|---|---|---|---|---|---|---|---|---|---|
| R.VIP | Parietal_Sup_R | DAN | L.LO2 | Occipital_Inf_L | Visual | 0.22 | 0.14 | 4.01 | 0.06 |
| R.7Pm | Precuneus_R | FPN | L.A1 | Rolandic_Oper_L | Somatomotor | -0.05 | -0.10 | 4.05 | 0.05 |
| R.6v | Precentral_R | Somatomotor | L.PF | SupraMarginal_L | VAN | 0.22 | 0.10 | 6.88 | 0.05 |
| L.OFC | Rectus_L | Limbic | L.IFSa | Frontal_Inf_Tri_L | FPN | 0.05 | 0.01 | 3.21 | 0.05 |
| R.7PL | Parietal_Sup_R | DAN | R.LO2 | Occipital_Inf_R | Visual | 0.09 | 0.04 | 2.85 | 0.05 |
| R.6mp | Supp_Motor_Area_R | Somatomotor | R.5L | Postcentral_R | DAN | 0.60 | 0.54 | 2.99 | 0.05 |
| R.p32pr | Cingulum_Mid_R | VAN | R.7Pm | Precuneus_R | FPN | -0.03 | -0.08 | 2.84 | 0.05 |
| L.p47r | Frontal_Inf_Tri_L | FPN | L.10d | Frontal_Sup_Medial_L | DMN | 0.10 | 0.05 | 2.16 | 0.05 |
| R.10d | Frontal_Sup_Medial_R | DMN | L.9p | Frontal_Sup_L | DMN | 0.25 | 0.34 | -3.93 | -0.03 |
| R.46 | Frontal_Mid_R | FPN | R.47m | Frontal_Inf_Orb_R | DMN | -0.08 | 0.01 | -5.19 | -0.03 |
| R.Ig | Insula_R | Somatomotor | R.FOP2 | Rolandic_Oper_R | Somatomotor | 0.47 | 0.54 | -3.97 | -0.04 |
| L.LBelt | Temporal_Sup_L | Somatomotor | L.Ig | Insula_L | Somatomotor | 0.24 | 0.30 | -3.88 | -0.04 |
| R.9m | Frontal_Sup_Medial_R | DMN | R.PCV | Precuneus_R | FPN | 0.00 | 0.11 | -5.49 | -0.04 |
| R.TPOJ1 | Temporal_Mid_R | VAN | R.4 | Precentral_R | Somatomotor | 0.08 | 0.17 | -5.65 | -0.04 |
| L.PFcm | Temporal_Sup_L | Somatomotor | L.PSL | Temporal_Sup_L | VAN | 0.40 | 0.45 | -2.62 | -0.05 |
| L.p10p | Frontal_Sup_L | DMN | L.25 | Olfactory_L | Limbic | -0.03 | 0.03 | -3.45 | -0.05 |
| R.47l | Frontal_Inf_Orb_R | DMN | L.47l | Frontal_Inf_Orb_L | DMN | 0.48 | 0.56 | -4.05 | -0.05 |
| L.VMV1 | Lingual_L | Visual | L.TPOJ3 | Occipital_Mid_L | DAN | 0.08 | 0.16 | -4.35 | -0.05 |
| R.Pir | Insula_R | Limbic | L.V3 | Occipital_Sup_L | Visual | -0.10 | -0.05 | -3.85 | -0.05 |
| R.TPOJ1 | Temporal_Mid_R | VAN | L.A5 | Temporal_Mid_L | DMN | 0.33 | 0.43 | -4.98 | -0.05 |

| | | | | | | | | | |
|---|---|---|---|---|---|---|---|---|---|
| R.p24 | Cingulum_Ant_R | FPN | L.9m | Frontal_Sup_Medial_L | DMN | 0.27 | 0.34 | -3.69 | -0.05 |
| R.TPOJ1 | Temporal_Mid_R | VAN | L.MT | Occipital_Mid_L | Visual | 0.05 | 0.16 | -5.72 | -0.06 |
| R.STSdp | Temporal_Sup_R | DMN | L.V4t | Occipital_Mid_L | Visual | -0.04 | 0.01 | -3.92 | -0.06 |
| R.MidB | - | Subcortical | R.46 | Frontal_Mid_R | FPN | -0.01 | 0.07 | -6.02 | -0.06 |
| R.8Ad | Frontal_Mid_R | DMN | R.FFC | Fusiform_R | Visual | -0.20 | -0.15 | -3.76 | -0.06 |
| R.5m | Paracentral_Lobule_R | Somatomotor | L.STV | Temporal_Sup_L | DMN | -0.03 | 0.04 | -4.74 | -0.06 |
| L.TE2p | Temporal_Inf_L | DAN | L.10d | Frontal_Sup_Medial_L | DMN | -0.18 | -0.12 | -3.69 | -0.06 |
| R.FFC | Fusiform_R | Visual | L.55b | Precentral_L | DAN | -0.07 | 0.00 | -4.34 | -0.06 |
| R.5L | Postcentral_R | DAN | L.STV | Temporal_Sup_L | DMN | -0.03 | 0.04 | -5.17 | -0.06 |
| R.9m | Frontal_Sup_Medial_R | DMN | L.SFL | Supp_Motor_Area_L | DMN | 0.24 | 0.30 | -3.21 | -0.06 |
| R.MidB | - | Subcortical | L.a9-46v | Frontal_Mid_L | FPN | 0.01 | 0.10 | -6.07 | -0.06 |
| R.Cereb | Cerebellum_R | Subcortical | L.44 | Frontal_Inf_Oper_L | DMN | -0.03 | 0.05 | -5.80 | -0.06 |
| L.Putamen | Putamen_L | Subcortical | R.A4 | Temporal_Sup_R | Somatomotor | -0.02 | 0.06 | -6.26 | -0.07 |
| R.p32 | Frontal_Sup_Medial_R | DMN | L.d23ab | Cingulum_Post_L | DMN | 0.16 | 0.22 | -3.66 | -0.07 |
| R.A4 | Temporal_Sup_R | Somatomotor | R.TE2p | Temporal_Inf_R | DAN | -0.05 | 0.04 | -6.49 | -0.07 |
| R.v23ab | Precuneus_R | DMN | L.v23ab | Precuneus_L | DMN | 0.96 | 1.05 | -4.49 | -0.07 |
| R.RSC | Cingulum_Post_R | DMN | L.RSC | - | DMN | 1.03 | 1.08 | -2.94 | -0.07 |
| R.PHA2 | ParaHippocampal_R | Visual | R.a24 | Cingulum_Ant_R | DMN | 0.09 | 0.14 | -3.22 | -0.07 |
| R.VMV1 | Lingual_R | Visual | L.PCV | Precuneus_L | DMN | -0.01 | 0.08 | -5.31 | -0.07 |
| L.A5 | Temporal_Mid_L | DMN | L.PIT | Fusiform_L | Visual | -0.08 | 0.00 | -5.49 | -0.07 |
| L.TPOJ3 | Occipital_Mid_L | DAN | L.ProS | Precuneus_L | Visual | 0.10 | 0.17 | -3.85 | -0.08 |
| R.TPOJ1 | Temporal_Mid_R | VAN | L.V4t | Occipital_Mid_L | Visual | 0.00 | 0.11 | -6.48 | -0.08 |

| | | | | | | | | | |
|---|---|---|---|---|---|---|---|---|---|
| R.VMV2 | Lingual_R | Visual | L.PCV | Precuneus_L | DMN | 0.04 | 0.12 | -4.36 | -0.08 |
| R.a10p | Frontal_Sup_Orb_R | FPN | L.47m | Frontal_Inf_Orb_L | DMN | -0.09 | 0.01 | -4.66 | -0.08 |
| R.MidB | - | Subcortical | B.Stem | - | Subcortical | 0.23 | 0.31 | -5.40 | -0.08 |
| R.TGv | Temporal_Inf_R | Limbic | R.TGd | Temporal_Pole_Mid_R | Limbic | 0.49 | 0.58 | -3.73 | -0.08 |
| R.8Av | Frontal_Mid_R | FPN | L.V3B | Occipital_Mid_L | Visual | -0.23 | -0.17 | -3.89 | -0.08 |
| R.25 | Olfactory_R | Limbic | R.SCEF | Supp_Motor_Area_R | VAN | -0.15 | -0.08 | -5.38 | -0.08 |
| R.VMV2 | Lingual_R | Visual | R.7Am | Precuneus_R | DAN | 0.08 | 0.14 | -3.79 | -0.08 |
| R.10pp | Frontal_Sup_Orb_R | Limbic | L.47m | Frontal_Inf_Orb_L | DMN | -0.09 | 0.01 | -4.50 | -0.09 |
| R.VMV2 | Lingual_R | Visual | R.DVT | Cuneus_R | DMN | 0.29 | 0.38 | -4.71 | -0.09 |
| R.Cereb | Cerebellum_R | Subcortical | R.LIPd | Angular_R | DAN | -0.12 | -0.06 | -5.12 | -0.09 |
| L.SFL | Supp_Motor_Area_L | DMN | L.POS2 | Precuneus_L | FPN | -0.08 | 0.00 | -5.44 | -0.09 |
| R.V6A | Occipital_Sup_R | Visual | L.9-46d | Frontal_Mid_L | VAN | -0.12 | -0.07 | -3.21 | -0.10 |
| R.23c | Cingulum_Mid_R | VAN | L.STSdp | Temporal_Mid_L | DMN | -0.09 | -0.03 | -4.05 | -0.10 |
| L.Pallidum | Pallidum_L | Subcortical | L.Thalamus | Thalamus_L | Subcortical | -0.02 | 0.05 | -5.07 | -0.10 |
| R.PGi | Angular_R | DMN | R.6v | Precentral_R | Somatomotor | -0.18 | -0.08 | -6.36 | -0.10 |
| L.HP | Hippocampus_L | Subcortical | B.Stem | - | Subcortical | 0.87 | 0.96 | -4.96 | -0.10 |
| R.RSC | Cingulum_Post_R | DMN | L.SFL | Supp_Motor_Area_L | DMN | -0.09 | -0.01 | -5.22 | -0.10 |
| R.A4 | Temporal_Sup_R | Somatomotor | L.A5 | Temporal_Mid_L | DMN | 0.37 | 0.51 | -6.16 | -0.10 |
| R.SFL | Supp_Motor_Area_R | DMN | L.POS1 | Precuneus_L | DMN | -0.18 | -0.08 | -6.57 | -0.10 |
| R.47s | Insula_R | DMN | L.a10p | Frontal_Sup_Orb_L | DMN | -0.01 | 0.08 | -4.94 | -0.10 |
| R.RSC | Cingulum_Post_R | DMN | L.10pp | Frontal_Sup_Orb_L | Limbic | 0.00 | 0.05 | -3.88 | -0.10 |
| R.pOFC | Olfactory_R | Limbic | L.TGv | Temporal_Inf_L | Limbic | 0.07 | 0.16 | -4.90 | -0.11 |
| R.HC | Hippocampus_R | Subcortical | R.46 | Frontal_Mid_R | FPN | 0.07 | 0.15 | -6.21 | -0.11 |

| | | | | | | | | | |
|---|---|---|---|---|---|---|---|---|---|
| L.VMV1 | Lingual_L | Visual | L.STV | Temporal_Sup_L | DMN | 0.01 | 0.09 | -5.18 | -0.12 |
| R.A5 | Temporal_Sup_R | Somatomotor | R.PBelt | Temporal_Sup_R | Somatomotor | 0.15 | 0.29 | -5.83 | -0.12 |
| R.Pir | Insula_R | Limbic | L.a10p | Frontal_Sup_Orb_L | DMN | -0.08 | -0.02 | -4.15 | -0.12 |
| L.LBelt | Temporal_Sup_L | Somatomotor | L.RI | Rolandic_Oper_L | Somatomotor | 0.57 | 0.65 | -3.96 | -0.13 |
| L.PFop | SupraMarginal_L | VAN | L.9m | Frontal_Sup_Medial_L | DMN | -0.30 | -0.19 | -5.77 | -0.13 |
| R.ProS | Lingual_R | Visual | L.TPOJ1 | Temporal_Mid_L | DMN | -0.02 | 0.06 | -4.96 | -0.15 |
| R.ProS | Lingual_R | Visual | L.TPOJ2 | Temporal_Mid_L | DAN | 0.00 | 0.07 | -4.70 | -0.15 |
| R.Amy | Hippocampus_R | Subcortical | L.IFSa | Frontal_Inf_Tri_L | FPN | 0.09 | 0.19 | -7.22 | -0.17 |
| R.25 | Olfactory_R | Limbic | R.24dd | Cingulum_Mid_R | Somatomotor | -0.11 | -0.04 | -4.77 | -0.17 |
| L.STSda | Temporal_Mid_L | DMN | L.MST | Occipital_Mid_L | Visual | -0.05 | 0.04 | -6.27 | -0.18 |
| R.PGi | Angular_R | DMN | R.SFL | Supp_Motor_Area_R | DMN | 0.16 | 0.24 | -4.44 | -0.18 |
| R.24dv | Cingulum_Mid_R | Somatomotor | L.PI | Temporal_Sup_L | VAN | 0.02 | 0.10 | -5.18 | -0.22 |
| R.p32 | Frontal_Sup_Medial_R | DMN | L.31a | Cingulum_Mid_L | DMN | 0.08 | 0.15 | -4.29 | -0.22 |
| R.s32 | Frontal_Med_Orb_R | DMN | R.PHA2 | ParaHippocampal_R | Visual | 0.11 | 0.16 | -3.52 | -0.22 |
| R.TE2p | Temporal_Inf_R | DAN | L.55b | Precentral_L | DAN | -0.12 | -0.05 | -4.40 | -0.22 |
| R.SFL | Supp_Motor_Area_R | DMN | L.IP1 | Occipital_Mid_L | DAN | -0.19 | -0.09 | -6.90 | -0.29 |
| R.Pallidum | Pallidum_R | Subcortical | R.Putamen | Putamen_R | Subcortical | 0.27 | 0.33 | -4.12 | -0.31 |
| R.pOFC | Olfactory_R | Limbic | R.10pp | Frontal_Sup_Orb_R | Limbic | 0.09 | 0.19 | -5.11 | -0.34 |
| R.s32 | Frontal_Med_Orb_R | DMN | R.H | Hippocampus_R | DMN | 0.12 | 0.20 | -4.76 | -0.81 |

**Abbreviations:** ASD: autism spectrum disorder, DAN: dorsal attention network, DMN: default mode network, FP: fronto-parietal, TDC: typically developing control, and VAN: ventral attention network.

**Table S10.** The list of discriminative FCs reproducible across the five datasets.

| ROI 1 | | | ROI 2 | | | Discovery dataset | | | Child cohort | | | Adolescent cohort | | | US adult cohort | | | Japanese adult cohort | | |
|---|---|---|---|---|---|---|---|---|---|---|---|---|---|---|---|---|---|---|---|---|
| **Glasser** | **AAL label** | **Network** | **Glasser** | **AAL label** | **Network** | ***z*(ASD)** | ***z*(TDC)** | ***t*-value** | ***z*(ASD)** | ***z*(TDC)** | ***t*-value** | ***z*(ASD)** | ***z*(TDC)** | ***t*-value** | ***z*(ASD)** | ***z*(TDC)** | ***t*-value** | ***z*(ASD)** | ***z*(TDC)** | ***t*-value** |
| R.Amy | Hippocampus_R | Subcortical | R.A5 | Temporal_Sup_R | Somatomotor | 0.12 | 0.01 | 8.03 | 0.15 | 0.11 | 1.84 | 0.10 | 0.06 | 1.26 | -0.02 | -0.06 | 1.33 | 0.10 | 0.00 | 2.77 |
| R.PBelt | Temporal_Sup_R | Somatomotor | L.POS2 | Precuneus_L | FPN | 0.03 | -0.08 | 7.42 | -0.03 | -0.09 | 2.88 | 0.01 | -0.06 | 3.49 | -0.02 | -0.06 | 1.50 | 0.08 | 0.01 | 2.00 |
| R.Amy | Hippocampus_R | Subcortical | R.6d | Precentral_R | Somatomotor | 0.07 | -0.04 | 7.20 | 0.05 | 0.05 | 0.32 | 0.07 | 0.05 | 0.85 | 0.06 | -0.09 | 4.65 | 0.13 | 0.01 | 3.21 |
| L.MidB | - | Subcortical | R.STSda | Temporal_Sup_R | DMN | 0.15 | 0.04 | 6.85 | 0.07 | 0.04 | 1.54 | 0.10 | 0.05 | 2.27 | 0.07 | 0.00 | 2.38 | 0.09 | 0.02 | 1.68 |
| R.MidB | - | Subcortical | R.PBelt | Temporal_Sup_R | Somatomotor | 0.17 | 0.06 | 6.65 | 0.13 | 0.12 | 0.66 | 0.09 | 0.06 | 1.50 | 0.06 | -0.02 | 2.44 | 0.14 | 0.07 | 1.60 |
| R.Putamen | Putamen_R | Subcortical | R.A5 | Temporal_Sup_R | Somatomotor | 0.05 | -0.03 | 6.36 | 0.04 | 0.03 | 0.81 | 0.05 | 0.02 | 1.36 | 0.00 | -0.06 | 1.91 | 0.04 | -0.01 | 1.70 |
| R.HC | Hippocampus_R | Subcortical | L.A5 | Temporal_Mid_L | DMN | 0.12 | 0.04 | 6.02 | 0.13 | 0.13 | 0.12 | 0.09 | 0.07 | 1.07 | 0.02 | -0.03 | 1.56 | 0.11 | 0.03 | 2.09 |
| R.Pallidum | Pallidum_R | Subcortical | L.MST | Occipital_Mid_L | Visual | 0.02 | -0.05 | 5.97 | 0.04 | 0.01 | 1.53 | 0.04 | 0.00 | 2.46 | 0.00 | -0.02 | 0.43 | 0.00 | -0.05 | 2.26 |

| | | | | | | | | | | | | | | | | | | | | |
|---|---|---|---|---|---|---|---|---|---|---|---|---|---|---|---|---|---|---|---|---|
| R.MidB | - | Subcortical | R.TGd | Temporal_Pole_Mid_R | Limbic | 0.03 | -0.06 | 5.70 | 0.00 | -0.05 | 2.34 | -0.02 | -0.05 | 1.25 | 0.03 | -0.05 | 2.60 | -0.05 | -0.08 | 0.82 |
| L.MidB | - | Subcortical | R.TPOJ1 | Temporal_Mid_R | DAN | 0.01 | -0.06 | 5.68 | 0.04 | -0.01 | 2.29 | 0.02 | -0.02 | 1.63 | -0.01 | -0.10 | 2.82 | 0.03 | -0.05 | 2.23 |
| L.V7 | Occipital_Mid_L | Visual | L.RSC | - | DMN | -0.04 | -0.12 | 5.23 | -0.01 | -0.06 | 2.57 | -0.03 | -0.05 | 1.31 | -0.04 | -0.07 | 1.02 | -0.09 | -0.15 | 1.88 |
| R.PFm | Parietal_Inf_R | DMN | R.d32 | Cingulum_Ant_R | DMN | 0.38 | 0.29 | 5.23 | 0.34 | 0.31 | 1.44 | 0.35 | 0.32 | 1.31 | 0.33 | 0.29 | 0.96 | 0.34 | 0.31 | 0.58 |
| R.MidB | - | Subcortical | R.TE1a | Temporal_Mid_R | DMN | -0.02 | -0.09 | 4.71 | -0.05 | -0.12 | 3.33 | -0.09 | -0.14 | 2.03 | -0.04 | -0.09 | 1.72 | -0.06 | -0.10 | 1.12 |
| R.SFL | Supp_Motor_Area_R | DMN | L.9-46d | Frontal_Mid_L | DAN | 0.14 | 0.05 | 4.71 | 0.13 | 0.09 | 1.88 | 0.12 | 0.03 | 3.66 | 0.10 | 0.06 | 0.85 | 0.12 | 0.07 | 0.73 |
| R.Amy | Hippocampus_R | Subcortical | R.TGd | Temporal_Pole_Mid_R | Limbic | -0.01 | -0.07 | 4.23 | -0.02 | -0.04 | 0.80 | 0.01 | -0.02 | 1.40 | 0.04 | -0.02 | 1.82 | -0.02 | -0.05 | 0.69 |
| R.7Pm | Precuneus_R | FPN | L.A1 | Rolandic_Oper_L | Somatomotor | -0.05 | -0.10 | 4.05 | -0.11 | -0.12 | 0.66 | -0.08 | -0.10 | 0.55 | -0.08 | -0.08 | 0.09 | -0.03 | -0.07 | 1.63 |
| R.PCV | Precuneus_R | FPN | L.AVI | Insula_L | FPN | 0.00 | -0.05 | 3.99 | -0.03 | -0.04 | 0.76 | -0.02 | -0.11 | 3.73 | -0.06 | -0.08 | 0.78 | 0.06 | -0.04 | 2.63 |
| R.25 | Olfactory_R | Limbic | R.IPS1 | Occipital_Sup_R | DAN | -0.04 | -0.09 | 3.94 | -0.05 | -0.11 | 3.00 | -0.07 | -0.08 | 0.19 | -0.01 | -0.03 | 0.62 | -0.06 | -0.09 | 1.09 |
| R.25 | Olfactory_R | Limbic | R.PHT | Temporal_Inf_R | DAN | 0.01 | -0.04 | 3.90 | -0.02 | -0.06 | 1.76 | -0.03 | -0.04 | 0.07 | -0.02 | -0.05 | 1.04 | -0.05 | -0.11 | 1.46 |

| | | | | | | | | | | | | | | | | | | | | |
|---|---|---|---|---|---|---|---|---|---|---|---|---|---|---|---|---|---|---|---|---|
| L.Pallidum | Pallidum_L | Subcortical | L.TE2a | Temporal_Inf_L | Limbic | 0.00 | -0.06 | 3.34 | -0.03 | -0.06 | 1.38 | 0.02 | 0.01 | 0.50 | 0.03 | 0.00 | 0.87 | -0.03 | -0.04 | 0.54 |
| R.s6-8 | Frontal_Sup_R | FPN | L.a9-46v | Frontal_Mid_L | FPN | 0.09 | 0.04 | 2.40 | 0.03 | 0.02 | 0.48 | 0.07 | 0.02 | 2.02 | 0.11 | 0.05 | 1.72 | 0.10 | 0.06 | 0.75 |
| L.p47r | Frontal_Inf_Tri_L | FPN | L.10d | Frontal_Sup_Medial_L | DMN | 0.10 | 0.05 | 2.16 | 0.15 | 0.06 | 2.99 | 0.17 | 0.06 | 3.24 | 0.05 | -0.03 | 1.56 | 0.04 | 0.02 | 0.21 |
| R.RSC | Cingulum_Post_R | DMN | L.RSC | - | DMN | 1.03 | 1.08 | -2.94 | 1.02 | 1.08 | -2.14 | 1.03 | 1.09 | -2.05 | 0.92 | 0.96 | -0.94 | 1.06 | 1.11 | -1.08 |
| L.LBelt | Temporal_Sup_L | Somatomotor | L.Ig | Insula_L | Somatomotor | 0.24 | 0.30 | -3.88 | 0.27 | 0.33 | -2.59 | 0.31 | 0.39 | -2.98 | 0.26 | 0.28 | -0.51 | 0.40 | 0.41 | -0.22 |
| R.STSdp | Temporal_Sup_R | DMN | L.V4t | Occipital_Mid_L | Visual | -0.04 | 0.01 | -3.92 | 0.02 | 0.05 | -1.52 | -0.02 | 0.05 | -3.01 | 0.00 | 0.03 | -1.04 | -0.04 | 0.04 | -2.08 |
| R.47l | Frontal_Inf_Orb_R | DMN | L.47l | Frontal_Inf_Orb_L | DMN | 0.48 | 0.56 | -4.05 | 0.53 | 0.55 | -0.71 | 0.52 | 0.59 | -2.56 | 0.54 | 0.58 | -1.06 | 0.61 | 0.62 | -0.01 |
| R.23c | Cingulum_Mid_R | VAN | L.STSdp | Temporal_Mid_L | DMN | -0.09 | -0.03 | -4.05 | -0.05 | -0.02 | -1.55 | -0.10 | -0.03 | -3.38 | -0.07 | -0.02 | -1.40 | -0.16 | -0.05 | -2.46 |
| R.FFC | Fusiform_R | Visual | L.55b | Precentral_L | DAN | -0.07 | 0.00 | -4.34 | 0.03 | 0.05 | -0.58 | 0.00 | 0.06 | -2.26 | 0.03 | 0.03 | -0.09 | 0.01 | 0.03 | -0.68 |
| R.ProS | Lingual_R | Visual | L.TPOJ2 | Temporal_Mid_L | DAN | 0.00 | 0.07 | -4.70 | 0.00 | 0.04 | -1.70 | 0.01 | 0.03 | -0.68 | -0.02 | 0.03 | -1.41 | 0.04 | 0.07 | -0.82 |

| | | | | | | | | | | | | | | | | | | | | |
|---|---|---|---|---|---|---|---|---|---|---|---|---|---|---|---|---|---|---|---|---|
| L.HP | Hippocampus_L | Subcortical | B.Stem | - | Subcortical | 0.87 | 0.96 | -4.96 | 0.82 | 0.86 | -1.82 | 0.91 | 0.94 | -1.46 | 0.82 | 0.84 | -0.47 | 1.06 | 1.13 | -1.55 |
| R.ProS | Lingual_R | Visual | L.TPOJ1 | Temporal_Mid_L | DMN | -0.02 | 0.06 | -4.96 | -0.01 | 0.03 | -1.67 | -0.05 | 0.02 | -2.72 | -0.02 | 0.04 | -1.77 | -0.03 | 0.04 | -2.26 |
| R.24dv | Cingulum_Mid_R | Somatomotor | L.PI | Temporal_Sup_L | DAN | 0.02 | 0.10 | -5.18 | 0.06 | 0.10 | -2.63 | 0.08 | 0.08 | -0.20 | 0.05 | 0.08 | -1.01 | 0.15 | 0.19 | -0.82 |
| R.MidB | - | Subcortical | B.Stem | - | Subcortical | 0.23 | 0.31 | -5.40 | 0.25 | 0.30 | -2.30 | 0.22 | 0.27 | -1.95 | 0.23 | 0.26 | -0.81 | 0.22 | 0.33 | -3.07 |
| R.9m | Frontal_Sup_Medial_R | DMN | R.PCV | Precuneus_R | FPN | 0.00 | 0.11 | -5.49 | 0.07 | 0.09 | -1.06 | 0.04 | 0.07 | -1.20 | -0.02 | 0.05 | -1.53 | 0.05 | 0.12 | -1.13 |
| R.TPOJ1 | Temporal_Mid_R | VAN | R.4 | Precentral_R | Somatomotor | 0.08 | 0.17 | -5.65 | 0.07 | 0.10 | -1.47 | 0.10 | 0.12 | -0.75 | 0.11 | 0.13 | -0.59 | 0.13 | 0.16 | -0.72 |
| R.TPOJ1 | Temporal_Mid_R | VAN | L.MT | Occipital_Mid_L | Visual | 0.05 | 0.16 | -5.72 | 0.10 | 0.12 | -0.81 | 0.09 | 0.13 | -1.66 | 0.10 | 0.14 | -1.07 | 0.08 | 0.15 | -1.29 |
| R.A5 | Temporal_Sup_R | Somatomotor | R.PBelt | Temporal_Sup_R | Somatomotor | 0.15 | 0.29 | -5.83 | 0.16 | 0.17 | -0.72 | 0.16 | 0.22 | -2.05 | 0.15 | 0.22 | -2.03 | 0.20 | 0.35 | -2.26 |
| R.MidB | - | Subcortical | R.46 | Frontal_Mid_R | FPN | -0.01 | 0.07 | -6.02 | 0.00 | 0.03 | -1.67 | -0.01 | 0.06 | -3.49 | -0.02 | 0.01 | -0.94 | -0.01 | 0.03 | -1.16 |
| R.MidB | - | Subcortical | L.a9-46v | Frontal_Mid_L | FPN | 0.01 | 0.10 | -6.07 | 0.04 | 0.05 | -0.37 | 0.03 | 0.08 | -2.43 | 0.03 | 0.09 | -2.10 | 0.03 | 0.08 | -1.30 |

| | | | | | | | | | | | | | | | | | | | | |
|---|---|---|---|---|---|---|---|---|---|---|---|---|---|---|---|---|---|---|---|---|
| R.A4 | Temporal_Sup_R | Somatomotor | L.A5 | Temporal_Mid_L | DMN | 0.37 | 0.51 | -6.16 | 0.39 | 0.44 | -1.88 | 0.34 | 0.47 | -4.56 | 0.35 | 0.44 | -2.08 | 0.67 | 0.70 | -0.35 |
| R.HC | Hippocampus_R | Subcortical | R.46 | Frontal_Mid_R | FPN | 0.07 | 0.15 | -6.21 | 0.06 | 0.08 | -1.33 | 0.07 | 0.10 | -1.53 | 0.03 | 0.08 | -1.72 | 0.07 | 0.12 | -1.14 |
| L.Putamen | Putamen_L | Subcortical | R.A4 | Temporal_Sup_R | Somatomotor | -0.02 | 0.06 | -6.26 | 0.06 | 0.06 | -0.31 | 0.05 | 0.07 | -1.06 | 0.03 | 0.12 | -2.81 | 0.09 | 0.10 | -0.43 |

**Abbreviations:** ASD: autism spectrum disorder, FP: fronto-parietal, DAN: dorsal attention network, DMN: default mode network, HP: hippocampus, MOG: middle occpital gyrus, MTG: middle temporal gyrus, MFG: middle frontal gyrus, SFG: superior frontal gyrus, FUS: fusiform gyrus, IFG: inferior frontal gyrus, Precent: Precentral gyrus, ITG: inferior temporal gyrus, IPL: inferior parietal lobule, ROI: region of interest, STG: superior temporal gyrus, TDC: typically developing control, VAN: ventral attention network

**Table S11**. Demographic information of SCZ and MDD datasets.

| | **SCZ** | | | **MDD** | | |
|---|---|---|---|---|---|---|
| | **N (M/F)** | **Age** | **Mean FD [mm]** | **N (M/F)** | **Age** | **Mean FD [mm]** |
| COI | - | - | - | 52 (23/29) | 45.23 (12.64) | 0.16 (0.06) |
| KUT | 42/ (20/22) | 41.90 (10.43) | 0.14 (0.06) | 15 (9/6) | 43.93 (11.60) | 0.10 (0.04) |
| SWA1 | 18 (14/4) | 42.83 (8.65) | 0.19 (0.06) | - | - | - |
| UTO1 | 34 (24/10) | 30.91 (10.42) | 0.11 (0.03) | 59 (35/24) | 38.19 (11.44) | 0.10 (0.04) |
| UTO2 | - | - | - | - | - | - |
| **Total** | 94 (58/36) | 38.11 (11.40) | 0.14 (0.06) | 126 (67/59) | 41.78 (12.35) | 0.13 (0.06) |

**Abbreviations:** ASD: autism spectrum disorder, COI: Center of Innovation, F: female, KUT: Kyoto University TimTrio, M: male, MDD: major depressive disorder, SCZ: schizophrenia, SD: standard deviation, SWA: Showa University, TDC: typically developing control, and UTO: University of Tokyo.

**Table S12**. The classification performance of the SCZ and MDD neuromarkers.

| | AUC | Accuracy [%] | Sensitivity [%] | Specificity [%] | MCC |
|---|---|---|---|---|---|
| **Classification performance of SCZ neuromarker** | | | | | |
| **All** | **0.89** | **81.99** | **82.98** | **81.82** | **0.51** |
| COI | - | 65.09 | - | 65.09 | - |
| KUT | 0.91 | 84.74 | 83.33 | 85.14 | 0.62 |
| SWA1 | 0.96 | 85.94 | 88.89 | 85.63 | 0.53 |
| UTO1 | 0.82 | 82.50 | 79.41 | 83.72 | 0.60 |
| UTO2 | - | 94.44 | - | 94.44 | - |
| **Classification performance of MDD neuromarker** | | | | | |
| **All** | **0.78** | **67.60** | **74.60** | **66.00** | **0.32** |
| COI | 0.65 | 56.96 | 71.15 | 50.00 | 0.20 |
| KUT | 0.85 | 73.01 | 80.00 | 72.30 | 0.32 |
| SWA1 | - | 69.54 | - | 69.54 | - |
| UTO1 | 0.75 | 68.28 | 76.27 | 62.79 | 0.38 |
| UTO2 | - | 77.78 | - | 77.78 | - |

**Abbreviations:** AUC: area under the curve, COI: Center of Innovation, KUT: Kyoto University TimTrio, MCC: Matthews correlation coefficient, SWA: Showa University, UTO: University of Tokyo,

**Table S13.** The list of discriminative FCs for SCZ.

| ROI 1 | | | ROI 2 | | | *z*(SCZ) | *z*(TDC) | *t*-value | Mean weight |
|---|---|---|---|---|---|---|---|---|---|
| Glasser's label | AAL label | Network | Glasser's label | AAL label | Network | | | | |
| L.MidB | - | Subcortical | L.V2 | Lingual_L | Visual | 0.00 | -0.12 | 6.90 | 0.20 |
| L.MidB | - | Subcortical | L.ProS | Precuneus_L | Visual | 0.11 | -0.01 | 6.29 | 0.17 |
| L.23c | Cingulum_Mid_L | VAN | L.7Pm | Precuneus_L | Frontoparietal | 0.34 | 0.21 | 5.04 | 0.13 |
| R.PeEc | ParaHippocampal_R | Limbic | L.ProS | Precuneus_L | Visual | 0.04 | -0.03 | 4.68 | 0.12 |
| L.HP | Hippocampus_L | Subcortical | L.3b | Postcentral_L | Somatomotor | 0.01 | -0.13 | 6.62 | 0.11 |
| R.Amy | Hippocampus_R | Subcortical | L.31pv | Cingulum_Post_L | DMN | 0.03 | -0.09 | 7.73 | 0.11 |
| L.PoI1 | Insula_L | VAN | L.6a | Frontal_Sup_L | DAN | 0.16 | 0.05 | 5.69 | 0.10 |
| R.AAIC | Insula_R | VAN | R.9p | Frontal_Sup_R | DMN | 0.15 | 0.07 | 4.44 | 0.10 |
| R.PSL | Temporal_Sup_R | VAN | L.9-46d | Frontal_Mid_L | VAN | 0.22 | 0.10 | 4.99 | 0.09 |
| R.Amy | Hippocampus_R | Subcortical | L.7m | Precuneus_L | DMN | 0.00 | -0.13 | 8.16 | 0.09 |
| R.2 | Postcentral_R | DAN | L.PF | SupraMarginal_L | VAN | 0.24 | 0.04 | 6.91 | 0.08 |
| R.MBelt | Heschl_R | Somatomotor | R.MIP | Occipital_Sup_R | DAN | -0.04 | -0.11 | 4.44 | 0.08 |
| L.MidB | - | Subcortical | L.VMV1 | Lingual_L | Visual | 0.11 | -0.02 | 6.63 | 0.07 |
| L.NAcc | Caudate_L | Subcortical | L.V8 | Fusiform_L | Visual | 0.04 | -0.11 | 6.77 | 0.06 |
| R.VMV3 | Lingual_R | Visual | R.V3A | Occipital_Sup_R | Visual | 0.34 | 0.48 | -5.42 | -0.05 |
| R.Ig | Insula_R | Somatomotor | L.Ig | Insula_L | Somatomotor | 0.45 | 0.66 | -7.75 | -0.05 |
| R.FOP2 | Rolandic_Oper_R | Somatomotor | R.3a | Precentral_R | Somatomotor | 0.19 | 0.31 | -5.24 | -0.07 |
| R.43 | Rolandic_Oper_R | Somatomotor | R.13l | Frontal_Inf_Orb_R | Limbic | -0.14 | -0.05 | -5.00 | -0.07 |

| | | | | | | | | | |
|---|---|---|---|---|---|---|---|---|---|
| L.3a | Postcentral_L | Somatomotor | L.4 | Precentral_L | Somatomotor | 0.76 | 1.03 | -8.14 | -0.07 |
| R.13l | Frontal_Inf_Orb_R | Limbic | R.24dv | Cingulum_Mid_R | Somatomotor | -0.13 | -0.05 | -5.00 | -0.08 |
| R.4 | Precentral_R | Somatomotor | L.3a | Postcentral_L | Somatomotor | 0.68 | 0.92 | -7.40 | -0.08 |
| R.47s | Insula_R | DMN | L.23c | Cingulum_Mid_L | VAN | -0.26 | -0.17 | -4.82 | -0.09 |
| R.STSdp | Temporal_Sup_R | DMN | L.FFC | Fusiform_L | Visual | -0.10 | 0.01 | -5.56 | -0.09 |
| L.Thalamus | Thalamus_L | Subcortical | L.p24 | Cingulum_Ant_L | DMN | 0.07 | 0.17 | -5.37 | -0.10 |
| R.4 | Precentral_R | Somatomotor | L.Ig | Insula_L | Somatomotor | 0.31 | 0.48 | -6.84 | -0.10 |
| R.3a | Precentral_R | Somatomotor | L.4 | Precentral_L | Somatomotor | 0.60 | 0.87 | -8.57 | -0.12 |
| R.HC | Hippocampus_R | Subcortical | R.FOP4 | Insula_R | VAN | 0.14 | 0.33 | -10.30 | -0.13 |
| L.FOP3 | Insula_L | VAN | L.23d | Cingulum_Mid_L | DMN | -0.17 | -0.05 | -6.33 | -0.15 |
| R.HC | Hippocampus_R | Subcortical | L.FOP3 | Insula_L | VAN | 0.10 | 0.25 | -9.41 | -0.15 |
| R.Amy | Hippocampus_R | Subcortical | R.FOP5 | Insula_R | VAN | 0.07 | 0.23 | -9.40 | -0.15 |
| R.3a | Precentral_R | Somatomotor | R.4 | Precentral_R | Somatomotor | 0.72 | 1.00 | -8.44 | -0.16 |
| R.HC | Hippocampus_R | Subcortical | L.p32pr | Cingulum_Mid_L | VAN | 0.10 | 0.26 | -9.32 | -0.16 |
| L.FOP3 | Insula_L | VAN | L.RSC | - | DMN | -0.13 | -0.02 | -6.08 | -0.18 |
| R.24dv | Cingulum_Mid_R | Somatomotor | L.OP2-3 | Rolandic_Oper_L | Somatomotor | 0.29 | 0.43 | -6.81 | -0.19 |
| R.HC | Hippocampus_R | Subcortical | R.p32pr | Cingulum_Mid_R | VAN | 0.09 | 0.26 | -9.80 | -0.21 |
| R.HC | Hippocampus_R | Subcortical | L.FOP4 | Insula_L | VAN | 0.10 | 0.28 | -10.66 | -0.24 |
| R.HC | Hippocampus_R | Subcortical | L.SCEF | Supp_Motor_Area_L | VAN | 0.06 | 0.22 | -9.07 | -0.27 |
| R.Amy | Hippocampus_R | Subcortical | L.SCEF | Supp_Motor_Area_L | VAN | 0.06 | 0.24 | -9.60 | -0.35 |
| R.Amy | Hippocampus_R | Subcortical | R.p32pr | Cingulum_Mid_R | VAN | 0.08 | 0.26 | -10.45 | -0.42 |
| R.Amy | Hippocampus_R | Subcortical | L.FOP1 | Rolandic_Oper_L | VAN | 0.13 | 0.30 | -9.74 | -0.44 |

| | | | | | | | | | |
|---|---|---|---|---|---|---|---|---|---|
| R.Amy | Hippocampus_R | Subcortical | L.a24pr | Cingulum_Mid_L | VAN | 0.08 | 0.27 | -10.41 | -1.31 |

**Abbreviations:** AAL: automated anatomical labelling, DAN: dorsal attention network, DMN: default mode network, FP: fronto-parietal, ROI: region of interest, SCZ: schizophrenia, TDC: typically developing control, and VAN: ventral attention network.

**Table S14. The list of discriminative FCs for MDD.**

| ROI 1 | | | ROI 2 | | | | | | |
|---|---|---|---|---|---|---|---|---|---|
| Glasser's label | AAL label | Network | Glasser's label | AAL label | Network | *z*(MDD) | *z*(TDC) | *t*-value | Mean weight |
| L.Amy | Amygdala_L | Subcortical | R.VMV1 | Lingual_R | Visual | -0.03 | -0.16 | 6.69 | 6.69 |
| R.MidB | - | Subcortical | L.3b | Postcentral_L | Somatomotor | 0.07 | -0.09 | 6.68 | 6.68 |
| L.VMV2 | Lingual_L | Visual | L.PF | SupraMarginal_L | VAN | 0.13 | 0.02 | 5.89 | 5.89 |
| R.a9-46v | Frontal_Mid_R | FPN | L.3b | Postcentral_L | Somatomotor | -0.02 | -0.11 | 4.98 | 4.98 |
| R.V1 | Calcarine_R | Visual | L.6ma | Frontal_Sup_L | VAN | 0.06 | -0.02 | 4.81 | 4.81 |
| L.A4 | Temporal_Sup_L | Somatomotor | L.23c | Cingulum_Mid_L | VAN | 0.20 | 0.11 | 4.75 | 4.75 |
| R.7Pm | Precuneus_R | FPN | L.TGd | Temporal_Pole_Mid_L | Limbic | -0.09 | -0.16 | 4.52 | 4.52 |
| R.PreS | Hippocampus_R | Visual | L.POS1 | Precuneus_L | DMN | 0.37 | 0.29 | 4.02 | 4.02 |
| L.11l | Frontal_Sup_Orb_L | FPN | L.LO2 | Occipital_Inf_L | Visual | -0.02 | -0.08 | 3.87 | 3.87 |
| R.13l | Frontal_Inf_Orb_R | Limbic | R.47m | Frontal_Inf_Orb_R | DMN | 0.26 | 0.36 | -4.03 | -4.03 |
| R.AVI | Insula_R | FPN | L.ProS | Precuneus_L | Visual | -0.05 | 0.01 | -4.07 | -4.07 |
| R.47m | Frontal_Inf_Orb_R | DMN | L.TA2 | Temporal_Sup_L | Somatomotor | 0.01 | 0.08 | -4.34 | -4.34 |
| R.47m | Frontal_Inf_Orb_R | DMN | L.MBelt | Temporal_Sup_L | Somatomotor | 0.00 | 0.07 | -4.47 | -4.47 |
| R.47m | Frontal_Inf_Orb_R | DMN | L.A1 | Rolandic_Oper_L | Somatomotor | -0.01 | 0.06 | -4.50 | -4.50 |
| R.PHA1 | Fusiform_R | Visual | R.PBelt | Temporal_Sup_R | Somatomotor | -0.04 | 0.03 | -4.57 | -4.57 |
| L.TGv | Temporal_Inf_L | Limbic | L.TE2a | Temporal_Inf_L | Limbic | 0.16 | 0.28 | -5.10 | -5.10 |

| | | | | | | | | | |
|---|---|---|---|---|---|---|---|---|---|
| R.A5 | Temporal_Sup_R | Somatomotor | R.OP2-3 | Rolandic_Oper_R | Somatomotor | 0.11 | 0.21 | -5.24 | -5.24 |
| R.Thalamus | Thalamus_R | Subcortical | R.p32pr | Cingulum_Mid_R | VAN | -0.19 | -0.10 | -5.37 | -5.37 |
| L.Ig | Insula_L | Somatomotor | L.POS1 | Precuneus_L | DMN | 0.04 | 0.13 | -5.38 | -5.38 |
| R.v23ab | Precuneus_R | DMN | L.Ig | Insula_L | Somatomotor | -0.03 | 0.06 | -5.61 | -5.61 |
| R.PreS | Hippocampus_R | Visual | R.p32pr | Cingulum_Mid_R | VAN | -0.10 | -0.01 | -5.69 | -5.69 |
| R.6mp | Supp_Motor_Area_R | Somatomotor | R.5L | Postcentral_R | DAN | 0.41 | 0.54 | -5.80 | -5.80 |
| R.a24 | Cingulum_Ant_R | DMN | L.52 | Temporal_Sup_L | VAN | -0.03 | 0.07 | -5.83 | -5.83 |
| R.LO1 | Occipital_Mid_R | Visual | L.LO1 | Occipital_Mid_L | Visual | 0.57 | 0.72 | -5.83 | -5.83 |
| L.52 | Temporal_Sup_L | VAN | L.a24 | Cingulum_Ant_L | DMN | 0.00 | 0.10 | -5.94 | -5.94 |
| R.A4 | Temporal_Sup_R | Somatomotor | L.5m | Precuneus_L | Somatomotor | 0.04 | 0.15 | -5.94 | -5.94 |
| R.Pallidum | Pallidum_R | Subcortical | R.Putamen | Putamen_R | Subcortical | 0.23 | 0.33 | -6.01 | -6.01 |
| R.OP4 | Rolandic_Oper_R | Somatomotor | L.5m | Precuneus_L | Somatomotor | 0.11 | 0.22 | -6.30 | -6.30 |
| R.OP1 | Rolandic_Oper_R | Somatomotor | L.5m | Precuneus_L | Somatomotor | 0.20 | 0.33 | -6.38 | -6.38 |
| R.5L | Postcentral_R | DAN | L.5L | Precuneus_L | Somatomotor | 0.71 | 0.87 | -6.40 | -6.40 |
| R.2 | Postcentral_R | DAN | L.5m | Precuneus_L | Somatomotor | 0.32 | 0.47 | -6.41 | -6.41 |
| R.A5 | Temporal_Sup_R | Somatomotor | L.A4 | Temporal_Sup_L | Somatomotor | 0.20 | 0.37 | -6.80 | -6.80 |
| R.A4 | Temporal_Sup_R | Somatomotor | R.4 | Precentral_R | Somatomotor | 0.13 | 0.27 | -6.81 | -6.81 |
| R.A4 | Temporal_Sup_R | Somatomotor | R.OP2-3 | Rolandic_Oper_R | Somatomotor | 0.17 | 0.31 | -6.87 | -6.87 |
| R.OP1 | Rolandic_Oper_R | Somatomotor | L.OP2-3 | Rolandic_Oper_L | Somatomotor | 0.40 | 0.55 | -6.90 | -6.90 |

| | | | | | | | | | |
|---|---|---|---|---|---|---|---|---|---|
| L.Ig | Insula_L | Somatomotor | L.3a | Postcentral_L | Somatomotor | 0.44 | 0.61 | -7.76 | -7.76 |
| L.Ig | Insula_L | Somatomotor | L.RI | Rolandic_Oper_L | Somatomotor | 0.42 | 0.58 | -7.86 | -7.86 |
| R.Ig | Insula_R | Somatomotor | L.Ig | Insula_L | Somatomotor | 0.46 | 0.66 | -8.25 | -8.25 |

**Abbreviations:** AAL: automated anatomical labelling, DAN: dorsal attention network, DMN: default mode network, FP: fronto-parietal, ROI: region of interest, MDD: major depressive disorder, TDC: typically developing control, and VAN: ventral attention network.

**Table S15.** Imaging scanning protocols for R-fMRI data in the discovery dataset.

| Imaging site | Center of Innovation in Hiroshima University | Kyoto University | Showa University | University of Tokyo | University of Tokyo |
|---|---|---|---|---|---|
| Abbreviation | COI | KUT | SWA1 | UTO1 | UTO2 |
| Data resource | SRPBS | SRPBS | SRPBS | SRPBS | - |
| Data category | | | Discovery | | |
| MRI scanner | *Siemens Verio* | *Siemens TimTrio* | *Siemens Verio* | *GE MR750w* | *Philips Achieva* |
| Magnetic field strength | | | 3.0 T | | |
| Head coil | 12 | 32 | 12 | 24 | 8 |
| Field-of-view [mm] | | 212 × 212 | | | 224 × 224, 220 × 220 |
| Matrix size | | 64 × 64 | | | 64 × 64, 80 × 80 |
| in-plane resolution [mm] | | 3.3125 × 3.3125 | | | 3.5 × 3.5, 2.75 × 2.75 |
| Slice thickness [mm] | | 3.2 | | | 3.5, 5.0 |
| Slice gap [mm] | | 0.8 | | | 0.0 |
| Number of slices | | 40 | | | 45, 34 |
| Multi-band factor | | | - | | |
| Number of volumes | | 240 | | | 200 |
| Number of runs | | | 1 | | |
| Repetition time [ms] | | | 2,500 | | |
| Echo time [ms] | | | 30 | | |
| Flip angle [degree] | | 80 | | | 75 |
| Slice acquisition order | | | Ascending | | |
| Phase encoding | AP | | PA | | |
| Eye-status | | Fixate | | | Mixed |

**Table S16.** Image scanning protocols for R-fMRI data in the validation datasets.

| Imaging site | Showa University | University of Leuven | New York University Langone Medical Center | University of Utah, School of Medicine | Georgetown University | Oregon Health and Science University | Kennedy Krieger Institute | University of California, Los Angeles | San Diego State University | Trinity Centre for Health Sciences | University of Michigan | Yale School of Medicine |
|---|---|---|---|---|---|---|---|---|---|---|---|---|
| Abbreviation | SWA2 | Leuven | NYU | USM | GU | OHSU | KKI | UCLA | SDSU | TRINITY | UM | YALE |
| Data resource | BMB | ABIDE-I | ADBIE-I/ABIDE-II | ABIDE-I/ABIDE-II | ABIDE-II | ABIDE-II | ABIDE-I/ABIDE-II | ABIDE-I/ABIDE-II | ABIDE-II | ABIDE-I | ABIDE-I | ABIDE-I |
| Data category | Japanese Adult | ABIDE Adult | ABIDE Adult/Child/Adolescent | ABIDEAdult/Adolescent | Child | Child/Adolescent | Child | Child/Adolescent | Adolescent | Adolescent | Adolescent | Adolescent |
| MRI scanner | *Siemens Skyra fit* | *Philips Intera* | *Siemens Allegra* | *Siemens TrioTim* | *Siemens TrioTim* | *Siemens TrioTim* | *Philips Achieva* | *Siemens TrioTim* | *GE MR750* | *Philips Achieva* | *GE Signa* | *Siemens TimTrio* |
| Magnetic field strength | 3.0 T | 3.0 T | | | | | | | | | | |
| Head coil | 32 | 8 | NA | NA | 12 | 12 | 8 | NA | 8 | 8 | NA | NA |
| Field-of-view [mm] | 206 × 206 | 230 × 230 | 240 × 192 | 220 × 220 | 192 × 192 | 240 × 240 | 256 × 256 | 192 × 192 | 220 × 220 | 240 × 240 | 220 × 220 | 220 × 220 |
| Matrix size | 86 × 86 | 64 × 64 | 80 × 64 | 64 × 64 | 64 × 64 | 64 × 64 | 84 × 84 | 64 × 64 | 64 × 64 | 80 × 80 | 64 × 64 | 64 × 64 |
| in-plane resoluti | 2.4 × 2.4 | 3.59 × 3.59 | 3.0 × 3.0 | 3.4 × 3.4 | 3.0 × 3.0 | 3.8 × 3.8 | 3.0 × 3.0 | 3.0 × 3.0 | 3.4 × 3.4 | 3.0 × 3.0 | 3.4 × 3.4 | 3.4 × 3.4 |

| | | | | | | | | | | | | |
|---|---|---|---|---|---|---|---|---|---|---|---|---|
| on [mm] | | | | | | | | | | | | |
| Slice thickness [mm] | 2.4 | 4.0 | 4.0 | 3.0 | 2.5 | 3.8 | 3.0 | 4.0 | 3.4 | 3.5 | 3.0 | 4.0 |
| Slice gap [mm] | 0.0 | 0.0 | 0.0 | 0.3 | 0.5 | 0.0 | 0.0 | 0.0 | 0.0 | 0.35 | 0.0 | 0.0 |
| Number of slices | 60 | 32 | 33 | 40 | 43 | 36 | 47 | 34 | 42 | 38 | 40 | 34 |
| Multi-band factor | 6 | - | | | | | | | | | | |
| Number of volumes | 375 | 250 | 180 | 240 | 154 | 120 | 156 | 120 | 180 | 150 | 300 | 200 |
| Number of runs | AP: 2, PA: 2 | 1 | 1 | 1 | 1 | 3 | 1 | 1 | 1 | 1 | 1 | 1 |
| Repetition time [ms] | 800 | 1,667 | 2,000 | 2,000 | 2,000 | 2,500 | 2,500 | 3,000 | 2,000 | 2,000 | 2,000 | 2,000 |
| Echo time [ms] | 34.4 | 33 | 15 | 28 | 30 | 30 | 30 | 28 | 30 | 28 | 30 | 25 |
| Flip angle [degree] | 52 | 90 | 90 | 90 | 90 | 90 | 75 | 90 | 90 | 98 | 90 | 60 |
| Slice acquisition order | Interleaved | Ascending | Interleaved | Interleaved | Interleaved | Interleaved | Ascending | Interleaved | Interleaved | Ascending | Interleaved | Interleaved |

| Phase encodin g | AP, PA | AP | RL | AP | AP | AP | AP | AP | RL | AP | AP | AP |
|---|---|---|---|---|---|---|---|---|---|---|---|---|
| Eye-status | Fixate | Fixate | Mixed | Eye-open | Eye-open | Fixate | Fixate | Fixate | Fixate | Eye-closed | Fixate | Eye-open |
